# Replication-dependent histone (Repli-Histo) labeling dissects the physical properties of euchromatin/heterochromatin in living human cells

**DOI:** 10.1101/2024.10.20.618801

**Authors:** Katsuhiko Minami, Satoru Ide, Kako Nakazato, Kazunari Kaizu, Koichi Higashi, Sachiko Tamura, Atsushi Toyoda, Koichi Takahashi, Ken Kurokawa, Kazuhiro Maeshima

**Affiliations:** Genome Dynamics Laboratory, National Institute of Genetics, Mishima, Shizuoka 411-8540, Japan; Graduate Institute for Advanced Studies, SOKENDAI, Mishima, Shizuoka 411-8540, Japan; Laboratory for Biologically Inspired Computing, RIKEN Center for Biosystems Dynamics Research, Kobe, Hyogo 650-0047, Japan; Cell Modeling and Simulation Group, The Exploratory Research Center on Life and Living Systems, National Institutes of Natural Sciences, Okazaki, Aichi 444-8787, Japan; Genome Evolution Laboratory, National Institute of Genetics, Mishima, Shizuoka 411-8540, Japan; Comparative Genomics Laboratory, National Institute of Genetics, Mishima, Shizuoka 411-8540, Japan

**Author notes:** Research Center for Genome & Medical Sciences, Tokyo Metropolitan Institute of Medical Science, Tokyo 156-8506, Japan (Satoru Ide). Correspondence, National Institute of Genetics, Mishima, Shizuoka 411-8540, Japan.

## Abstract

A string of nucleosomes, where genomic DNA is wrapped around histones, is organized in the cell as chromatin. Chromatin in the cell varies greatly, from euchromatin to heterochromatin, in its genome functions. It is important to understand how heterochromatin is physically different from euchromatin. However, their specific labeling methods in living cells are limited. To address this, we have developed replication-dependent histone labeling (Repli-Histo labeling) to label nucleosomes in euchromatin and heterochromatin based on DNA replication timing. We investigated local nucleosome motion in the four chromatin classes from euchromatin to heterochromatin of living human and mouse cells. We found that more euchromatic regions (earlier replicated regions) show greater nucleosome motion. Notably, the motion profile in each chromatin class persists throughout interphase. Genome chromatin is essentially replicated from regions with greater nucleosome motions, even though the replication timing program is perturbed. Our findings, combined with computational modeling, suggest that earlier replicated regions have more accessibility and local chromatin motion can be a major determinant of genome-wide replication timing.

## Introduction

DNA is wrapped around core histones and forms a nucleosome ^1-4^. How are nucleosomes, together with other non-histone proteins/RNAs, organized in the cell as chromatin, and how does chromatin behave in living cells ^5, 6^? A wide range of imaging evidence has demonstrated that a string of nucleosomes is rather irregularly folded in the condensed chromatin domains in higher eukaryotic cells ^7-15^. Genome-wide genomic analyses such as Hi-C ^16^ also revealed chromatin domains with distinct epigenetic marks ^17-20^.

In order to conduct genome functions, chromatin in the cell is highly variable, ranging from euchromatin to heterochromatin ^15, 21-25^. A typical textbook model describes euchromatin as open and heterochromatin as closed and condensed ^26, 27^. However, recent studies suggest that euchromatin domains are not fully open, but rather condensed, except for enhancers and active transcription start sites ^10, 12, 13, 15, 28^. Given that both euchromatin and heterochromatin form condensed domains, how is heterochromatin physically different from euchromatin? In addition to their histone modifications (e.g., active and inactive ones) and non-histone components (e.g., HP1) ^21-25^, their dynamics might also be different ^29-33^.

Various chromatin labeling and microscopy systems for live-cell imaging and quantitative analyses have been developed to investigate chromatin dynamics and have contributed to understanding chromatin organization and function in live cells ^34 8, 10, 12, 29, 31-33, 35-46^. Among them, single-nucleosome imaging sensitively and accurately measures local chromatin dynamics in a whole nucleus ^33, 47-52^, or mitotic chromosomes ^33, 53^, and provides structural information on how chromatin organizes in living cells. Recently, single-nucleosome imaging revealed local nucleosome motion (on average) is almost constant throughout interphase ^54^ and highly constrained during mitosis ^33, 53^. These local nucleosome motions seem to be mainly driven by thermal fluctuations ^54^.

Although it is important to understand how heterochromatin is physically different from euchromatin, specific labeling methods for euchromatin and heterochromatin in living cells have been limited ^8, 12, 29, 32, 34, 46, 50, 55^ and are still challenging, except for specific heterochromatin regions, such as the nuclear periphery and around nucleoli ^30, 33, 56^. To label euchromatin and heterochromatin specifically in live cells, we have developed a new method called replication-dependent histone labeling (Repli-Histo labeling) based on DNA replication timing ^57, 58^. We investigated local nucleosome motion in the four chromatin classes from euchromatin to heterochromatin (historically named IA, IB, II, and III ^59 60-62^) in living human and mouse cells. We reveal that the more euchromatic regions have a larger nucleosome motion. The nucleosome motion profile or accessibility of each chromatin class from euchromatin to heterochromatin seems to be maintained throughout interphase. Even when the replication timing program is disrupted, genome chromatin is essentially replicated from regions with higher nucleosome motion. Our methodology and findings dissect physical properties of euchromatin and heterochromatin in live cells, relevant to the stochastic nature of DNA replication initiation.

## Results

### Development of the replication-dependent histone (Repli-Histo) labeling

How can we mark euchromatin/heterochromatin specifically in live human cells? To this aim, we took advantage of DNA replication timing, which is a unique feature of eukaryotic DNA replication. Euchromatin replicates in the early S phase and heterochromatin replicates in the late S phase (Fig. 1a)^57, 58^. During this process, newly synthesized histones are provided to the replicated nucleosomes (Fig. 1b). Labeling new histones in the early and late S phases can mark euchromatin and heterochromatin, respectively (Fig. 1c). Based on this concept, we have developed replication-dependent histone (Repli-Histo) labeling.

**Fig. 1:**
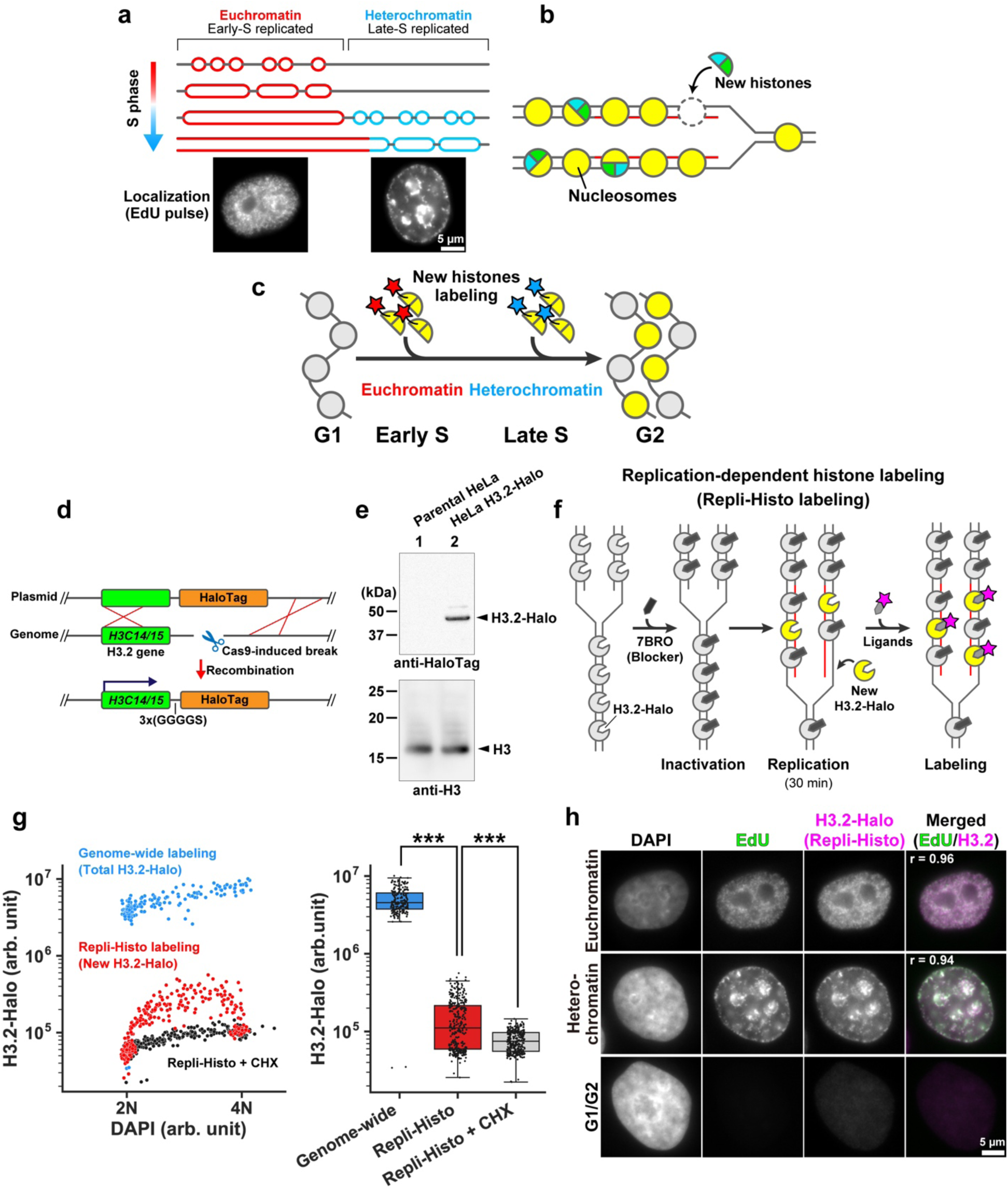
Development of the Repli-Histo labeling. **a**, Eukaryotic cells replicate euchromatin in the early S phase, followed by heterochromatin replication in the late S phase. **b**, Newly synthesized histones are incorporated into the replicated nucleosomes. **c**, Pulse labeling of the new histones in the early and late S phases allows visualization of euchromatin and heterochromatin, respectively. **d**, Schematics of Cas9-mediated HaloTag fusion with endogenous H3.2 genes (H3C14/15). **e**, Western blots of H3.2-Halo in lysates of HeLa cells (lane 2) using an anti-HaloTag antibody (top) and an anti-histone H3 antibody (bottom). **f**, Schematic of the replication-dependent histone labeling (Repli-Histo labeling). **g**, Left: a two-dimensional plot of DAPI and TMR-labeled H3.2-Halo fluorescence in nuclei. Cyan, genome-wide H3.2-Halo labeling (n = 192 cells); red, Repli-Histo labeling (n = 326 cells); black, Repli-Histo labeling with cycloheximide (CHX) treatment (n = 316 cells). Right: the fluorescent intensity of H3.2-Halo on the left. ***: *P* = 1.9 × 10^-77^ for genome-wide H3.2-Halo and Repli-Histo labeling, and *P* = 4.62 × 10^-15^ for Repli-Histo labeling and Repli-Histo labeling + CHX by two-sided Mann-Whitney U-test. **h**, Representative HeLa cells with DAPI staining, EdU pulse labeling, and Repli-Histo labeling. The right column shows the merged image with the Pearson’s correlation coefficient (EdU-Alexa488, green; Repli-Histo labeling, magenta). Top, euchromatin labeling; middle, heterochromatin labeling; bottom, G1/G2 cell without EdU/Repli-Histo signals.

To label nucleosomes, we focused on one of the core histone variants, H3.2, which is expressed in a replication-dependent manner during S phase and incorporated into nucleosomes ^63-66^. Once incorporated, H3.2 is stable for hours without exchange ^35^, allowing us to trace the H3.2 incorporated region. Using the CRISPR/Cas9 system ^67^, we introduced a HaloTag sequence at the C-terminus of endogenous H3.2 genes (Fig. 1d). The HaloTag can be visualized with the HaloTag ligand tetramethylrhodamine (TMR) or other Janelia Fluor (JF) dyes ^68^. The cell clones stably expressing H3.2-HaloTag (H3.2-Halo) were labeled with TMR and were isolated by fluorescence-activated cell sorting (FACS) (Fig. S1a) ^69^. The proper insertion of the tag sequence and the expression of H3.2-Halo were verified by polymerase chain reaction (PCR) (Fig. S1b) and western blotting (Figs. 1e and S1c). Expressed H3.2-Halo comprises about 20% of the endogenous H3.1/H3.2 (Fig. S1c). The distribution of expressed H3.2-Halo in HeLa cells resembles DAPI staining (r = 0.84, Fig. S1d), suggesting that expressed H3.2-Halo is stochastically incorporated into nucleosomes genome-wide during the S phase, consistent with the previous reports ^70, 71^. Stepwise salt washing of nuclei isolated from the H3.2-Halo expressing cells confirmed that the biochemical behavior of H3.2-Halo is like that of endogenous H3.1/H3.2 (Fig. S1e), suggesting that H3.2-Halo was properly incorporated into the nucleosomes in the expressed cells.

For Repli-Histo labeling, we blocked all the existing H3.2-Halo in the cell with non-fluorescent HaloTag ligand, 7-bromo-1-heptanol (7BRO) ^72^ and inactivated them (Fig. 1f). In S phase cells, the new H3.2-Halo was synthesized and incorporated into new nucleosomes (Fig. 1g) ^63-66, 73^. Then, we pulse-labeled the new H3.2-Halo nucleosomes with fluorescent ligands TMR or other JF dyes for 30 min (Fig. 1f). Repli-Histo labelings in the early and late S phases highlight euchromatin and heterochromatin, respectively (Figs. 1h and right in S1f). Euchromatic or heterochromatic labeling was judged from its labeling pattern because the replicated chromatin in the nucleus shows unique patterns depending on when the region replicated (Figs. 1h and right in S1f) ^59 60-62^. The Repli-Histo signals with H3.2-Halo-TMR were nicely colocalized with nascent DNA pulse-labeled with 5-ethynyl-2′-deoxyuridine (EdU) (Fig. 1h, r = 0.96 in euchromatin and r = 0.94 in heterochromatin). The Repli-Histo signals were abolished by a protein synthesis inhibitor cycloheximide (CHX), validating that the Repli-Histo labeling marks only newly synthesized H3.2-Halo (Fig. 1g, black).

### Genomic localization of Repli-Histo labeled nucleosomes

To verify that Repli-Histo labeling specifically marks euchromatin and heterochromatin in live HeLa cells, we determined the genomic localizations of nucleosomes labeled with Repli-Histo labeling (Fig. 2a). We first synchronized HeLa cells at the G1/S boundary by double thymidine block. After a 1-h (for euchromatin labeling) or 5-h (for heterochromatin) release from the block (Fig. 2a(i)), we inactivated the existing H3.2-Halo with 7BRO and chased for 1 h with new H3.2-Halo incorporation (Fig. 2a(ii)). Then the new H3.2-Halo-nucleosomes from isolated nuclei (Fig. 2a(iii)) were conjugated with HaloTag PEG-biotin ligand (Fig. 2a(iv)). The nucleosomes with the H3.2-Halo-PEG-biotin were pulled down using streptavidin magnetic beads (Figs. 2a(iv) and S2a). Nucleosomal DNAs were purified (Figs. 2a(v) and S2b) and subjected to sequencing (Figs. 2b and S2c-d) as done previously ^50^.

**Fig. 2:**
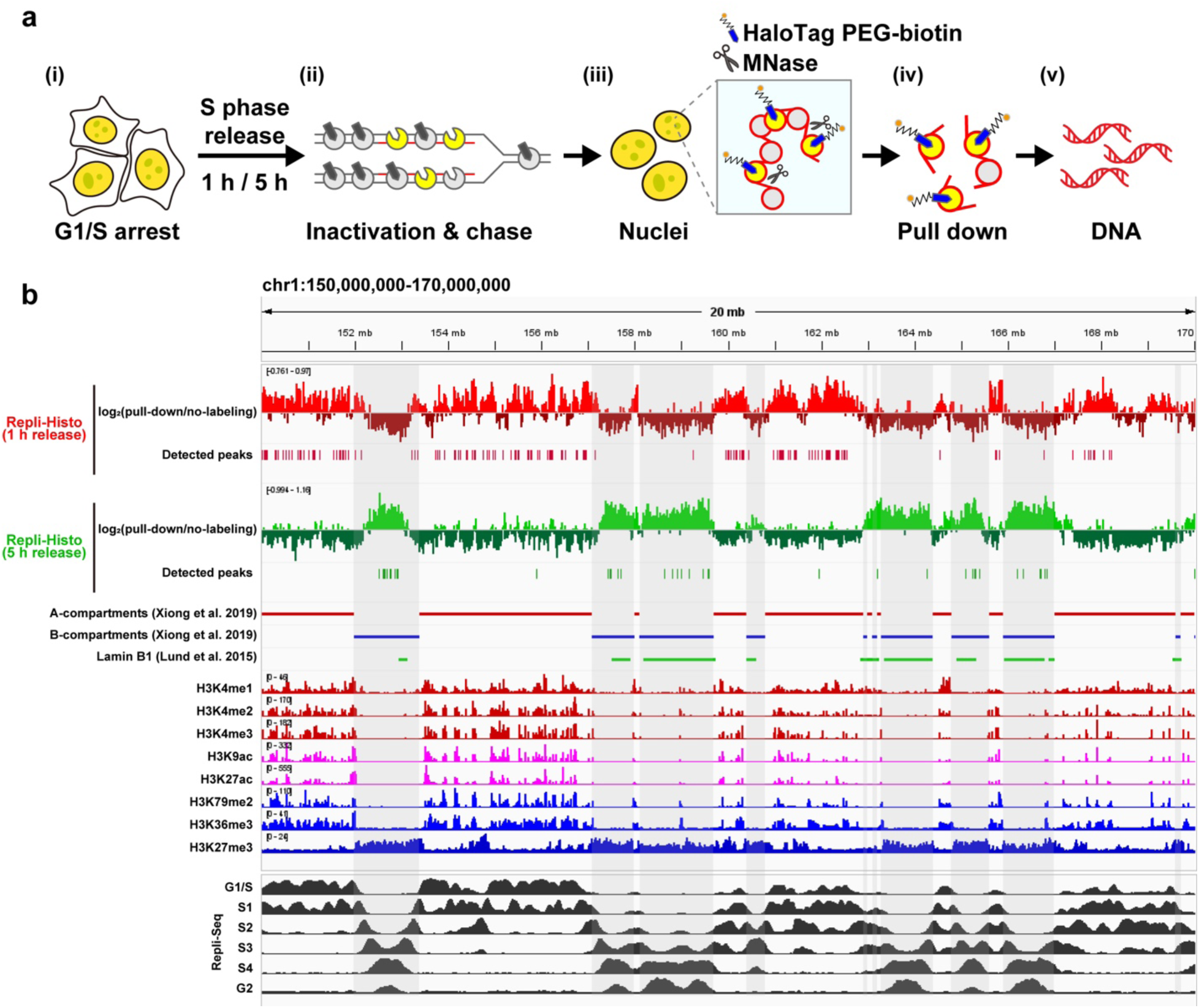
Genomic localizations of the Repli-Histo labeled genomic regions. **a**, A purification scheme for the Repli-Histo-labeled H3.2-Halo-nucleosomes and their associated genomic DNA. The purified DNA fractions were indexed, amplified, and sequenced. **b**, Genomic distribution of the Repli-Histo-labeled H3.2-Halo at 1 h and 5 h after release from double thymidine blocks; view from chr1_150,000,000-170,000,000 (also see Fig. S2d for another genomic region). The first and second rows show the labeled H3.2-Halo enrichment relative to a no-labeling control (log2 ratio) at 10-kb bins. The gray stripes indicate the Hi-C B-compartments^75^. Previously published data are also shown as indicated on the left.

While peaks of control H2B-Halo nucleosome data ^50^ and H3.2-Halo without 7BRO treatment distributed genome-wide (Fig. S2c), we identified 3,750 nucleosome peaks associated with H3.2-Halo of the 1-h-release fraction and 1,211 nucleosome peaks of the 5-h-release fraction in the annotated HeLa S3 genome (hg19) (Figs. 2b and S2c-d). The nucleosome peaks of the 1-h-release and 5-h-release fractions are distributed complimentary, consistent with the previous Repli-Seq data (^74^; https://doi.org/doi:10.17989%2FENCSR647UES). A total of 93.6% of the 1-h-release Repli-Histo fraction belonged to the Hi-C A-compartment ^20, 75^, and 91.8% of the 5-h-release fraction overlapped with the Hi-C B-compartment ^20, 75^. The 1-hr-release and 5-h-release Repli-Histo fractions were significantly enriched with active chromatin marks (H3K4me1–3, H3K9ac, H3K27ac, H3K36me3, and H3K79me2) and an inactive chromatin mark (H3K27me3), respectively (Fig. S3a-j) ^74, 76^, in good agreement with previous Repli-Seq and related studies ^57, 77, 78^. Consistent with our imaging data (Fig. 1h), the 5-h-release Repli-Histo fraction was associated with lamina-associated domains (45.5% of the fraction; Figs. 2b, S2d, and S3k)^79^. Taken together, our genomics analysis indicates that Repli-Histo labeling (1-h-release and 5-h-release fractions) specifically highlights the euchromatin (Hi-C A-compartment) and heterochromatin (Hi-C B-compartment), respectively.

### Single-nucleosome imaging in euchromatin and heterochromatin regions in living cells

Next, we combined Repli-Histo labeling and single-nucleosome imaging ^47, 48, 50, 52^ to measure local nucleosome motion in euchromatin and heterochromatin. We conducted Repli-Histo labeling on asynchronous HeLa cells (left, Fig. 3a). In this case, about 46% of the cells were in the S phase (Fig. S1f) and were labeled with Repli-Histo labeling (left, Fig. 3a). Here, we used two colors for the HaloTag ligands to visualize the labeling patterns (a high concentration of TMR) and to image single nucleosomes (a very low concentration of JF646) ^68^ (right, Fig. 3a). After DNA replication, we monitored the nucleosome motion at the G2 phase (right, Fig. 3a), excluding the possibility that the different DNA content and nuclear size affect the result.

**Fig. 3:**
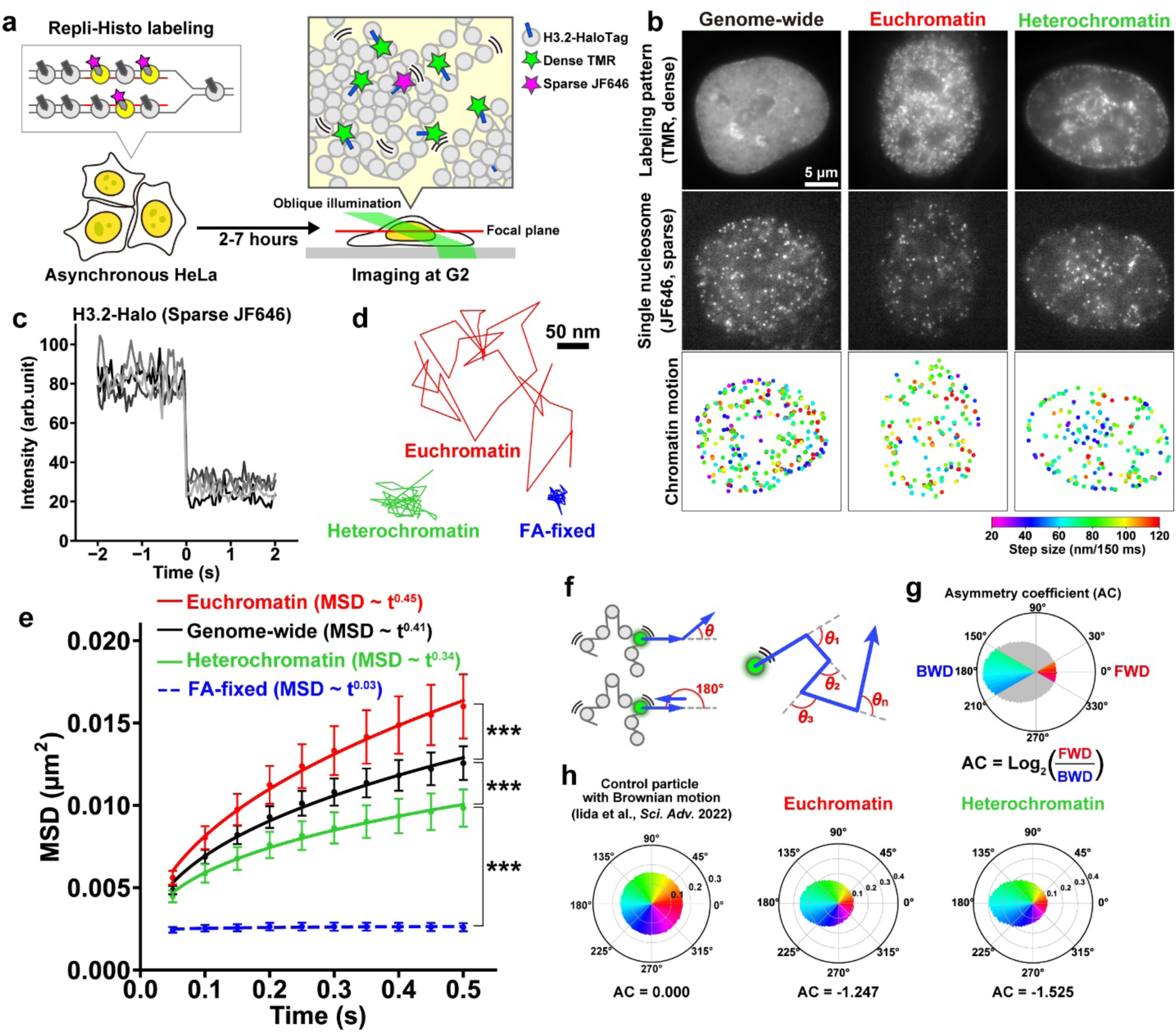
Euchromatin-/heterochromatin-specific single-nucleosome imaging with Repli-Histo labeling. **a**, Schematic of the Repli-Histo single-nucleosome imaging. Left: The S phase HeLa cells (∼46% of the asynchronous population; Fig. S1f) were labeled with Repli-Histo labeling. Right: The Repli-Histo-labeled cells were observed at G2 phase using oblique illumination microscopy system. Only a thin optical layer (red) within the nucleus was illuminated with a low background using a light sheet (green). A small fraction of H3.2-Halo was fluorescently labeled with JF646 (magenta star) for single-nucleosome imaging, and the rest of the H3.2-Halo was densely labeled with TMR (green stars) to visualize the labeling pattern. **b**, HeLa cells with two-color HaloTag labeling. Left: genome-wide H3.2-Halo labeling. Center and right: Repli-Histo labeling with euchromatin and heterochromatin patterns, respectively. Top row: the labeling patterns with dense TMR labeling, middle row: single nucleosomes with sparse JF646 labeling, bottom row: the displacement of single nucleosomes as 2D heatmaps (^8, 56^). Here, larger movements appear as more ‘‘red’’ (or hot), and smaller movements appear as more ‘‘blue’’ (or cold) colors (see the color bar at the bottom right). **c**, Single-step photobleaching of five representative nucleosome (H3.2-Halo-JF646) dots. The horizontal axis shows the time before and after photobleaching. **d**, Representative trajectories of tracked single nucleosomes in euchromatin (red) and heterochromatin (green) in live HeLa cells and in formaldehyde (FA)-fixed HeLa cells (blue). **e**, MSD plots (± SD among cells) of single nucleosomes in euchromatin-labeled (red, n = 45), heterochromatin-labeled (green, n = 36), and genome-wide-labeled (black, n = 38) living HeLa cells from 0.05 to 0.5 s. Blue, MSD from FA-fixed control cells (n = 30). The plots were fitted as a sub-diffusive curve. ***: *P* = 2.0 × 10^-16^ (euchromatin vs. genome-wide), *P* = 7.9 × 10^-12^ (genome-wide vs. heterochromatin), *P* = 3.6 × 10^-19^ (heterochromatin vs. FA-fixed) by the two-sided Kolmogorov-Smirnov test. **f**, Schematic for angle-distribution analysis. **g**, Schematic for the asymmetric coefficient (AC). See Methods for details. AC shows deviation from a homogeneous distribution and is negative for angular distributions, where pulling back force is dominant. **h**, The angle distribution data from a particle with Brownian motion (left), nucleosomes in euchromatin-labeled (center, n = 45 cells), and heterochromatin-labeled (right, n = 36 cells) living HeLa cells. Their AC values are shown at the bottom. The control data was reproduced from Iida et al. (2022) ^54^.

Using oblique illumination microscopy, which allowed us to illuminate a thin area within a single nucleus (right, Fig. 3a) ^47, 80^, we observed individual nucleosomes in euchromatin and heterochromatin as clear dots and recorded their motions at 50 ms per frame (200 frames, 10 s in total) (Fig. 3b; Movies S1 and S2). Each JF646 dot showed a single-step photobleaching (Fig. 3c) while TMR dots did not (Fig. S4a), confirming that each JF646 dot represented a single H3.2-Halo-JF646 molecule in a single nucleosome. The individual dots were fitted with a 2D Gaussian function to estimate the precise position of the nucleosome ^81-83^ and were tracked using u-track software ^84^ to obtain the nucleosome trajectory data (Fig. 3d). The position determination accuracy of fluorescent H3.2-Halo-JF646 dots was 9.5 nm (Fig. S4b and see Methods). We only characterized the behavior of the H3.2-Halo stably incorporated into nucleosomes because the free H3.2-Halo molecules moved too quickly to be tracked with a 50 ms/frame image acquisition. We also confirmed that 7BRO treatment did not significantly affect nucleosome motion (Fig. S4c).

Fig. 3b (bottom) displays the heat maps of nucleosome motions in euchromatin and heterochromatin (more movements show more red, and fewer movements with more blue). From the tracked nucleosome trajectories, we calculated the nucleosome displacement (Fig. S4d), and then the mean square displacement (MSD) (Fig. 3e), which shows the spatial extent of motion in a certain time window (Fig. S4e)^85^. The MSD plots show a sub-diffusive curve, and their motions were severely suppressed after chemical fixation with formaldehyde (FA) (Fig. 3e; Movie S3). The plots indicate that nucleosomes in euchromatin are more dynamic than heterochromatin (Fig. 3e; Movies S1 ad S2), while the control MSD data of H2B-Halo ^54^ and H3.2-Halo without 7BRO treatment are in the middle of those of euchromatin and heterochromatin (Genome-wide in Fig. 3e; Movie S4). The MSD exponent in euchromatic nucleosomes (0.45) is higher than that of heterochromatic nucleosomes (0.34) (Figs. 3e and S4f). Furthermore, we examined the moving angle (motion vector) distribution of individual nucleosomes (Fig. 3f) and calculated the asymmetry coefficient, AC, value (Fig. 3g). Nucleosomes in heterochromatin have more pulling back force (i.e., a smaller AC value, Figs. 3h and S4g) and are more constrained than in euchromatin.

### The more-euchromatic regions (earlier replicated regions) have a larger nucleosome motion

To obtain more information on the nucleosome behaviors in euchromatin and heterochromatin, we categorized whole chromatin into four regions from euchromatin to heterochromatin based on the replication timing and labeling pattern (Figs. 4a and S5a-c; Movies S5-S8) (i.e., replication foci; see Table 1 for more details) ^60, 61, 62^. Historically, the four classified regions were named Class IA and IB, which are earlier replicated regions (more euchromatic), and Class II and III, which are later replicated regions (more heterochromatic) ^60, 61, 62^. Although the cells with the Class III pattern have a large amount of freely diffusing H3.2-Halo (Movie S8; Figs. 4a and S5b), we only focused on the behavior of the H3.2-Halo stably incorporated into nucleosomes in the Class III pattern (Fig. S6a).

**Fig. 4:**
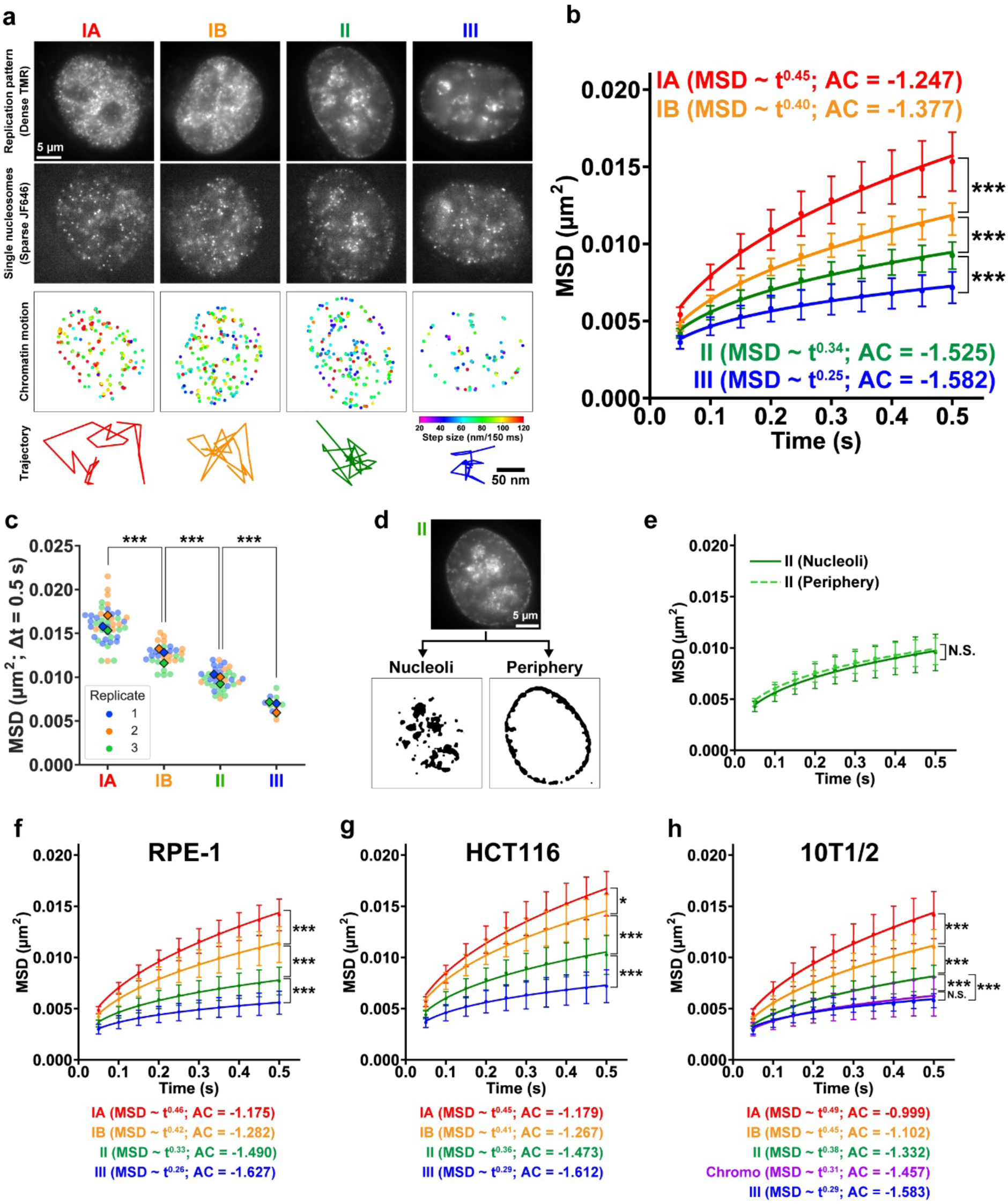
Local nucleosome movements in the four classes of chromatin regions. **a**, HeLa cells with two-color Repli-Histo labeling of the chromatin regions Class IA, IB, II, and III. 1st row: labeling patterns with dense TMR labeling; 2nd row: single nucleosomes with sparse JF646 labeling; 3rd row: displacement of single nucleosomes shown as 2D heatmaps; 4th row: representative trajectories of tracked single nucleosomes. **b**, MSD plots (± SD among cells) of single nucleosomes in Class IA (red, n = 45), IB (orange, n = 29), II (green, n = 36), and III (blue, n = 10) in living HeLa cells from 0.05 to 0.5 s. Classes IA and II are reproduced from Fig. 3e (euchromatin and heterochromatin). MSD exponents from Fig. S6d and the asymmetry coefficient (AC) from Fig. S6e are also shown in the brackets. ***: *P* = 1.3 × 10^-11^ for IA versus IB, *P* = 3.8 × 10^-10^ for IB versus II, *P* = 6.5 × 10^-7^ for II versus III by the two-sided Kolmogorov-Smirnov test. **c**, MSD values at 0.5 s in each cell from (**b**). Blue, orange, and green dots show three biological replicates. Large diamonds show the mean values in each replicate. **d**, Sub-categorization of the Class II chromatin. Nucleosome motions in the nucleolus-associated region (bottom left) and peri-nuclear region (bottom right) are individually analyzed in (**e**). **e**, MSD plots (± SD among cells) of single nucleosomes in the Class II nucleolus-associated region (Nucleoli, n = 22) and peri-nuclear region (Periphery, n = 22). N.S.: *P* = 0.87 by Kolmogorov-Smirnov test. **f**–**h**, MSD plots (± SD among cells) of single nucleosomes in Class IA (red), IB (orange), II (green), and III (blue) in various cell lines from 0.05 to 0.5 s. MSD exponents and the asymmetry coefficients (AC) are also shown in the brackets. **f**, RPE-1 cells, ***: *P* = 8.6 × 10^-8^ for Class IA (n = 36) versus IB (n = 31), *P* = 2.9 × 10^-8^ for Class IB versus II (n = 29), and *P* = 2.1 × 10^-4^ for Class II versus III (N= 19) by the two-sided Kolmogorov-Smirnov test. **g**, HCT116 cells, *: *P* = 0.024 for Class IA (n = 29) versus IB (n = 21), ***: *P* = 3.5 × 10^-6^ for Class IB versus II (n = 22) and *P* = 1.0 × 10^-4^ for Class II versus III (n = 20) by the two-sided Kolmogorov-Smirnov test. **h**, Mouse 10T1/2 cells, ***: *P* = 1.1 × 10^-8^ for Class IA (n = 74) versus IB (n = 39), *P* = 7.4 × 10^-10^ for Class IB versus II (n = 43), *P* = 9.3 × 10^-4^ for Class II versus III (n = 5), and *P* = 1.4 × 10^-6^ for Class II versus chromocenter (n = 27) by the two-sided Kolmogorov-Smirnov test. N.S.: *P* = 0.63 for Class III versus chromocenter.

**Table 1:**
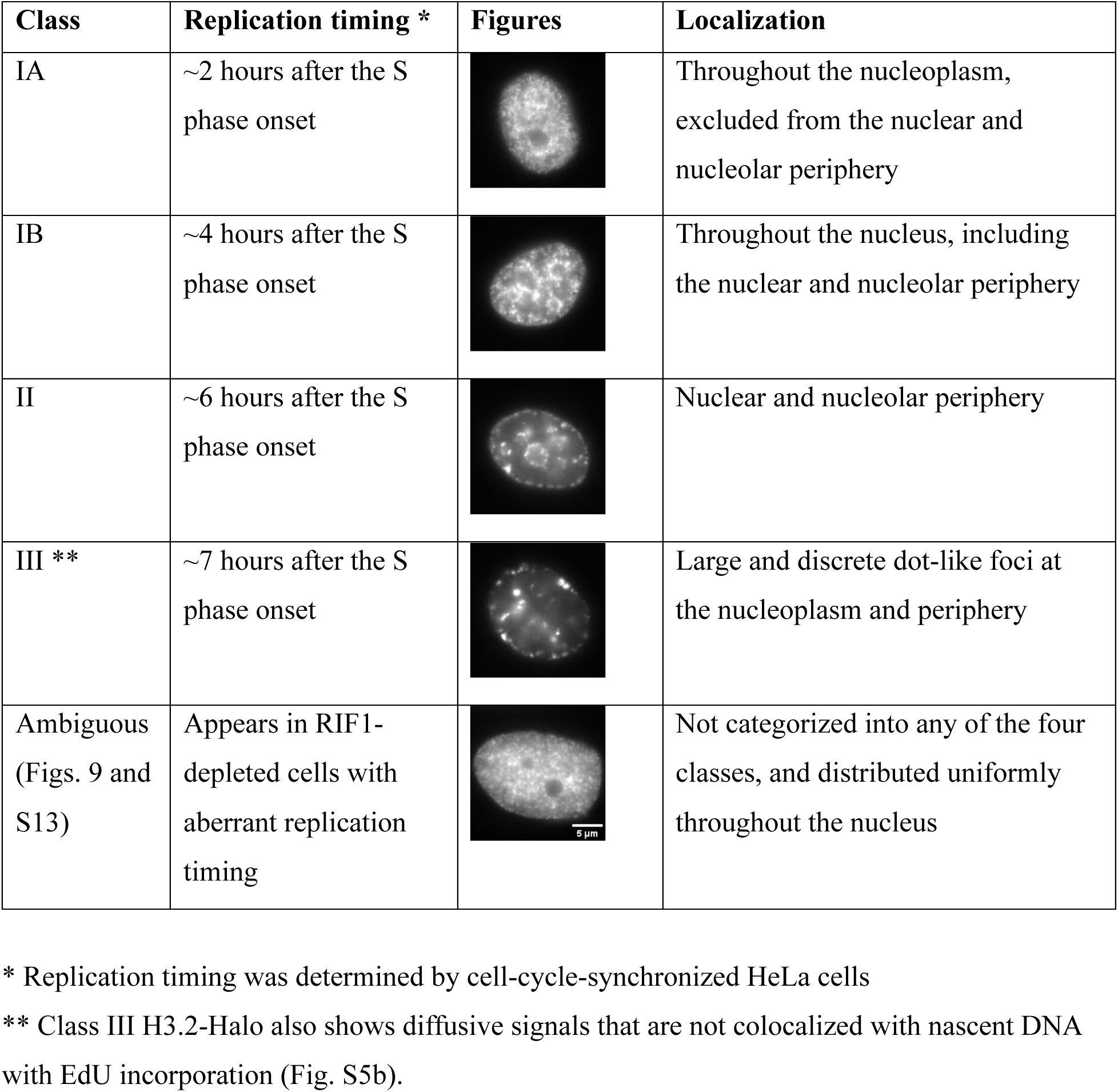
Description of the replication pattern localization.

| Class | Replication timing * | Figures | Localization |
| --- | --- | --- | --- |
| IA | ~2 hours after the S phase onset |  | Throughout the nucleoplasm, excluded from the nuclear and nucleolar periphery |
| IB | ~4 hours after the S phase onset |  | Throughout the nucleus, including the nuclear and nucleolar periphery |
| II | ~6 hours after the S phase onset |  | Nuclear and nucleolar periphery |
| III ** | ~7 hours after the S phase onset |  | Large and discrete dot-like foci at the nucleoplasm and periphery |
| Ambiguous (Figs. 9 and S13) | Appears in RIF1-depleted cells with aberrant replication timing |  | Not categorized into any of the four classes, and distributed uniformly throughout the nucleus |

\* Replication timing was determined by cell-cycle-synchronized HeLa cells
\*\* Class III H3.2-Halo also shows diffusive signals that are not colocalized with nascent DNA with EdU incorporation (Fig. S5b).

Interestingly, nucleosome motion is progressively more constrained as the Class progresses from IA through III (Figs. 4b and S6b; Movies S5-S8, see also Fig. S7). The radius of constraint (Rc; p = 6/5 × Rc^2^, where p is the plateau value of the MSD) ^85^ of nucleosomes in each Class was estimated (mean ± SD): 132 ± 17 nm (IA), 115 ± 8.6 nm (IB), 105 ± 12 nm (II), and 89 ± 14 nm (III) (Fig. S6c). Their MSD exponents α in the time range between 0 and 0.5 s gradually decreased: IA, 0.45; IB, 0.40; II, 0.34; III, 0.25 (Figs. 4b and S6d). Decreases in AC values were also observed (Figs. 4b and S6e): IA, -1.247; IB, -1.377; II, -1.525; III, -1.582. These data indicate that the more euchromatic regions (earlier replicated regions) have a larger nucleosome motion with higher MSD exponents and AC values. Notably, almost no overlaps or gaps were observed among the four classified regions (Fig. 4b-c). Interestingly, the nucleosomes around the nuclear periphery (i.e., lamina-associated domains, LADs ^86^) and around nucleoli (nucleolus-associated domains, NADs ^87^) in Class II (Fig. 4d) showed similar MSD values (Fig. 4e), suggesting that chromatin replicated at the same time exhibits a similar local nucleosome behavior, regardless of its intranuclear location.

To investigate the generality of the motion profile in euchromatin/heterochromatin (Class IA, IB, II, and III), we tagged the endogenous H3.2 gene in RPE-1 (Fig. S8a) and HCT116 cells (Fig. S8b) with HaloTag. We also tagged the endogenous H3.1 gene in mouse 10T1/2 fibroblast cells (Fig. S8c) ^88, 89^ to examine pericentromeric heterochromatin foci (chromocenters) ^90^. Repli-Histo labeling was performed in various stages of their S phase cells (Fig. S8d-f) after validation by PCR and western blotting (Fig. S8a-c). We observed their nucleosome motions at the G2 phase. We found a similar motion profile in not only HeLa but also RPE-1, HCT116, and mouse 10T1/2 cells (Fig. 4f-h; Movies S9-12). Interestingly, nucleosomes in the pericentromeric heterochromatin behave similarly to those of Class III in 10T1/2 cells (Fig. 4h). Nucleosomes in the putative inactive X chromosome in RPE-1 cells, which are highly condensed, also seem to belong to Class III and are highly constrained (Fig. S8d; Movie S12). Overall, the more euchromatic regions (earlier replicated regions) have larger nucleosome motions with higher MSD exponents and AC values (Fig. 4f-h). The nucleosome motion profiles (MSD exponents and AC values) in euchromatin/heterochromatin seem to be a general feature in mammalian cells.

### The nucleosome motion profile of each chromatin Class is maintained throughout interphase

We wondered whether we would see a similar motion profile in the G1 phase after cell division. To this aim, we arrested the Repli-Histo labeled cells at the late-G1 phase using L-mimosine (Fig. 5a) ^91^. We found that the Repli-Histo labeling patterns were retained among the G1-arrested cells (Fig. 5b), consistent with previous reports that replication foci patterns are retained throughout several cell cycles ^40, 92^. Again, the Rc, MSD exponents α, and AC values all gradually decreased as Class progressed from IA through III (Figs. 5c and S9; Movies S13-16). The nucleosome motion profile of each chromatin Class (IA, IB, II, and III) was maintained between the G1 and G2 phases. This finding is consistent with our previous data, which shows that the average motion of nucleosomes labeled by H2B-Halo in the G1 and G2 phases is very similar ^54^.

**Fig. 5:**
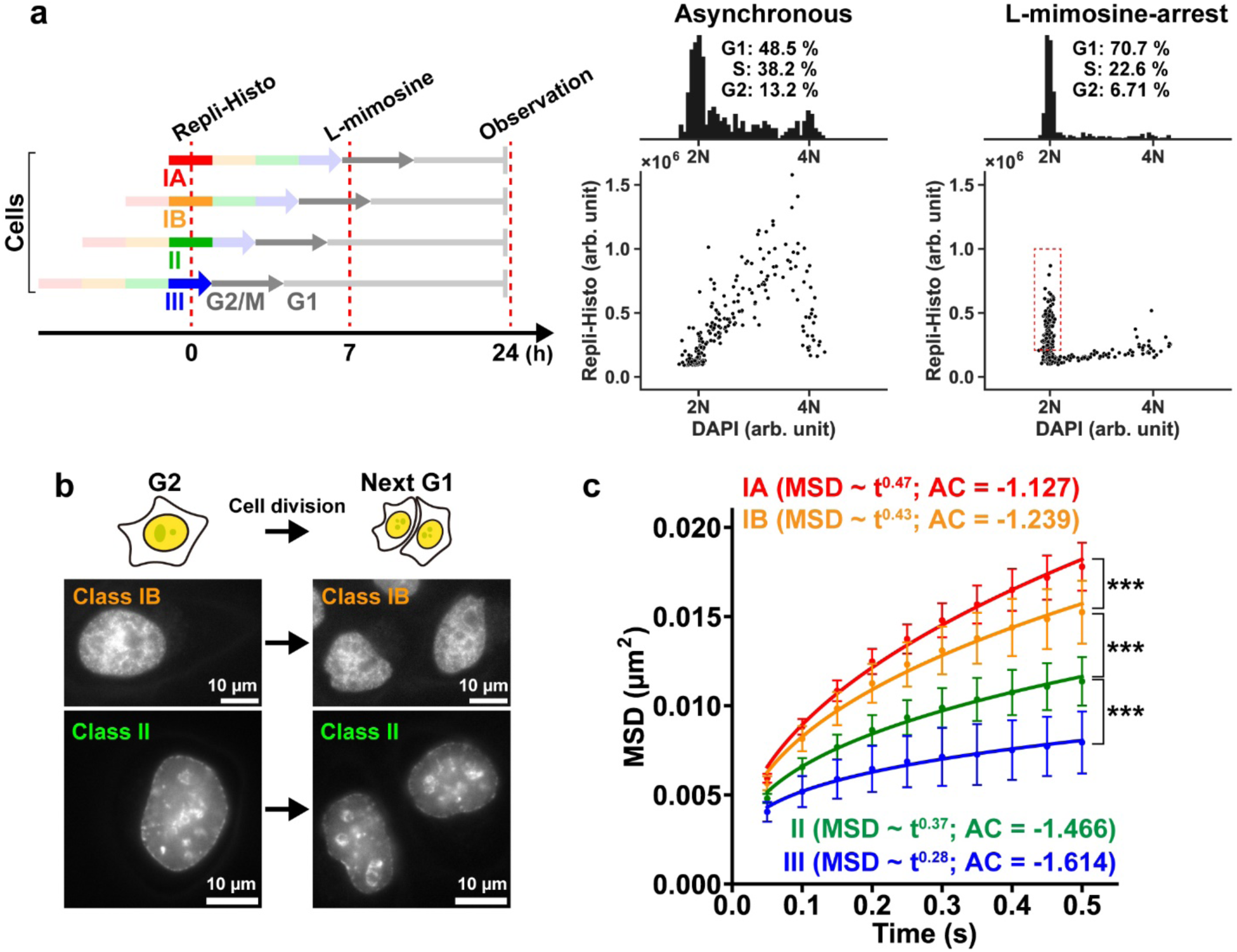
The nucleosome motion profile of each chromatin Class in the G1 phase. **a**, Left: schematics of the G1 synchronization using L-mimosine. Right: two-dimensional plots of DAPI and Repli-Histo-labeled H3.2-Halo fluorescence in asynchronous and L-mimosine-treated HeLa cells. The top shows the histograms of the DAPI intensity indicating cell cycle profiles. The cell cycle profiles were estimated by the Dean-Jett-Fox method ^150^. Most of the Repli-Histo-labeled cells are arrested at the G1 phase (red rectangle) in L-mimosine-treated cells. **b**, Time-lapse imaging of HeLa cells with Repli-Histo labeling. The Repli-Histo labeling patterns are retained among the G1 cells after cell division. **c**, MSD plots (± SD among cells) of single nucleosomes in Class IA (red, n = 37), IB (orange, n = 18), II (green, n = 15), and III (blue, n = 10) in G1-arrested living HeLa cells from 0.05 to 0.5 s. MSD exponents and the asymmetry coefficient (AC) values from Fig. S7 are also shown in the brackets. ***: *P* = 4.5 × 10^-6^ for Class IA versus IB, *P* = 2.4 × 10^-4^ for Class IB versus II, *P* = 4.6 × 10^-5^ for Class II versus III by the two-sided Kolmogorov-Smirnov test.

Next, we questioned whether the nucleosome motion profile of each chromatin Class is also maintained throughout the S phase. A significant fraction of the H3.2-Halo in the cells with the Class III pattern diffused freely (Movie S8; Fig. S5b), and this fraction disturbed the observation at the subsequent S phase. We thus fused HaloTag to the C-terminus of endogenous CDC45, the component of active DNA helicase, which stably binds to the replicating chromatin region ^93^, to generate HeLa cells expressing endogenous CDC45-HaloTag (Fig. 6a). We confirmed that all endogenous CDC45 were tagged with HaloTag (Figs. 6b and S10a-b) in a clone, which proliferated normally (Fig. S10c), confirming that the CDC45-Halo is functional. While the labeled CDC45-Halo was diffusive in the G1/G2 phase (2^nd^ row in Fig. 6c and Movie S17), CDC45-Halo formed several specific foci patterns during the S phase (2^nd^ row in Fig. 6c), which were analogous to those of the EdU-labeled nascent DNA (replication foci), depending on the replication timing (3^rd^ row in Fig. 6c) ^94-96^. They correspond to Class IA, IB, II, and III. We performed single-molecule imaging of CDC45-Halo stably bound fractions (Fig. 6d and Movies S18-21) to track the local behaviors of active replication sites in the various S phase stages. Observed MSD values of CDC45-Halo throughout the S phase stages were akin to those of single nucleosomes in corresponding Class IA, IB, II, and III in the G2 phase (Figs. 6d and S10d-e). We concluded that the local motion profile of each chromatin Class is maintained throughout interphase, even where DNA replication is ongoing.

**Fig. 6:**
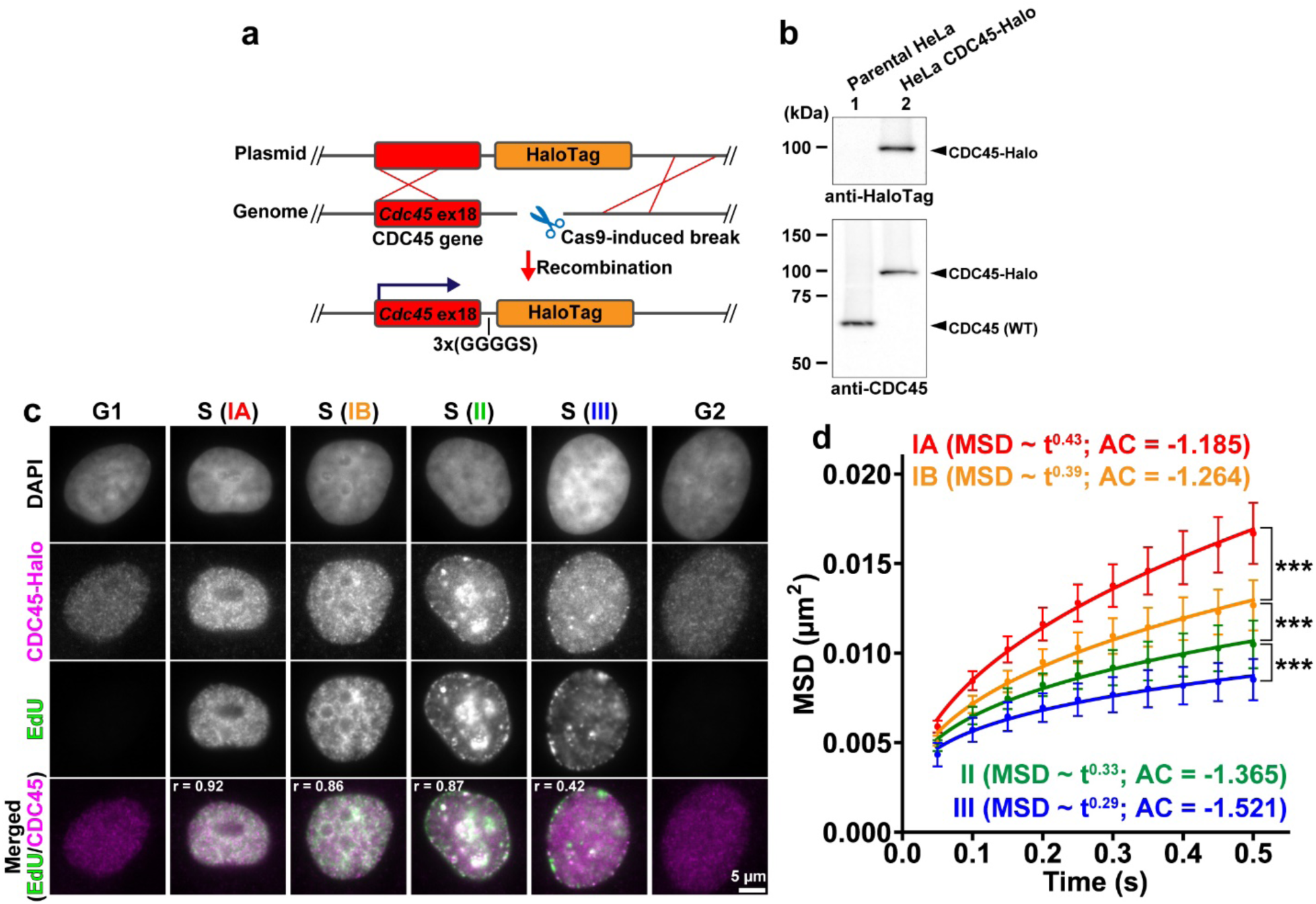
Single-CDC45 imaging reveals the nucleosome motion profile during the S phase. **a**, Schematics of Cas9-mediated HaloTag fusion with the endogenous CDC45 gene (Cdc45). **b**, Western blots of CDC45-Halo in lysates of HeLa cells using an anti-HaloTag antibody (top) and an anti-CDC45 antibody (bottom). Lane 1: parental HeLa cells, lane 2: HeLa cells expressing endogenous CDC45-Halo. **c**, HeLa cells with CDC45-Halo labeled with JF646 (2^nd^ row) and EdU-pulse labeling (3^rd^ row). Chromatin classes (IA, IB, II, and III) were categorized as described in Table 1. The bottom row shows the merged image of EdU (green) and CDC45-Halo (magenta) with their corresponding Pearson’s correlation coefficients. **d**, MSD plots (± SD among cells) of single CDC45-Halo with Class IA (red, n = 41), IB (orange, n = 15), II (green, n = 23), and III (blue, n = 24) patterns in living HeLa cells from 0.05 to 0.5 s. MSD exponents and the asymmetry coefficient (AC) values from Fig. S10d-e are also shown in the brackets. ***: *P* = 1.2 × 10^-5^ for Class IA versus IB, *P* = 7.7 × 10^-4^ for Class IB versus II, *P* = 2.0 × 10^-5^ for Class II versus III by the two-sided Kolmogorov-Smirnov test.

### Repli-Histo labeling dissects physical properties of euchromatin and heterochromatin in live human cells

To further investigate the physical natures of euchromatin and heterochromatin using Repli-Histo labeling, we focused on the distances between two nucleosomes in close proximity and examined the mean squared distance between them (two-point MSD) (Fig. 7a) ^12, 97^ because the distance between nucleosomes is much less affected by either translational or rotational movements of the chromatin domain during imaging. We performed Repli-Histo labeling with a very low concentration of TMR and JF646 and tracked two spatially neighboring single-nucleosomes. Cell-cycle synchronization allowed the labeling of euchromatic (IA) and heterochromatic (II) domains (Fig. S5c; see also Methods). Single-nucleosomes labeled with TMR and JF646 were imaged using a beam splitter system (Figs. 7b and S11a).

**Fig. 7:**
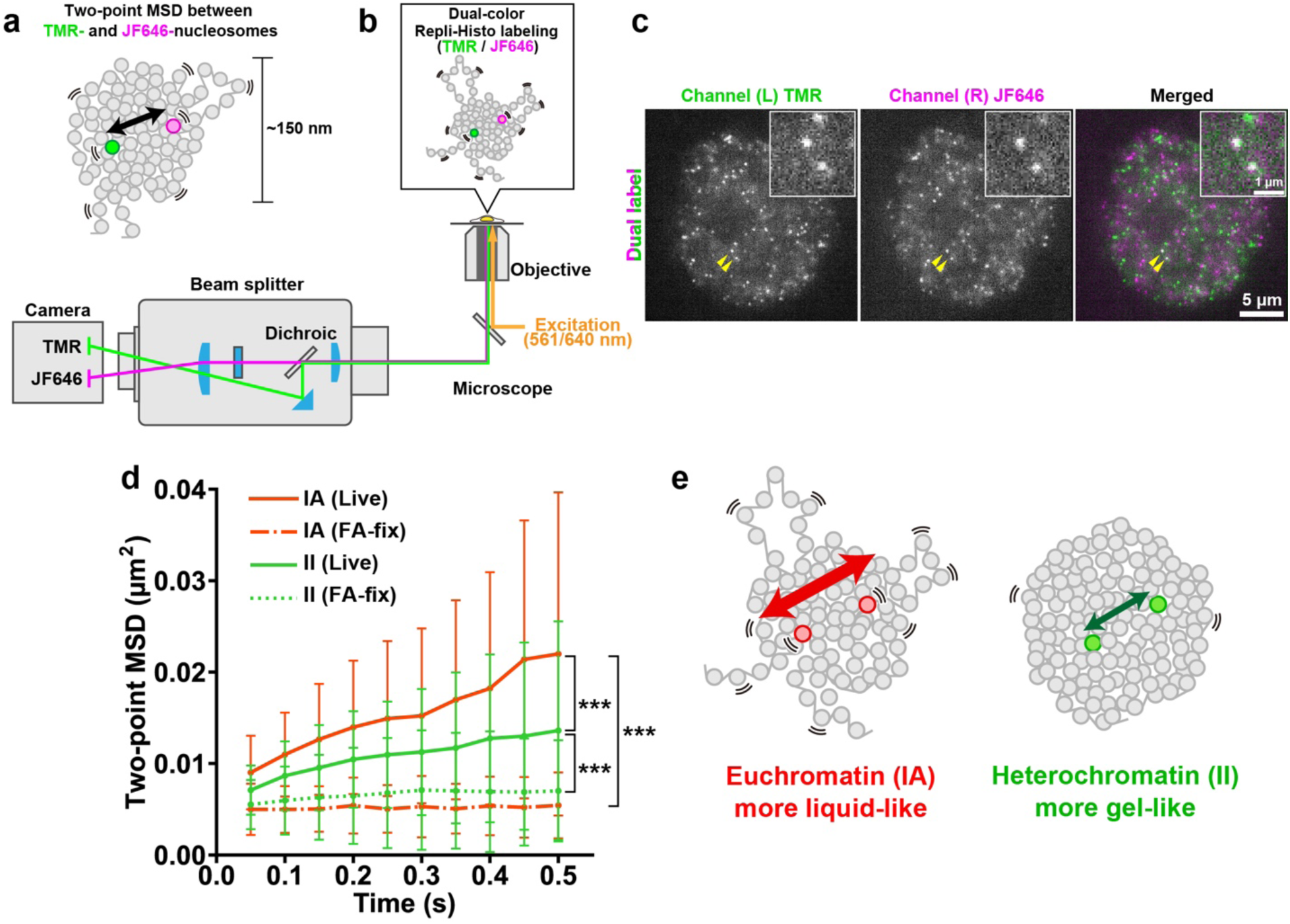
Two-point MSD analysis to dissect the physical properties of euchromatin and heterochromatin. **a**, Schematic for two-point MSD analysis. The mean square relative distances between the TMR- and JF646-nucleosomes were tracked. The neighboring dot pairs whose averaged distances were < 150 nm were analyzed. **b**, Schematic for dual-color imaging with a beam splitter system (W-VIEW GEMINI, Hamamatsu Photonics). The images of two single nucleosomes with different colors were simultaneously acquired with a single sCMOS. **c**, Representative images of single-nucleosomes simultaneously labeled with TMR (left) and JF646 (center) by Repli-Histo labeling in living HeLa cells. Right: the merged image. The two pairs of nucleosomes in close proximity (yellow arrowheads) are enlarged (inset). See also Movie S23. **d**, Two-point MSD plots (mean ± SD among dot pairs) between TMR- and JF646-nucleosomes in HeLa cells with indicated conditions: Repli-Histo labeling with Class IA pattern in live cells (red solid line, n = 48 pairs); Class IA pattern in FA-fixed cells (red dashed line, n = 63 pairs); Class II pattern in live cells (green solid line, n = 168 pairs); Class II pattern in FA-fixed cells (green dashed line, n = 140 pairs). ***: *P* = 6.7 × 10^-4^ for Class IA (live) versus II (live), *P* = 7.4 × 10^-17^ for Class IA (live) versus IA (FA-fixed), and *P* = 1.5 × 10^-11^ for Class II (live) versus II (FA-fixed) by the two-sided Kolmogorov-Smirnov test. **e**, Schematic of how euchromatin behaves more liquid-like with larger nucleosome fluctuation inside the condensed chromatin domain than heterochromatin.

We simultaneously observed clear dots in each color channel (Fig. 7c and Movie S22; position determination accuracy was 11.2 nm (TMR) and 10.9 nm (JF646) in Fig. S11b). Note that there was essentially no cross-talk between the TMR and JF646 signals (Fig. S11c-d). Among them, we found many pairs of nucleosomes with TMR and JF646 signals located in close proximity (insets, Fig. 7c; Movie S23). We collected nucleosome pairs whose distance is less than 150 nm (Fig. 7a; Movies S22 and S24), which are likely in the same chromatin domain ^12^, and analyzed their motions. Two-point MSD plots showed rather sub-diffusive curves (Fig. 7d), suggesting that nucleosomes fluctuate inside the chromatin domain like a liquid. This fluctuation was severely suppressed after the chemical fixation with FA (Fig. 7d, dashed lines; Movies S25-26). The nucleosome pairs in euchromatin domains showed larger two-point MSD than in heterochromatin domains (Fig. 7d, red and green solid lines). Two-point MSD plots using Repli-Histo labeling suggest nucleosomes inside the euchromatin domain fluctuate more than those in the heterochromatin domain (Fig. 7e). Nucleosomes in heterochromatin are more constrained and seem closer to a gel-like state, presumably due to more crosslinks such as HP1.

### Local nucleosome motion facilitates chromatin accessibility to large proteins

Our findings using Repli-Histo labeling revealed that genome chromatin is primarily replicated from regions with greater nucleosome motions (Fig. 4b and 4f-h). We hypothesized that local nucleosome fluctuation governs chromatin accessibility to a limited number of replication initiation factors to fire replication origins (Fig. 8a) ^98-101^.

**Fig. 8:**
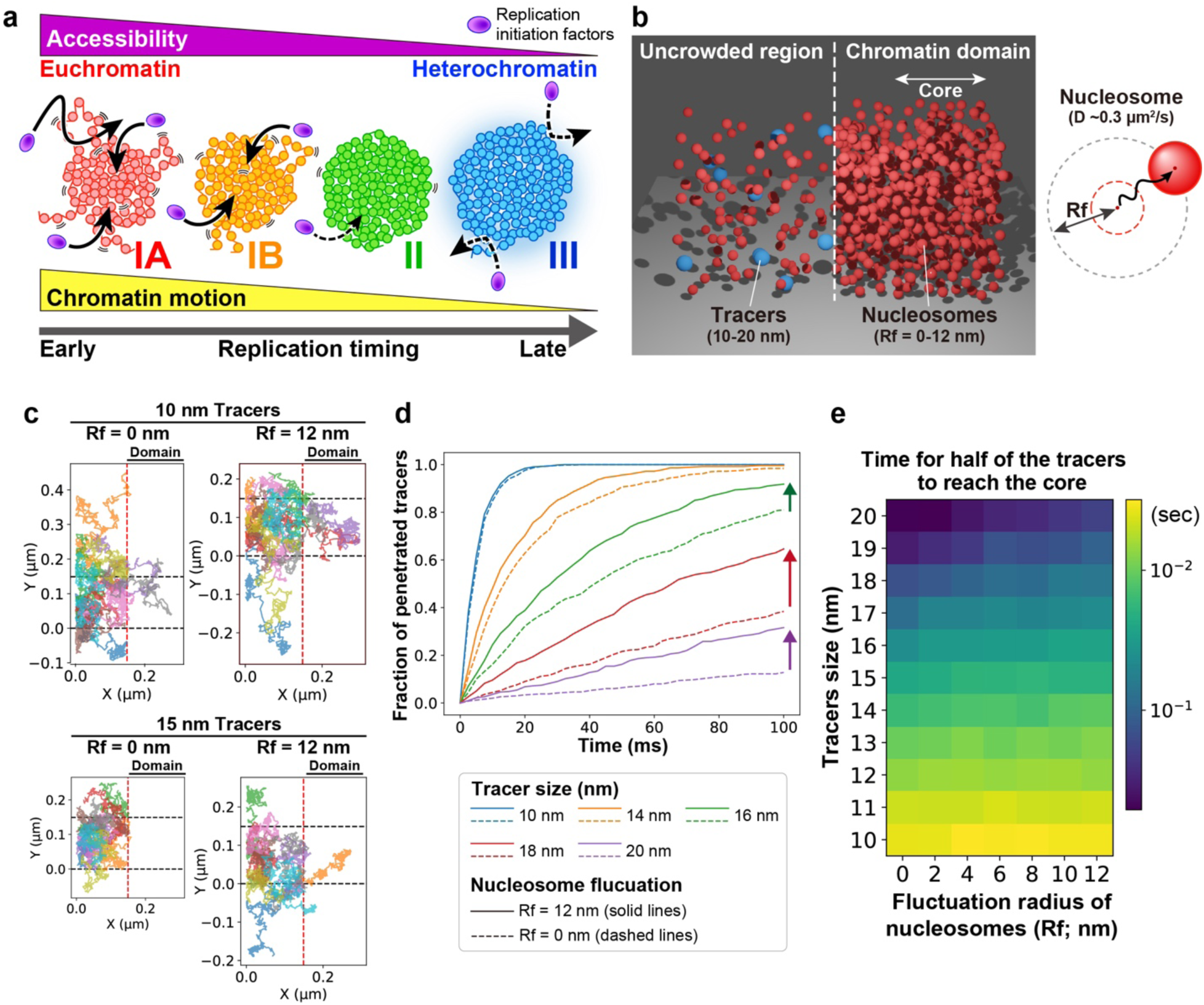
Local nucleosome motion facilitates accessibility of condensed chromatin domains. **a**, The landscape of local nucleosome motion from euchromatin (early) to heterochromatin (late) replication can govern the chromatin accessibility to rate-limiting replication initiation factors, which determines the probability of origin firing. **b**, Left: schematic of the reconstructed chromatin environment *in silico* by the Monte Carlo simulation. The model diffusion proteins (tracer; blue) are initially put in the left uncrowded space. The model nucleosomes (red) are placed on the right (“chromatin domain”) with a defined nucleosome fluctuation radius (Rf) ^104,106^. Right: scheme for fluctuation radius (Rf) of the model nucleosomes. **c**, Representative (n = 10) trajectories of the tracers during 1 ms of the simulation. The tracer diameters and the nucleosome Rfs are shown on the top. Red dashed lines, the boundary of chromatin domains; black dashed lines, the periodic boundary of the reconstructed environment. **d**, Fractions of the tracers with various diameters that penetrated the core region of the chromatin domain with various nucleosome Rfs. An experimentally estimated diffusion coefficient of nucleosomes (*D* ∼0.3 µm^2^/s) was used (for details, see Fig. S12). Note that the penetration rates of large tracers are facilitated by nucleosome fluctuation (arrows). **e**, The time it takes for half of the tracers to reach the core of the chromatin domain. Simulations with various tracer diameters and nucleosome Rfs were conducted.

To tackle this hypothesis, we reconstructed the chromatin environment *in silico* using the Metropolis Monte Carlo method ^102, 103^ to simulate the accessibility of fluctuating chromatin domains. The model nucleosomes were placed on one side of the simulation space (“chromatin domain”) at 0.5 mM (Fig. 8b). The nucleosomes in the domain are mobile with defined radius of fluctuation (Rf) ^104-106^. We put model diffusing proteins (tracers) in the uncrowded space and examined how many diffusing proteins access the core region of the chromatin domain (see Methods for details). This simple model might not necessarily reflect the chromatin state in live cells because the mimic nucleosome spheres are not connected by linker DNA. However, they are still useful for testing our hypothesis.

Although small tracers (< 10 nm) freely moved in the crowded chromatin domains regardless of nucleosome Rf (Fig. 8c; Movie S27-28), large tracers (> 10 nm), which correspond to replication initiation complexes/factors ^107-109^, could not reach the core region of the chromatin domains with immobile nucleosomes (Fig. 8c; Movie S29), consistent with previous reports that local nucleosome fluctuation increased the accessibility to large proteins (Movie S30)^104-106^. Interestingly, the accessibility depends both on the Rf of nucleosomes and the tracer diameter (Fig. 8d). This suggests that the nucleosome motion affects the kinetics of the tracer (i.e., model protein) penetration. Indeed, the larger the nucleosome fluctuates, the faster the model protein can penetrate the chromatin domain (Fig. 8d-e). In particular, the model proteins with a 20 nm diameter, which correspond to replication initiation complexes/factors ^107-109^, had a prominent effect and penetrated only the chromatin domain with nucleosome fluctuation (Fig. 8d-e).

### Local chromatin motion governs DNA replication timing

Finally, we wondered whether the genome-wide landscape of local chromatin motion (Fig. 8a) contributes to the regulation of DNA replication timing. To address this question, we perturbed the genome-wide replication program by knocking down one of the key replication timing regulators, RIF1 ^110, 111^(Fig. S13a-b). Depletion of RIF1 perturbs the replication timing by increasing cell-to-cell heterogeneity of the replication timing ^112^. Consistent with the previous study ^112^, 40% of cells with RIF1 knockdown by siRNA showed an “ambiguous Repli-Histo labeling pattern,” which could not be categorized into the four classes, ensuring effective perturbation of the replication timing (Fig. S13c; Table 1).

For single-nucleosome imaging in RIF1-knockdown cells, we first pre-labeled the existing H3.2-Halo with Rhodamine 110 Direct (R110) (Fig. 9a) so that the total intensity of R110 could identify the S phase stage (or replication timing) in the cell (1^st^ row, Fig. 9b). We then performed Repli-Histo labeling for the newly replicated nucleosomes with dual color (Fig. 9a; Movie S31): a high concentration of TMR to visualize the Repli-Histo labeling patterns and a very low concentration of JF646 to image single nucleosomes (Fig. 9b). After DNA replication, we monitored the nucleosome motion at the following G2 phase (Fig. 9c-d).

**Fig. 9:**
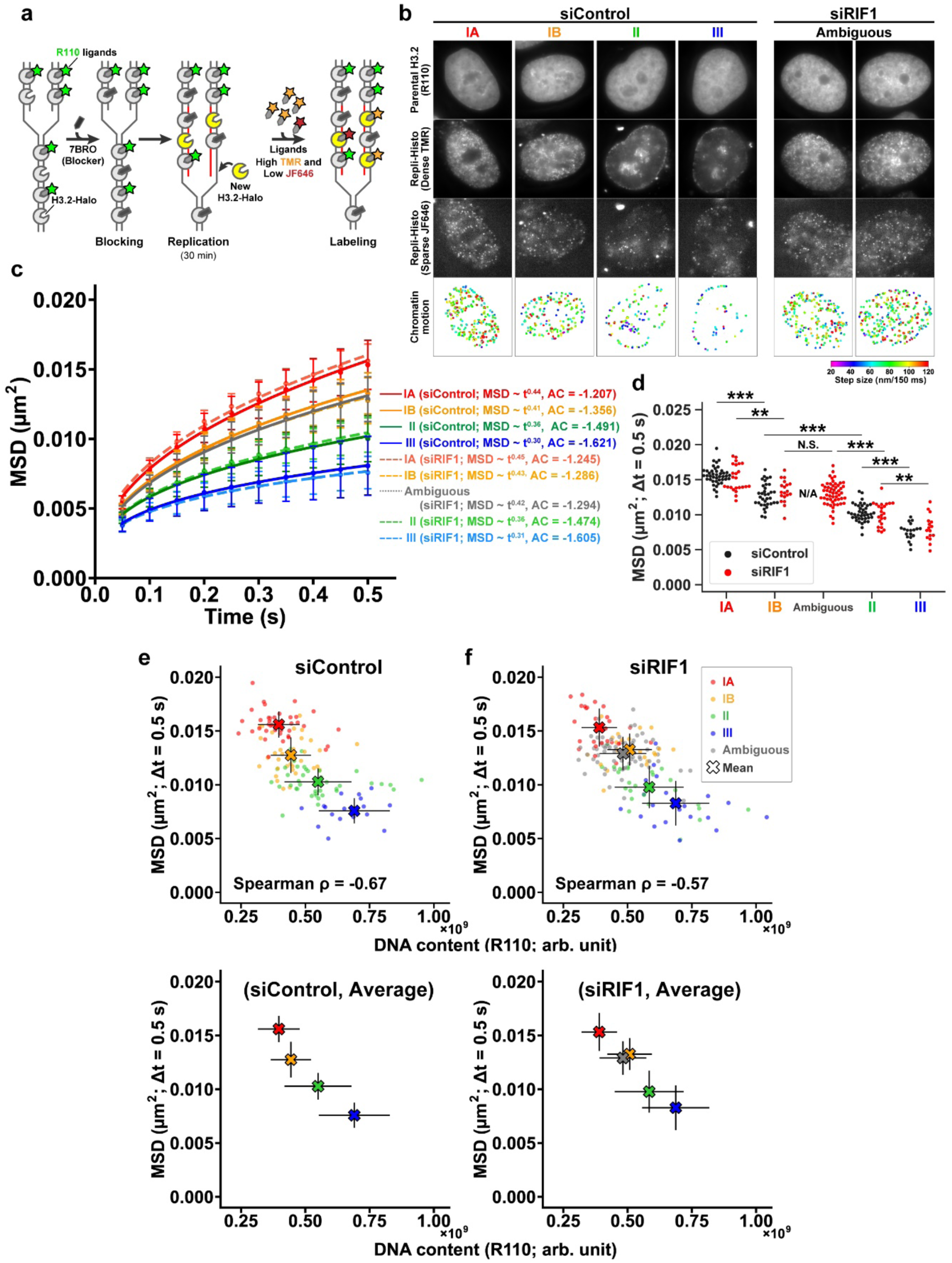
Local chromatin motion governs DNA replication timing. **a**, Schematic of the three-color Repli-Histo labeling for single-nucleosome imaging. Parental H3.2-Halo was labeled with green R110-direct HaloTag ligands. The remaining H3.2-Halo were then blocked with 7BRO, followed by pulse labeling with the two HaloTag ligands (TMR/JF646). **b**, Representative control (siControl) and RIF1-depleted (siRIF1) HeLa cells with the three-color Repli-Histo labeling. 1^st^ row: parental H3.2-Halo labeled with R110, identifying the S phase stage or replication timing; 2^nd^ and 3^rd^ rows: the Repli-Histo labeling patterns (TMR) and corresponding single-nucleosomes (JF646); 4^th^ row: the displacement of single nucleosomes as 2D heatmaps. See Table 1 for the description of the labeling patterns. **c**, MSD plots (± SD among cells) of single nucleosomes in Class IA (red; siControl, n = 43; siRIF1, n = 27), IB (orange; siControl, n = 35; siRIF1, n = 22), II (green; siControl, n = 40; siRIF1, n = 24), and III (blue; siControl, n = 19; siRIF1, n = 23), and ambiguous patterns (grays; iControl, n = 0; siRIF1, n = 72) in siRIF1 (solid lines) or siControl (dashed lines) cells from 0.05 to 0.5 s. MSD exponents and AC values from Fig. S13d–e are also shown in the brackets. **d**, MSD values at 0.5 s in each cell from (**c**). Black, siControl; Red, siRIF1. For siControl, ***: *P* = 5.6 × 10^-10^ for Class IA versus IB, *P* = 1.9 × 10^-8^ for Class IB versus II, and *P* = 1.6 × 10^-8^ for Class II versus III by the two-sided Kolmogorov-Smirnov test. For siRIF1, **: *P* = 0.0012 for Class IA versus IB, *P* = 0.0035 for Class II versus III, and ***: *P* = 5.7 × 10^-10^ for ambiguous patterns versus Class II. **: *P* = 3.2 × 10^-9^ for Class IB versus II. N.S.: *P* = 0.70 for Class IB versus ambiguous patterns, by the two-sided Kolmogorov-Smirnov test. **e**, Correlation between MSD values at t = 0.5 s and R110 signals (i.e., DNA content at the time of Repli-Histo labeling). The bold “X” and bars show each stage’s mean and SD (also shown in the bottom plots). Spearman’s correlation coefficients among all cells are ρ = -0.67 for siControl and ρ = -0.57 for siRIF1.

The control cells with mock depletion exhibited a clear anti-correlation between local nucleosome motion and DNA contents (i.e., S phase stage in the labeling or replication timing) (Fig. 9e; Spearman’s correlation coefficient ρ = -0.67). The earlier replicated regions have a larger nucleosome motion with higher MSD exponents and AC values (Figs. 9c and S13d-e). Strikingly, this anti-correlation was also retained in the RIF1-depleted cells (Fig. 9f; ρ = -0.57), suggesting that genome chromatin is essentially replicated from regions with greater nucleosome motions, even though the genomic replication timing program is perturbed.

Interestingly, RIF1 depletion slightly upregulated the average local nucleosome motion (Fig. S13f), indicating that RIF1 somehow constrains the local nucleosome motions in specific chromatin regions or whole chromatin. Which chromatin regions are constrained by RIF1? It is likely that the Class II chromatin region is constrained by RIF1 because of the following reasons.

First, Class II chromatin was reduced after RIF1 depletion (Fig. S13c). RIF1 seems to constrain the local nucleosome motion of Class II (mid-S replicating) chromatin and maintain its mid-replication (bottom, Fig. 9e-f). Consistently, RIF1 localization is similar to Class II chromatin (2^nd^-3^rd^ rows in Fig. S13a), as reported previously ^110, 111^. Second, chromatin with the ambiguous Repli-Histo patterns upon RIF1 depletion showed a similar nucleosome motion and replication timing to the Class IB chromatin (Fig. 9c-d; ambiguous). Taken together, our results suggest that RIF1 knockdown diminishes the constraints of mid-replicating chromatin, upregulates its nucleosome motion, and facilitates the earlier replication of the regions. Again, this finding supports that genome chromatin is primarily replicated from regions with greater nucleosome motions (Fig. 8a).

## Discussion

To specifically label euchromatin and heterochromatin in living mammalian cells, we have developed Repli-Histo labeling in this study. Repli-Histo labeling allows chromatin visualization of the four regions (Class IA, IB, II, and III; Fig. 4a) historically categorized based on replication timing^60, 61, 62^. We have quantitatively measured nucleosome motion in these four chromatin regions at a high spatiotemporal resolution, which had yet to be conducted. We have demonstrated that the more euchromatic regions have larger local chromatin motion, and the heterochromatic regions are more constrained (Fig. 4b-c). Interestingly, the nucleosomes in the heterochromatin around the nuclear periphery and nucleoli showed similar MSD values (Fig. 4e), suggesting that chromatin replicated at the same timing exhibits similar physical properties, regardless of its intranuclear location. 2-point MSD analyses of two neighboring nucleosome distances suggested the fluidity of their chromatin domains. Euchromatin looks more liquid-like and heterochromatin is more gel-like, presumably due to additional crosslinks in heterochromatin such as HP1 and lamina (Fig. 7d-e) ^21-24, 86^. Since local chromatin motion can facilitate chromatin accessibility to large proteins, greater motion in euchromatin empowers transcription competency while both types of chromatin retain a certain degree of accessibility (Fig. 8a). Our findings provide a new insight into the physical nature of euchromatin and heterochromatin.

Our data also show that chromatin cannot be bimodally classified into euchromatin and heterochromatin. Since the nucleosome motion is increasingly more constrained as the Class progresses from IA through III, chromatin exhibits a rather continuous chromatin-state spectrum. This concept is in good agreement with the Hi-C sub-compartments idea ^20^ and “chromatin color” formed by multiple combinations of the chromatin modifications ^78, 113^.

In contrast to the motion difference between different chromatin classes (IA, IB, II, and III), the chromatin motion in each Class is maintained constant throughout the G1, S, and G2 phases (Figs. 4b, 5c, and 6d). This finding is consistent with our previous genome-wide observation ^54^ showing that the local chromatin motion, on average, remains constant throughout interphase. Since the constant chromatin motion allows cells to conduct housekeeping functions (e.g., transcription and DNA replication) under similar environments throughout interphase, cells can do their routines in each genome chromatin region under a similar condition during interphase. The constant chromatin motion in each chromatin Class also maintains the reactivity preference among the classes, including euchromatin and heterochromatin, during interphase. Our finding that the chromatin motion in each Class is similar between the S phase and G1/G2 phases is consistent with our previous report that the rapid degradation of MCM2 by the AID system (i.e., disassembly of the DNA replisome) ^114^ did not significantly affect local chromatin motion ^54^. This implies that DNA replication complexes, such as the replisome, do not actively induce nucleosome motions, at least on the second time-scale.

Our present study provides a possible physiological role of the chromatin motion landscape in DNA replication timing. We propose that local nucleosome fluctuation is a stochastic factor in replication timing that determines the origin firing efficiency of a given genomic region (Fig. 8a) for the following reasons. First, multiple studies have demonstrated that eukaryotic replication origins fire stochastically with a certain probability^115-117^, supporting the stochastic nature of replication timing ^118-120^. Second, several studies have reported that limited numbers of initiation factor(s), such as SLDs and CDC45, are competed for among origins during the firing process ^99,100^. With greater nucleosome motions, initiation factors bind to the origins more efficiently (Fig. 8d-e). Consistently, almost no overlaps or gaps between the MSD plots in the four Class regions were observed (Fig. 4b-c and 4f-h). On the other hand, the process of DNA replication elongation does not appear to be affected by differences in nucleosome motion between the four Class regions, as CDC45 foci (representing active replication helicase) were clearly detected even in the heterochromatic Class III chromatin (Fig. 6c).

Then, if we upregulate nucleosome fluctuation of a certain region, does the region replicate earlier in the S phase? Yes. First, as we showed in this study, the knockdown of RIF1 seems to increase nucleosome motion in Class II regions (mid-S replicating) and facilitates earlier replication of the regions (bottom, Fig. 9e-f). Second, HDAC inhibitor treatment, which upregulates nucleosome motions globally ^8, 12^, shifts the replication timing of chromocenter heterochromatin earlier ^121^. Numerous studies have reported that many factors, including trans-acting factors such as RIF1, histone modifications, and spatial organization ^112, 122-127^, can affect the replication timing. In addition to the RIF1-protein phosphatase 1 (PP1) axis counteracting the DDK-mediated origin activation ^128-130^, these factors may govern nucleosome fluctuation and chromatin accessibility to the rate-limiting replication initiation factors for origin firing (Fig. 8a)^98-101^.

Finally, it would be informative to discuss some attractive applications for Repli-Histo labeling technology. In addition to super-resolution live-cell imaging of euchromatin and heterochromatin regions using PALM ^131^ or 3D-SIM ^10^, a promising application is the use of correlative light and electron microscopy (CLEM). CLEM combines the global contrast and high resolution of EM with the molecular specificity of fluorescence microscopy ^132, 133^. Integration of these techniques now offers a closer match in resolution between the two modalities, light and electron microscopy, allowing specific euchromatin and heterochromatin to be visualized at nanoscale resolution in the context of the crowded intracellular environment ^134^.

## Methods

### Plasmid construction

Construction of pX330-H3.2-gRNA, pX330-CDC45-HaloTag, and pX330-H3.1-gRNA were carried out as described previously ^135^. Gene-specific guide RNA sequences for human histone H3.2 (*H3C14/H3C15*), CDC45 (*CDC45*), and mouse histone H3.1 (*Hist1h3a*) genes were designed using the CRISPR design website (CHOPCHOP ^136^) and inserted into the *Bbs*I cloning site in pX330 (#42230, Addgene), as previously described ^67^. The guide RNA sequence was 5’-CCGCCGCATCCGTGGAGAGC-3’ for H3.2, 5’-GAACCCCTGTAACTCACCCT-3’ for CDC45, and 5’-ACAATTACGCCCTCTCCCCG-3’ for mouse H3.1.

Construction of donor plasmids pGEM-H3.2-HaloTag, pGEM-CDC45-HaloTag, and pGEM-H3.1-HaloTag was carried out as follows. The HaloTag coding sequence was PCR amplified using KOD FX (KFX-101; Toyobo) from the pFC14A HaloTag CMV Flexi Vector (G965A; Promega). The homology arms for human H3.2, CDC45, and mouse H3.1 flanked from the HaloTag sequence (∼700 bp each) were PCR amplified from HeLaS3 and 10T1/2 genomic DNA, which was isolated using a Wizard Genomic DNA Purification kit (Promega). The homology arms and the HaloTag fragment were inserted between the *Eco*RI and *Sal*I sites of the pGEM-T (Easy) vector (A137A; Promega) using In-Fusion HD Cloning Kit (639648; Clontech).

### Cell lines

HeLaS3 ^137^, RPE-1, and 10T1/2 ^88, 89^(a gift from Dr. Michael J. Hendzel at University of Alberta) cells were cultured at 37 °C in 5% CO_2_ in Dulbecco’s Modified Eagle’s medium (DMEM) (D5796-500ML; Sigma-Aldrich) supplemented with 10% fetal bovine serum (F7524; Sigma-Aldrich). HCT116 cells were gifted from Prof. Masato Kanemaki (National Institute of Genetics, Mishima, Japan) and were cultured at 37 °C in 5% CO_2_ in McCoy’s 5A medium (SH30200.01; HyClone) supplemented with 10% fetal bovine serum. The HeLaS3 cell line stably expressing H2B-Halo was previously established as described in ^8^.

To establish HeLaS3, RPE-1, and HCT116 cells expressing endogenous human H3.2 or CDC45 tagged with HaloTag, a CRISPR/Cas9 genome editing system was used. The constructed plasmid pGEM-H3.2-HaloTag or pGEM-CDC45-HaloTag, as well as the Cas9/sgRNA expression plasmid pX330-H3.2-gRNA or pX330-CDC45-gRNA, were electroporated into the HeLaS3, RPE-1, or HCT116 cells. The electroporation was performed using the Neon Transfection System (Invitrogen). After one to two weeks, the cells expressing H3.2-HaloTag or CDC45-HaloTag were fluorescently labeled with 50 nM HaloTag TMR ligand (G8252; Promega) or JF646 ligand (GA1120; Promega) overnight at 37 °C in 5% CO_2_. The HaloTag TMR ligand-positive cells were then collected using flow cytometry (SH800S; Sony Biotechnology). The collected cells were subcloned, and the cells stably expressing H3.2-HaloTag or CDC45-HaloTag were selected. The selected clones were rechecked with PCR for correct insertion of the HaloTag coding sequence downstream of the endogenous H3.2/CDC45 loci. To establish 10T1/2 cells expressing endogenous mouse H3.1 tagged with HaloTag, a similar CRISPR/Cas9 genome editing was performed as follows. The constructed plasmids pGEM-H3.1-HaloTag and pX330-H3.1-gRNA were electroporated into the 10T1/2 cells using the Neon Transfection System (Invitrogen). The 10T1/2 clone stably expressing mouse H3.1-HaloTag was selected as described above. The selected clone was rechecked with PCR for correct insertion of the HaloTag coding sequence downstream of the endogenous H3.1 locus.

### RNA interference

siRNA transfection was performed using Lipofectamine RNAiMAX (13778-075; Thermo Fisher Scientific) according to the manufacturer’s instructions. The transfected cells were used for subsequent studies 48 h after transfection. The siRNA oligonucleotide targeting RIF1 sequence (Invitrogen; Sense: 5’-GAAUGAGCCCCUAGGGAAATT-3’) ^138^ was used. A control oligonucleotide (4390843; Thermo Fisher Scientific) was used for mock depletion.

### Western blotting

Cells were lysed in Laemmli sample buffer supplemented with 10% 2-mercaptoethanol (133-1457; Wako) and incubated at 95 °C for 5 min to denature proteins. The cell lysates, equivalent to 1 × 10^5^ cells per well, were subjected to SDS–polyacrylamide gel electrophoresis (12.5% for histone detection and 8% for CDC45 detection). For western blotting, the fractionated proteins in the gel were transferred to a polyvinylidene difluoride (PVDF) membrane (IPVH00010; Millipore) by a semi-dry blotter (BE-320; BIO CRAFT). After blocking with 5% skim milk (190-12865; Fujifilm Wako), the membrane-bound proteins were probed by the rabbit anti–H3 (ab1791; Abcam, 1:25,000), mouse anti-H3.1/H3.2 (CEC-006; Cosmo Bio, 1:1,000), mouse anti–HaloTag (G9281; Promega, 1:1,000), or rabbit anti-CDC45 (11881S; Cell Signaling Technology, 1:1,000), followed by the appropriate secondary antibody: anti-rabbit IgG (170-6515; Bio-Rad, 1:5,000) or anti-mouse IgG (170-6516; Bio-Rad, 1:5,000) horse radish peroxidase-conjugated goat antibody. Bands were detected by chemiluminescence reactions (WBKLS0100; Millipore) and images were acquired with the EZ-Capture MG (AE-9300H-CSP; ATTO).

### Biochemical fractionation of nuclei from cells expressing H3.2-HaloTag

Nuclei were isolated from HeLa cells expressing endogenous H3.2-HaloTag as described previously ^54^. Briefly, collected cells were suspended in nuclei isolation buffer [3.75 mM Tris-HCl (pH 7.5), 20 mM KCl, 0.5 mM EDTA, 0.05 mM spermine, 0.125 mM spermidine, aprotinin (1 µg/ml) (T010A; TaKaRa), and 0.1 mM phenylmethylsulfonyl fluoride (PMSF) (P7626-1G; Sigma-Aldrich)] and centrifuged at 1936 × *g* for 7 min at room temperature. The cell pellets were resuspended in nuclei isolation buffer and again centrifuged at 1936 × *g* for 7 min at room temperature. Subsequent steps were performed at 4 °C unless otherwise noted. Cell pellets were resuspended in nuclei isolation buffer containing 0.025% Empigen (45165-50ML, Sigma-Aldrich) (nuclei isolation buffer+) and homogenized immediately with 10 downward strokes of a tight Dounce pestle (357546; Wheaton). The cell lysates were centrifuged at 4336 × *g* for 5 min. The nuclear pellets were washed in nuclei isolation buffer+. The nuclei were incubated on ice for 15 min in the following buffers containing various concentrations of salt: HE [10 mM HEPES-NaOH (pH 7.5), 1 mM EDTA, and 0.1 mM PMSF], HE + 100 mM NaCl, HE + 500 mM NaCl, HE + 1 M NaCl, and HE + 2 M NaCl. After each buffer incubation with increasing concentrations of salt, centrifugation was performed to separate the nuclear suspensions into supernatant and pellet fractions. The proteins in the supernatant fractions were precipitated by using 17% trichloroacetic acid (208-08081; Wako) and cold acetone. Both pellets were suspended in a Laemmli sample buffer and subjected to 12.5% SDS-PAGE, followed by Coomassie brilliant blue (031-17922; Wako) staining and western blotting using rabbit anti-H3 (ab1791; Abcam) and rabbit anti-HaloTag (G9281; Promega) antibodies to confirm H3.2-HaloTag expression.

### HaloTag-labeling

To check the distribution of the expressed HaloTag-fused proteins, H2B-Halo, H3.2-Halo, or CDC45-Halo were fluorescently labeled with 50 nM HaloTag TMR or JF646 ligands overnight at 37 °C in 5% CO_2_. Subsequently, the cells were fixed with 1.85% formaldehyde (064–00406; Wako) for 15 min and then permeabilized with 0.5% Triton X-100 (T-9284; Sigma Aldrich) for 5 min and stained with 0.5 µg/ml DAPI (10236276001; Roche) for 5 min before embedded in PPDI [20 mM HEPES (pH 7.4), 1 mM MgCl_2_, 100 mM KCl, 78% glycerol, and 1 mg/ml paraphenylene diamine (PPDI, 695106-1G; Sigma-Aldrich)].

For single-nucleosome imaging, H2B-Halo, H3.2-Halo, and H3.1-Halo molecules were fluorescently labeled with 50 pM HaloTag TMR ligand for 20 min at 37 °C in 5% CO_2_. The cells were washed with Hank’s balanced salt solution (1× HBSS, H1387; Sigma-Aldrich) three times and then incubated in a phenol red-free DMEM (21063–029; Thermo Fisher Scientific) or McCoy’s 5A (1-18F23-1, BioConcept) for over 2 h before live-cell imaging.

For single-CDC45 imaging, CDC45-Halo in asynchronous HeLa cells was fluorescently labeled with 5 nM HaloTag JF646 ligand for 30 min at 37 °C in 5% CO_2_, followed by labeling with 10 nM HaloTag TMR ligand overnight. The cells were then washed with 1× HBSS three times and then incubated in a medium without phenol red for more than 4 h before live-cell imaging. We observed the S phase cells, which formed CDC45-Halo foci corresponding to Class IA, IB, II, and III (Movies S18-21).

### Repli-Histo labeling

The replication-dependent histone labeling (Repli-Histo labeling) was performed as follows. First, asynchronous cells expressing endogenous H3.2-HaloTag were incubated with the medium supplemented with 10 µM 7-bromo-1-heptanol (7BRO, B1852; Tokyo Chemical Industry) ^72^ for 60 min at 37 °C in 5% CO_2_. The cells were then washed three times with medium without 7BRO. The Repli-Histo labeling with 10T1/2 cells expressing mouse H3.1-Halo was performed similarly.

For fixed cell imaging, the cells were exposed to the medium supplemented with 100 nM HaloTag TMR ligand and 10 µM 5-ethynyl-2′-deoxyuridine (EdU) (C10337; Thermo Fisher Scientific) for 30 min immediately after washing out 7BRO. For negative control, 100 µg/ml cycloheximide (037-20991; Fujifilm Wako) was also added to inhibit the new H3.2-Halo synthesis. The cells were then fixed and permeabilized as described above. Subsequently, EdU was labeled with Alexa Fluor 488 or 647 azide using Click-iT EdU Cell Proliferation Kit (C10337; Thermo Fisher Scientific) and stained with 0.5 µg/ml DAPI for 5 min before being embedded in PPDI as described above.

For Fig. S5b, the soluble H3.2-Halo was pre-extracted with 0.5% Triton X-100 in ice-cold CSK buffer [10 mM HEPES-KOH (pH 7.5), 300 mM sucrose, 100 mM NaCl, 3 mM MgCl_2_, 1x cOmplete Mini protease inhibitor cocktail (11836153001; Roche) and 0.1 mM phenylmethylsulfonyl fluoride (PMSF, P7626; Sigma-Aldrich)] for 2 min and washed with the same ice-cold buffer without Triton X-100 (twice for 3 min)^137^. Subsequent operations were carried out as described above.

For single nucleosome imaging, the cells were exposed to the mixture of 2 nM HaloTag TMR ligand and 5 nM HaloTag JF646 ligand for 30 min. The cells were then washed three times with a medium without phenol red, followed by incubation in a medium without phenol red for 1 h before live-cell imaging.

For Fig. 9, the parental H3.2-HaloTag was labeled with 10 nM HaloTag R110Direct ligands (G3221; Promega) overnight before the blocking with 7BRO. The following Repli-Histo labeling for single-nucleosome imaging was carried out as described above.

To characterize the replication timing of the labeled regions, the labeled patterns (historically called replication foci) of the Repli-Histo labeling were manually categorized into the four classes (IA, IB, II, and III) based on previous studies ^60,61^ (See Table 1). Alternatively, the replication timing of the Repli-Histo-labeled region was estimated from the total nuclear pixel intensities (Arbitrary units: arb. unit) of R110Direct-labeled parental H3.2-HaloTag (Fig. 8).

### Cell cycle synchronization

To obtain HeLa cells arrested in the late G1 phase, asynchronous HeLa cells were first labeled with the Repli-Histo labeling, chased for 7 h, and then treated with 400 µM L-mimosine (M0253-25MG; Sigma) for 17 h ^91^. For cell cycle synchronization of HeLa cells in the early/late S phase, double thymidine block and release were performed as follows. HeLa cells (1.0 × 10^5^ cells) were plated with 3 mM thymidine (T9250, Sigma-Aldrich) for 18 h, released into a thymidine-free medium for 9 h, and again treated with 3 mM thymidine for 17 h for the G1/S arrest. The arrested cells were then released into a thymidine-free medium for 1 h (Class IA), 3 h (Class IB), 5 h (Class II), or 6 h (Class III).

### HaloTag pull-down sequencing

Cell nuclei were isolated as previously described ^50, 139, 140^ with minor modifications. First, we arrested HeLa cells at G1/S by double thymidine block treatment as described above. The cells were next released into the S phase for 1 h or 5 h, followed by 10 nM 7BRO treatment for 30 min. After a three-time wash of 7BRO with PBS, the cells were incubated with a 7BRO-free medium for 1 h for the deposition of newly synthesized H3.2-Halo. The cells were then collected and washed with nuclei isolation buffer [3.75 mM Tris-HCl (pH 7.5), 20 mM KCl, 0.5 mM EDTA, 0.05 mM spermine, 0.125 mM spermidine, 0.1% Trasylol (14716; Cayman Chemical), and 0.1 mM phenylmethylsulfonyl fluoride (PMSF, P7626-1G; Sigma-Aldrich)] by two cycles of centrifugal spins at 193 × *g* for 7 min at 23 °C. The pellets were then resuspended in nuclei isolation buffer containing 0.05% Empigen BB detergent (45165-50ML; Sigma-Aldrich) (nuclei isolation buffer+) and immediately homogenized with 10 downward strokes using a tight Dounce pestle. After a 5-min centrifugation of the cell lysates at 440 × *g*, the pellets were washed and resuspended in nuclei isolation buffer+. After another centrifugation at 440 × *g* at 4 °C for 5 min, the supernatant was removed and resuspended in 10 ml of nuclei isolation buffer+. This step was repeated four times before the samples were ready for nucleosome purification.

Chromatin purification was carried out as described by ^50, 134, 141^, with some modifications. The nuclei (equivalent to ∼2 mg of DNA) in nuclei isolation buffer [10 mM Tris–HCl (pH 7.5), 1.5 mM MgCl_2_, 1.0 mM CaCl_2_, 0.25 M sucrose, and 0.1 mM PMSF] were digested with 50 U of micrococcal nuclease (LS004797; Worthington) at 30 °C for 1 h. In this digestion step, 5 µM HaloTag PEG-biotin ligand (G859A; Promega) or an equivalent amount of DMSO was added. After centrifugation at 440 × *g* at 4 °C for 5 min, the nuclei were lysed with lysis buffer [10 mM Tris–HCl (pH 8.0), 5 mM EDTA, and 0.1 mM PMSF] on ice for 5 min. The lysate was dialyzed against dialysis buffer [10 mM HEPES-NaOH (pH 7.5), 0.1 mM EDTA, and 0.1 mM PMSF] at 4 °C overnight using Slide-A-Lyzer (66380; Thermo Fisher Scientific). The dialyzed lysate was centrifuged at 20,400 × *g* at 4 °C for 10 min. The supernatant was recovered and used as the purified nucleosome fraction (mainly mono nucleosomes). To verify complete MNase digestion, DNA was purified, electrophoresed on a 1.5% agarose gel, and visualized by staining with ethidium bromide (Fig. S2b).

The biotinylated or untreated nucleosome fraction (28 µg) was diluted with an equal volume of 2× binding buffer [10 mM HEPES-NaOH (pH 7.5), 200 mM NaCl, 2 mM DTT, and cOmplete™ EDTA-free protease inhibitor cocktail (11873580001; Roche)]. As shown previously ^142^, the chromatin solution was added to streptavidin-FG beads (TAS8848 N1170; Tamagawa-Seiki) pre-equilibrated with coating buffer [PBS, 2.5% BSA, 0.05% Tween 20] for 1 h. The mixtures were incubated at 5 °C for 2 h on a shaker. The beads were collected with a magnetic rack and subjected to the washing procedure as described by Gatto et al. ^73^. Briefly, the beads were washed once with 1 ml of cold 1× binding buffer [10 mM HEPES-NaOH (pH 7.5), 100 mM NaCl, 1 mM DTT], twice with 1 ml of cold wash buffer 1 [HEPES-NaOH (pH 7.5), 140 mM NaCl, 1% Triton X-100, 0.5% Nonidet P-40 substitute (NP40, 11332473001; Roche), 0.1% SDS], twice with 1 ml of cold wash buffer 2 [HEPES-NaOH (pH 7.5), 360 mM NaCl, 1% Triton X-100, 0.5% NP40, 0.1% SDS], twice with 1 ml of cold wash buffer 3 [HEPES-NaOH (pH 7.5), 250 mM LiCl, 0.5% Triton X-100, 0.5% NP40] and once with TE buffer [10 mM Tris–HCl (pH 8.0), 1 mM EDTA]. Beads were resuspended in 50 μl TE buffer.

For DNA extraction, 40 µl of beads underwent RNase digestion [4 μl RNase A (10109169001; Roche), 1 mg/ml, 30 min at 37 °C] and subsequent proteinase K digestion in the presence of SDS [4 μl of 20 mg/ml proteinase K (169-21041; Wako) and 10 μl of 10% SDS, at 1,200 rpm and 50 °C on shaker for 1 h]. Beads were removed by magnetic separation and DNA extracted using AMPure XP beads (A63880; Beckman Coulter) according to the manufacturer’s instructions, and eluted in Milli-Q water. Total DNA content of each sample was measured using Qubit system (Q32851; Thermo Fishe Scientific), and the quality of DNA samples was assessed by an Agilent 2100 Bioanalyzer using an Agilent High Sensitivity DNA kit (5067-4626; Agilent). cDNA libraries were synthesized by the ThruPLEX DNA-seq Kit (R400675; Takara). The size distributions of the libraries were checked by an Agilent 2100 Bioanalyzer using an Agilent High Sensitivity DNA kit. The pooled amplicon library was sequenced with paired-end 2 × 100 bp reads on the Illumina NovaSeq 6000 platform.

For protein analysis of the HaloTag pull-down fraction (Fig. S2a), the remaining 10 µl of beads were resuspended with an equal volume of 2× Laemmli sample buffer ^143^ supplemented with 10% 2-mercaptoethanol (133-1457; Wako) and incubated at 95 °C for 10 min. Beads were removed by centrifugation at 10,000× *g* for 3 min. The Input sample and the pull-down samples from the biotinylated and untreated nucleosome (negative control) were separated using SDS–polyacrylamide gel electrophoresis and were subjected to western blotting with anti-HaloTag antibody (G9211, Promega).

For data analysis of purified H3.2-Halo labeled nucleosomal DNA, the nf-core ^144, 145^ ChIP-seq pipeline (nfcore/chipseq: version 2.0.0) was utilized, employing a Docker configuration profile and executed using the default parameters. The human genome hg19, retrieved from Illumina’s iGenomes, was used for the reference genome. Peaks were detected using the broad mode of MACS2 ^146^. To evaluate the overlap between the H3.2-HaloTag labeled regions and contact domains (i.e., Hi-C A/B-compartment and histone modification patterns), we compared the H3.2-HaloTag regions with previously published data as described in ^12, 50^. Compartment annotations on the HeLa cell genome are SNIPER predictions obtained from ^75^. Peak lists on hg19 genome defined by the ChIP-seq analysis for the histone modifications were obtained from the ENCODE portal ^74 147^ with the following identifiers: H3K4me2 (ENCFF108DAJ), H3K4me3 (ENCFF447CLK), H3K9ac (ENCFF723WDR), H3K27ac (ENCFF144EOJ), H3K79me2 (ENCFF916VLX), H3K36me3 (ENCFF001SVY), H3K4me1 (ENCFF162RSB), and H3K27me3 (ENCFF252BLX). Peak lists on hg19 genome defined by ChIP-seq analysis for the lamin B1 were obtained from ^79^ (GEO accession number: GSE57149).

### Indirect immunofluorescence

Immunostaining was performed as described previously ^104^, and all processes were performed at room temperature. Cells were fixed in 1.85% FA in PBS for 15 min and then treated with 50 mM glycine in HMK [20 mM HEPES (pH 7.5) with 1 mM MgCl_2_ and 100 mM KCl] for 5 min and permeabilized with 0.5% Triton X-100 in HMK for 5 min. After washing twice with HMK for 5 min, the cells were incubated with 10% normal goat serum (NGS, 143-06561; Wako) in HMK for 30 min. The cells were incubated with the diluted primary antibody, rabbit anti–RIF1 (1:1,000, A300-568A; Bethyl Laboratories), in 1% NGS in HMK for 1 h. After being washed with HMK four times, the cells were incubated with the diluted secondary antibody, goat anti–rabbit IgG Alexa Fluor 488 (1:500, A11006; Thermo Fisher Scientific), in 1% NGS in HMK for 1 h followed by a wash with HMK four times. For DNA staining in fixed cells, 0.5 μg/ml DAPI was added to the cells for 5 min followed by washing with HMK. The stained cells were embedded in PPDI.

### Fluorescent microscopy on fixed samples

Image stacks of fixed cells were acquired by using Delta Vision Ultra (GE Healthcare) with an Olympus UPLXAPO 100× (NA 1.45) or UPLXAPO 60x (NA 1.42) objective. Optical sections at a thickness of 0.2 µm were imaged. For Fig. S1d, images were deconvolved using SoftWoRx software (Applied Precision / GE Healthcare). For interphase nuclei, the best-focused section to the middle of nuclei was extracted after correcting the chromatic aberration. Image analysis was performed using Fiji/ImageJ ^148, 149^.

To quantify the intranuclear signal intensity of Repli-Histo labeling, DAPI-stained nuclear regions were segmented based on Huang’s fuzzy thresholding method with Fiji/ImageJ, and the total nuclear pixel intensities (arb. unit) and mean nuclear intensities (arb. unit) for DAPI and all other signals of interest were measured. DNA content was plotted on the x-axis using total DAPI intensities. The cell cycle profiles in Fig. 5a were estimated by the Dean-Jett-Fox method ^150^. At least 100-200 random interphase cells were imaged in all scatter plots. The colocalization of the EdU and the Repli-Histo signals was evaluated by the pixel-wise Pearson’s correlation coefficient between each channel in the nuclear regions.

### Single-nucleosome imaging microscopy

Established cell lines were cultured on poly-L-lysine (P1524-500MG; Sigma-Aldrich) coated glass-based dishes (3970-035; Iwaki). H2B-HaloTag and H3.2-HaloTag molecules were fluorescently labeled with 50 pM HaloTag JF646 ligand, or Repli-Histo labeled as described above.

Single nucleosome imaging was performed as described previously ^12, 47, 50, 54, 89^. To maintain cell culture conditions (37 °C, 5% CO_2_, and humidity) under the microscope, a live-cell chamber and a digital gas mixer (STXG-WSKMX-SET; Tokai Hit) were used. Single nucleosomes were observed by using an inverted Nikon Eclipse Ti2 microscope, an ILE 400 lase combiner (Andor) with 100-mW 561-nm and 140-mW 637-nm laser systems, and a sCMOS ORCA-Fusion BT camera (C15440-20UP, Hamamatsu Photonics). Fluorescently labeled histones in living cells were excited by the 561/637-nm laser through an objective lens (100× PlanApo TIRF, NA 1.49; Nikon) and detected at 582-626/676-786 nm. An oblique illumination system with the TIRF unit (Nikon) was used to excite the labeled histone molecules within a limited thin area in the cell nucleus and reduce background noise. R110direct-labeled H3.2-Halo was imaged with LED epi-illumination (X-Cite XYLIS; Excelitas Technologies) and detected at 503–547 nm. Sequential image frames were acquired using NIS elements software (AR v5.30.03 64bit, Nikon) at a frame rate of 50 ms under continuous illumination. The frame rate and exposure time were controlled using a NI-DAQ board (PCI3-6353; National Instruments). Dual-color imaging was performed through a beam splitter W-VIEW GEMINI (Hamamatsu Photonics) detected at 573-613 nm (TMR) and 662-690 nm (JF646).

### Chemical treatment

For single-nucleosome imaging on chemically fixed cells, cells grown on poly–L–lysine coated glass-based dishes were incubated in 2% formaldehyde in 1×HBSS at 37 °C for 15 min and washed with 1×HBSS three times. The following single-nucleosome imaging was performed as described above.

### Single-nucleosome tracking analysis

The methods for image processing, single-molecule tracking, and single-nucleosome movement analysis were described previously ^8^. The background noise signals in the acquired sequential TIFF images were subtracted with the rolling ball background subtraction (rolling ball radius: 50 pixels) using Fiji/ImageJ. The nuclear regions in the images were manually extracted. Following this step, the diffraction-limited fluorescent dots in each image were fitted to a Gaussian function to obtain the center of the distribution, and its trajectory was tracked with u-track (MATLAB package ^84^). To ascertain the position determination accuracy of the H3.2-HaloTag-nucleosomes in FA-fixed cells, distributions of nucleosome displacements from the centroids of the trajectories in the x-and y-planes (n = 11 or 12 molecules) were fitted to Gaussian functions. The calculated position determination accuracy in each experiment was described in Figs. S4b, S11b, and S12b.

For single-nucleosome movement analysis, the displacement and MSD of the fluorescent dots were calculated based on their trajectory using a Python script. The originally calculated MSD was in 2D. To obtain the 3D value, the 2D value was multiplied by 1.5 (4 to 6 Dt). Statistical analyses of the obtained single-nucleosome MSD captured through each histone protein were performed using Python.

### Two-point MSD analysis

For dual-color single nucleosome imaging, the HeLa cells were first arrested at G1/S with a double thymidine block. 1 h or 5 h after the cells were released into a thymidine-free medium, the cells were incubated with 10 µM 7BRO for 60 min at 37 °C in 5% CO_2_ followed by a wash with medium without 7BRO three times. Subsequently, the cells were labeled with the mixture of 500 pM HaloTag TMR ligand and 10 nM JF646 ligand for 10-15 min.

Image processing, dot detection, and tracking of the dual-color single nucleosome tracking images were carried out as described above. The calculated position determination accuracy through the beam splitter was 11.0 nm (TMR particles) and 10.4 nm (JF646 particles) (See Fig. S11b). Positions of spots labeled by TMR and JF646 colors were corrected by second-order polynomial transformation. Parameters of the transformation were estimated based on the 0.1-μm TetraSpeck bead (T7279; Thermo Fisher Scientific) imaging and calculations were performed using the Python library (Scikit image; ^151^). To analyze the movements of two nucleosomes labeled with two colors (TMR and JF646), we picked closely located (< 150 nm) pairs of TMR and JF646 trajectories from each image movie. Around 60 to 150 pairs of the dots were obtained from a single experiment. The two-point MSD of the fluorescent dots was calculated as described in ^12, 97^. In our single nucleosome tracking, trajectories of TMR- and JF646-dots on the XY plane, 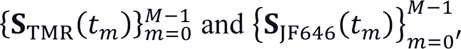 were simultaneously acquired, where the time interval was Δ*t* = 0.05 *s* and *t_m_* = *m* Δ*t* (*m* = 0,1,2, ⋯, *M* − 1). To evaluate dynamic fluctuations between two points, we dealt with the relative vector between two trajectories, ***Q***(*t_m_*) = **S**_TMR_(*t_m_*) – **S**_JF646_(*t_m_*). Then, we calculated the two-point MSD for the lag time *t_n_* = *n* Δ*t* by

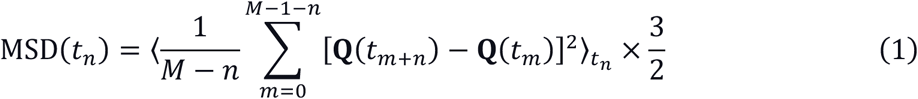

where 〈·〉*_t_n__* represents the ensemble average for trajectories at the lag time *t_n_* and the coefficient 3/2 is a correction factor for conversion from 2D to 3D values.

### Segmentation of the Class III chromatin regions

To extract the single nucleosome trajectories colocalized with late-S replication foci (Fig. S6a), we segmented the late-S replicated chromatin regions using dense TMR-labeled images. First, 10-frame (0.5 s) averaged TMR images were processed with an ImageJ band-pass filter function with the filtering of large structures down to 30 pixels and small structures up to 5 pixels. The filtered images were then binarized using Otsu’s method. The 2-pixel-dilated binary masks were defined as the late-S replication foci region. The detected JF646 dots located inside the foci regions were extracted based on the centroids of their trajectories (i.e. time average of xy-coordinates) as described previously ^53^. The categorized trajectories were used in subsequent analyses.

### Angle distribution analysis

Angle distribution analysis was performed as described in ^54^. For the tracked consecutive points {(x_0_, y_0_), (x_1_, y_1_), ⋯, (x_n_, y_n_), ⋯} of a single nucleosome on the xy-plane, we converted the data into a set of displacement vectors, Δ***r***_n_ = (x_n + 1_ − x_n_, y_n + 1_ − y_n_)^t^. Then, we calculated the angle between two vectors Δ***r***_n_ and Δ***r***_n + 1_. We carried out this procedure for all the points of each trajectory in our experiments. We plotted the normalized polar histogram by our Python program ^53^. The angle distribution was normalized by 2π, and the values correspond to the probability density.

### Monte Carlo simulation of nucleosomes and model proteins

We performed a Metropolis Monte Carlo method without long-range potential or hydrodynamic interactions to determine the diffusive motion of molecules ^102, 103^. The diameter of the crowding agents used in this simulation was 9.6 nm, comparable to those of a single nucleosome molecule determined by its volume ^2^. The diffusion coefficients (Ds) of the crowding agents were 0.3 µm^2^ s^−1^. The diameters of tracers were 10-20 nm. The Ds of tracers were given as 90 µm^2^ s^−1^ divided by its diameter in nanometers (9-4.5 µm^2^ s^−1^). These Ds were determined by the Stokes-Einstein relationship based on parameters from the EGFP monomer, the diameter and D of which were 3.8 nm and 23.5 µm^2^ s^−1^, respectively ^104^. Simulations were conducted in a 298-nm x 149-nm x 149-nm cuboid with reflective and periodic boundaries on long and other axes, respectively. These boundary conditions avoid problems caused by finite space. The simulation space had two different crowding conditions, which corresponded to 11.5 mg/ml (uncrowded region) and 85.9 mg/ml (chromatin domain) densities ^105^. 96 and 692 copies of 9.6-nm spheres (nucleosomes) were randomly placed in the left and right halves of the cuboid, respectively. The behavior of these crowding agents mimicked the diffusion of nucleosomes, which displaced less than a few nanometers (0-12 nm) from their initial positions (the “dog on a leash” model; see also ^104^ and ^106^). In addition, the centers of crowding agents were restricted in each half to keep the initial concentration. Initially, 10 tracers were randomly placed in the left (uncrowded) region. These tracers were allowed to diffuse freely in the whole cuboidal space. The motion of the tracers and crowding agents was iteratively simulated following previously described procedures ^104, 106^. The simulation time step, Δt, was 10 ns. 100-ms simulations were run, recording the position of the tracers every 10 µs. To analyze the search in genomic regions by molecules, the first collision times were logged for all pairs of tracers and crowding agents. 50 simulations (500 tracers in total) were performed for each condition. For the analysis of penetration kinetics, the time courses of the proportion of particles meeting the given conditions were fitted with an exponential function, 1 − *e*^(-*at* - *b*)^.

## Data availability

Data supporting the findings of this work are available in the Main text, Methods, Supplementary information, or Supplementary data. Source data are provided with this paper.

The genomics data have been deposited with links to BioProject accession number PRJDB19038 in the DDBJ BioProject database.

## Code availability

The numerical code in the chromatin simulation is publicly available on GitHub at https://github.com/kaizu/Minami2024. The scripts for track-sorting and angle-distribution analysis are available at https://zenodo.org/records/12672197.

## Acknowledgments

We are grateful to Dr. K. M. Marshall for critical reading and editing of this manuscript, Ms. K. Sato for establishing a H3.2-Halo expressing cell line, Dr. M. T. Kanemaki for providing HCT116 cells and Dr. Michael J. Hendzel for providing 10T1/2 cells. We thank Dr. H. Araki, Dr. H. Masai, Dr. Y. Saito, Dr. K. Hibino, Dr. M. T. Kanemaki, Dr. Y. Shimamoto, Dr. T. Torisawa, Dr. H. Niki, and Maeshima laboratory members for helpful discussions and support. This work was supported by the Japan Society for the Promotion of Science (JSPS) and MEXT KAKENHI grants (JP20H05936, JP21H02453, JP23K17398, JP22H04925 (PAGS), and JP24H00061) to K. Maeshima and JSPS and MEXT KAKENHI grants (21H02535 and JP22H05606) to S. I., and Takeda Science Foundation to K. Maeshima. K. Minami was a SOKENDAI Special Researcher (JST SPRING JPMJSP2104) and is a JSPS Fellow (JP23KJ0998).

## Author contributions

K. Minami and K. Maeshima designed the project. K. Minami generated H3.2-Halo expressing cells with the help of S. I. and performed single-nucleosome imaging, fluorescence imaging, and analyses. K. N. contributed to single-nucleosome imaging. K. Minami, S. T., S. I., and K. Maeshima performed nucleosome pull-down. K. Kaizu. and K. T. performed computational modeling. S. T. performed biochemical experiments and scientific illustrations. K. H., A. T., and K. Kurokawa contributed to the genomics analysis. K. Minami and K. Maeshima wrote the manuscript with input from all other authors.

## Competing interests

The authors declare no competing interests.

## Figures and Legends

**Fig. S1:**
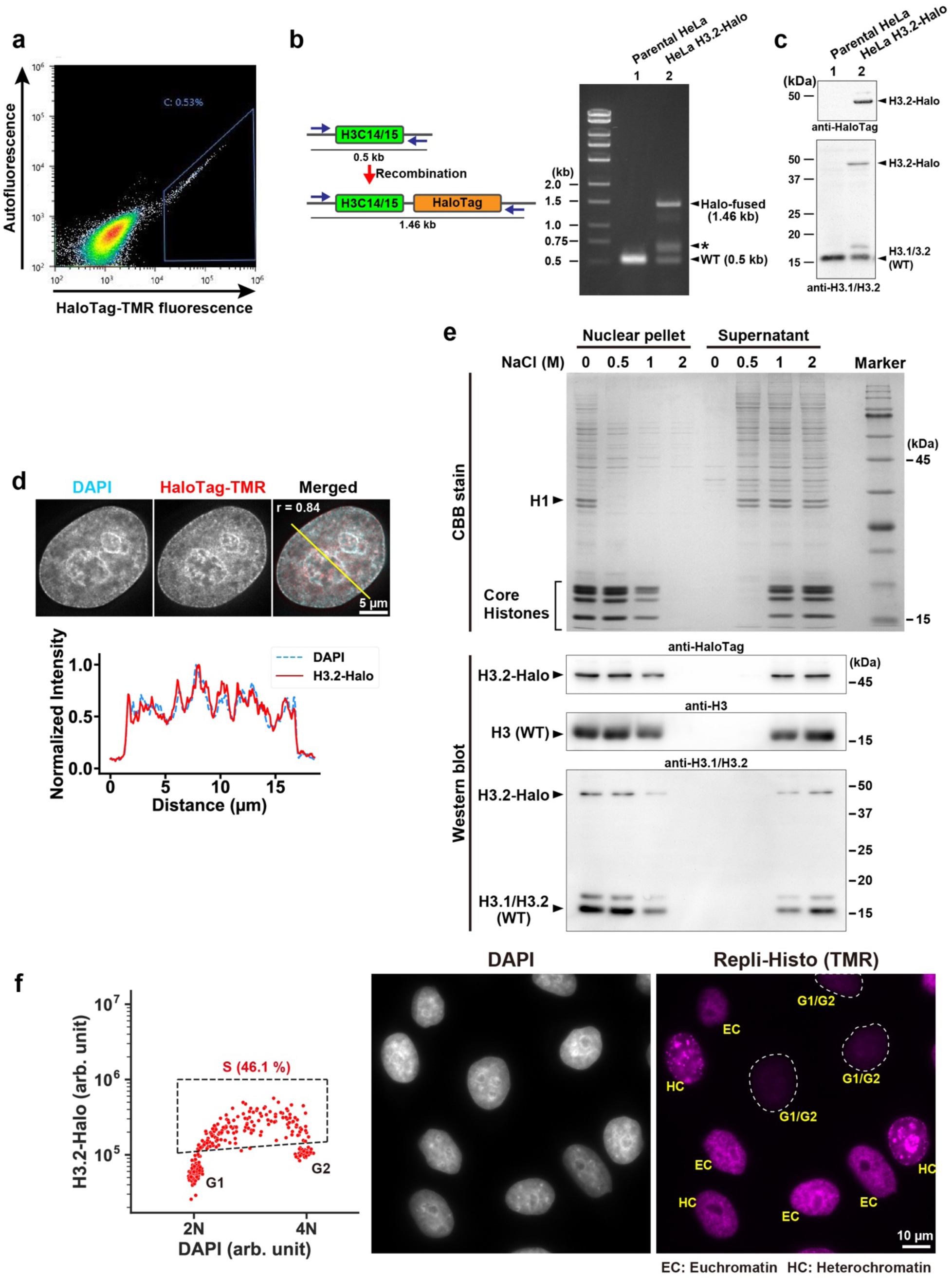
Establishment of HeLa cells expressing endogenous H3.2-HaloTag. **a**, The fluorescence-activated cell sorting (FACS) profile of the transfected HeLa cells. 0.53% of the cells show H3.2-Halo (TMR) fluorescence (blue gate). **b**, Polymerase chain reaction (PCR) to detect tagging of H3.2 with HaloTag. Expected PCR products and the primers used are shown on the left. A 1.46-kb product shows HaloTag insertion and a 0.5-kb product is from WT loci. The asterisk indicates a non-specific band derived from a DNA hybrid of the 1.46- and 0.5-kb fragments created during PCR. **c**, Western blots of H3.2-Halo in lysates of HeLa cells using an anti-histone H3.1/H3.2 antibody. Expressed H3.2-Halo is about 20% of H3.1/H3.2. The data with anti-HaloTag was reproduced from Fig. 1e. **d**, HeLa cells expressing H3.2-Halo labeled with HaloTag-tetramethylrhodamine (TMR) ligand (center). Left: DAPI staining, right: a merged image with Pearson’s correlation coefficient (DNA, cyan; H3.2-Halo, red). Bottom: the intensity line profiles of DAPI and H3.2-Halo taken across the yellow line shown in the top merged panel. Note that the localization of H3.2-Halo is like the DAPI-stained DNA. **e**, Stepwise salt washing of nuclei isolated from the HeLa H3.2-Halo. The nuclei isolated from HeLa cells expressing H3.2-Halo were washed with the indicated buffers including various concentrations of NaCl. The resultant nuclear pellets (left) and supernatants (right) were analyzed by SDS-PAGE, and subsequently stained with Coomassie brilliant blue (top) or immunoblotted with anti-HaloTag, anti-H3, and variant-specific anti-H3.1/H3.2 antibodies (bottom). Positions of core histones and linker histone H1 are indicated in the Coomassie brilliant blue stain. Note that H3 and H3.2-Halo started to dissociate from chromatin with 1 M NaCl and were detected in the supernatant fraction, suggesting that H3.2-Halo was incorporated into nucleosome structures like that of endogenous H3. **f**, Left: 46.1% of the asynchronous HeLa cells are in the S-phase with Repli-Histo signals (inside dotted line enclosure). The plot was replicated from Fig. 1g. Left: DAPI staining; right: the representative image of the Repli-Histo labeled cells. EC, euchromatin labeling; HC, heterochromatin labeling. Note that G1/G2 cells did not show the Repli-Histo signals.

**Fig. S2:**
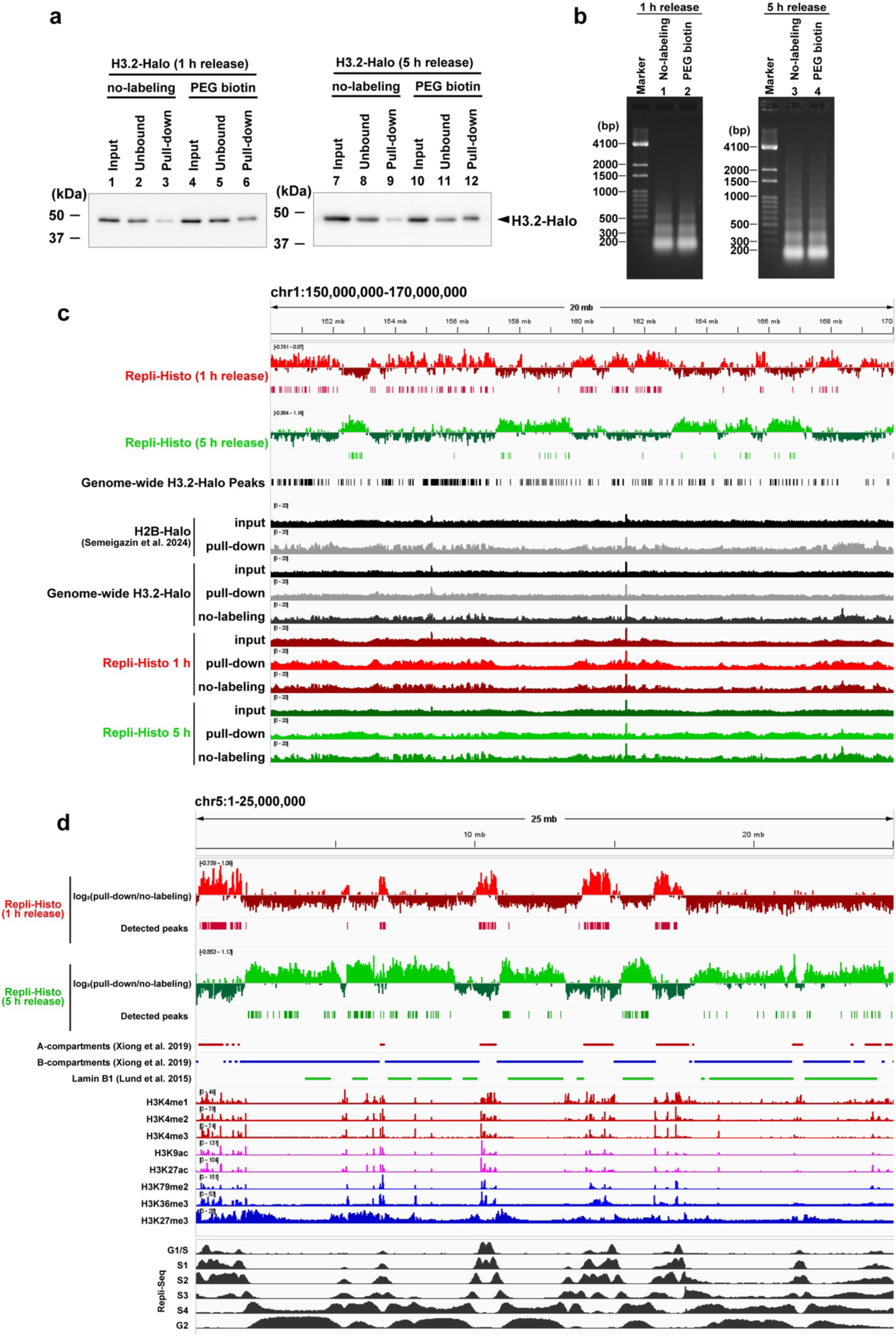
Pull-down and sequencing of the Repli-Histo labeled genomic regions. **a**, Enrichment of the pull-downed nucleosomes validated by western blotting with an anti-HaloTag antibody. Lanes 6 and 12 show the samples with a 1-h (euchromatin labeling) and 5-h (heterochromatin labeling) release from double thymidine block, respectively. **b**, Agarose gel electrophoresis of nucleosomal DNA purified from HeLa H3.2-Halo cells. Note that the major fractions are mononucleosomal DNA. **c**, The raw sequence profiles of the pull-downed DNA, view from chr1_150,000,000-170,000,000. Data for the H2B-Halo incorporated region were reproduced from ^50^. “No-labeling” indicates the non-specific precipitation. **d**, Another example of the Repli-Histo-labeled H3.2-Halo regions at 1 h and 5 h after release from thymidine blocks, view from chr5_1-25,000,000 using the Integrative Genomics Viewer (IGV) Browser.

**Fig. S3:**
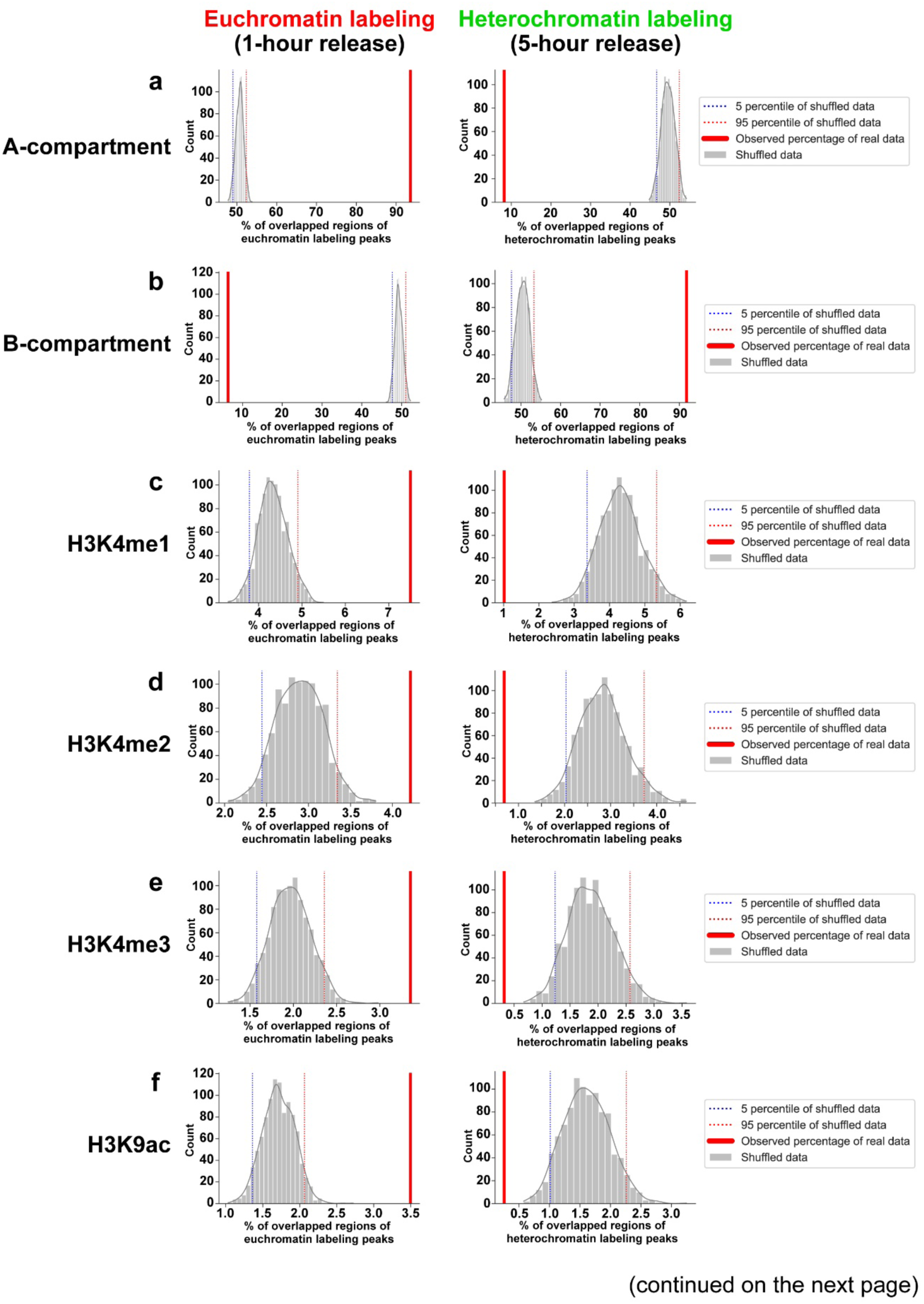
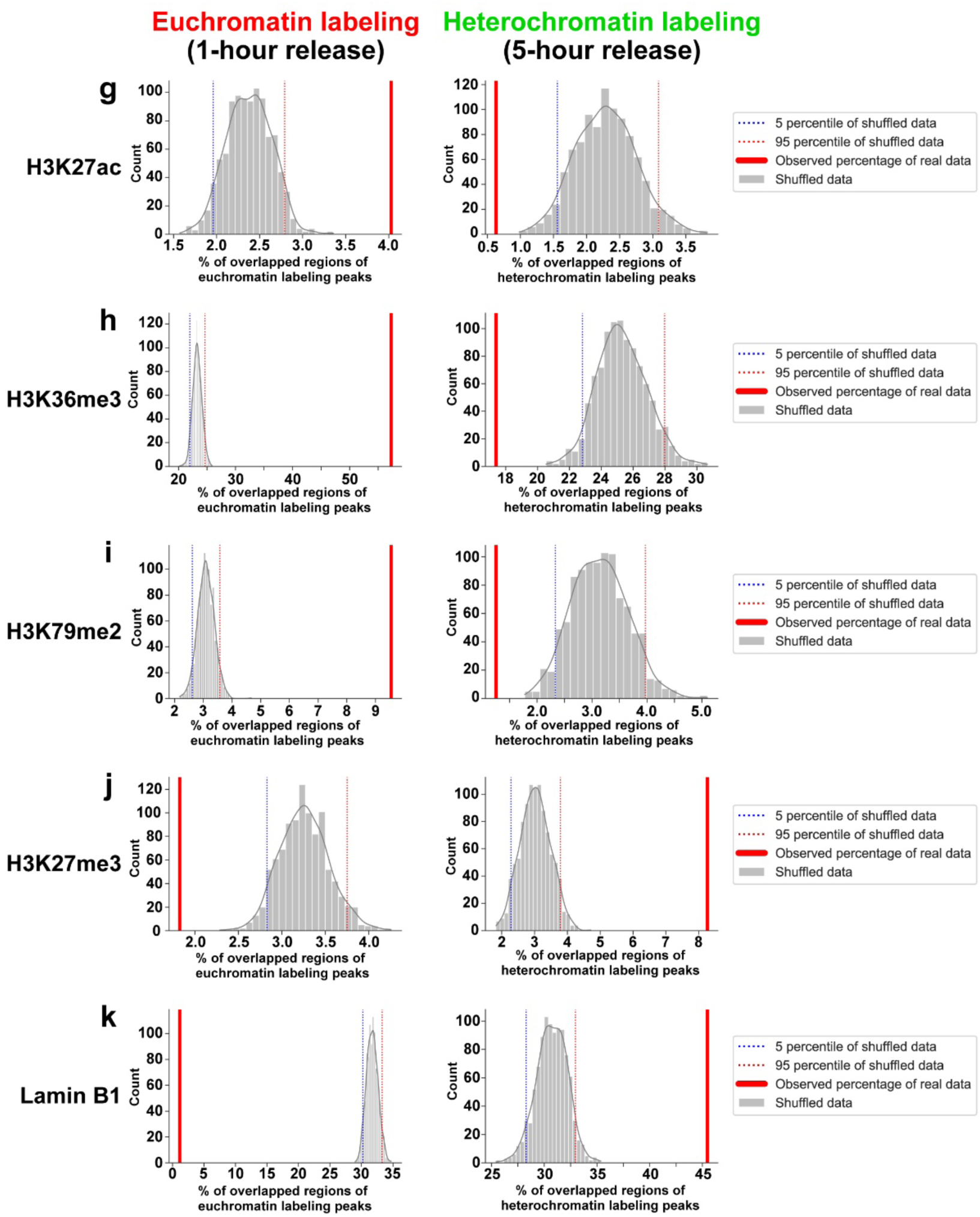
Statistical analysis between histone marks and Repli-Histo-labeled regions. **a**,**b**, Percentage distribution of regions of the Repli-Histo labeled peaks overlapped with (**a**) Hi-C A-compartment and (**b**) B-compartment in each of the 1,000 randomly shuffled datasets. The dotted lines show the 95th percentile of the shuffled distribution, and the red solid line shows the percentage observed in the real dataset of Repli-Histo-labeled (1-h or 5-h release) regions. **c**–**j**, Percentage distribution of regions of the Repli-Histo labeled peaks overlapped with active histone marks for H3K4me1 (**c**), H3K4me2 (**d**), H3K4me3 (**e**), H3K9ac (**f**), H3K27ac (**g**), H3K36me3 (**h**), H3K79me2 (**i**), and an inactive histone mark H3K27me3 (**j**). **k**, Percentage distribution of regions of the Repli-Histo labeled peaks overlapped with Lamin B1-associated regions ^79^.

**Fig. S4:**
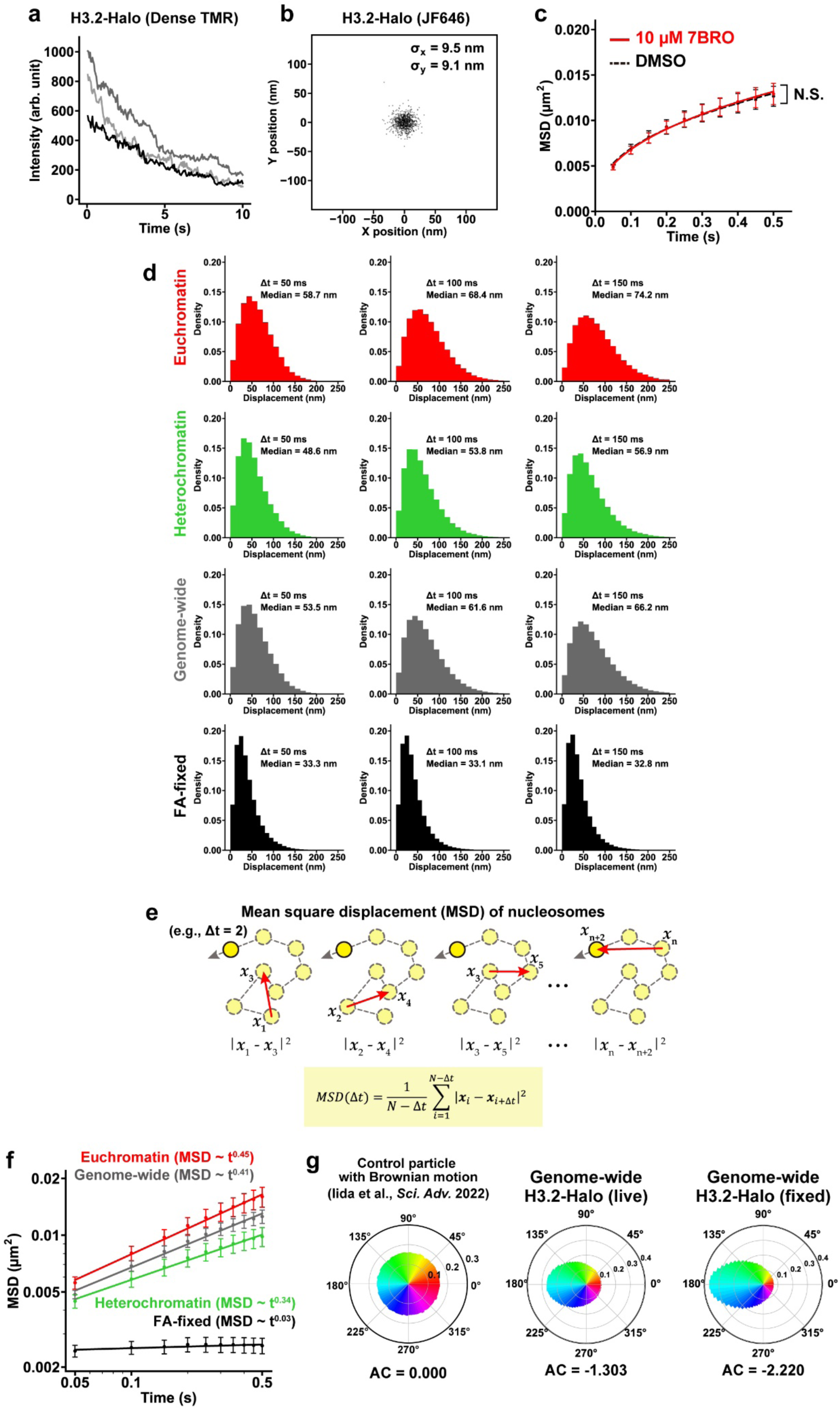
Euchromatin-/heterochromatin-specific single-nucleosome imaging with Repli-Histo labeling. **a**, Multi-step photobleaching of three typical TMR-labeled replication foci. Note that the gradual decrease in fluorescent intensity indicates that each focus was labeled with multiple TMR molecules. **b**, The position determination accuracy of JF646-labeled nucleosomes. Distribution of nucleosome displacements from the centroids in the XY-plane in the 50-ms interval, n = 11 nucleosomes in FA-fixed HeLa cells. SDx and SDy of fitted Gaussian functions were 9.5 nm and 9.1 nm, respectively. **c**, MSD plots (± SD among cells) of single nucleosomes in DMSO (black, n = 24 cells) or 7BRO treated (red, n = 23 cells) HeLa cells from 0.05 to 0.5 s. Genome-wide H3.2-Halo was labeled before the 7BRO treatment. N.S.: *P* = 0.4 by the two-sided Kolmogorov-Smirnov test. **d**, Distributions of the nucleosome displacement (movement) at 50 (left), 100 (center), and 150 (right) ms. Medians of displacement are indicated in each panel. **e**, Schematic of the mean square displacement (MSD) calculation. **f**, The log-log plot of MSDs from the plot of Fig. 3e. The plots were fitted linearly. The anomalous exponent calculated from the fitted lines is shown. **g**, Left: control angle distribution data from a particle with Brownian motion (also shown in Fig. 3h). The data was reproduced from ^54^. Center, right: motion angle distributions of nucleosomes in genome-wide-labeled living HeLa cells (n = 38 cells) and the FA-fixed control cells (n = 30 cells). Their AC values are shown at the bottom.

**Fig. S5:**
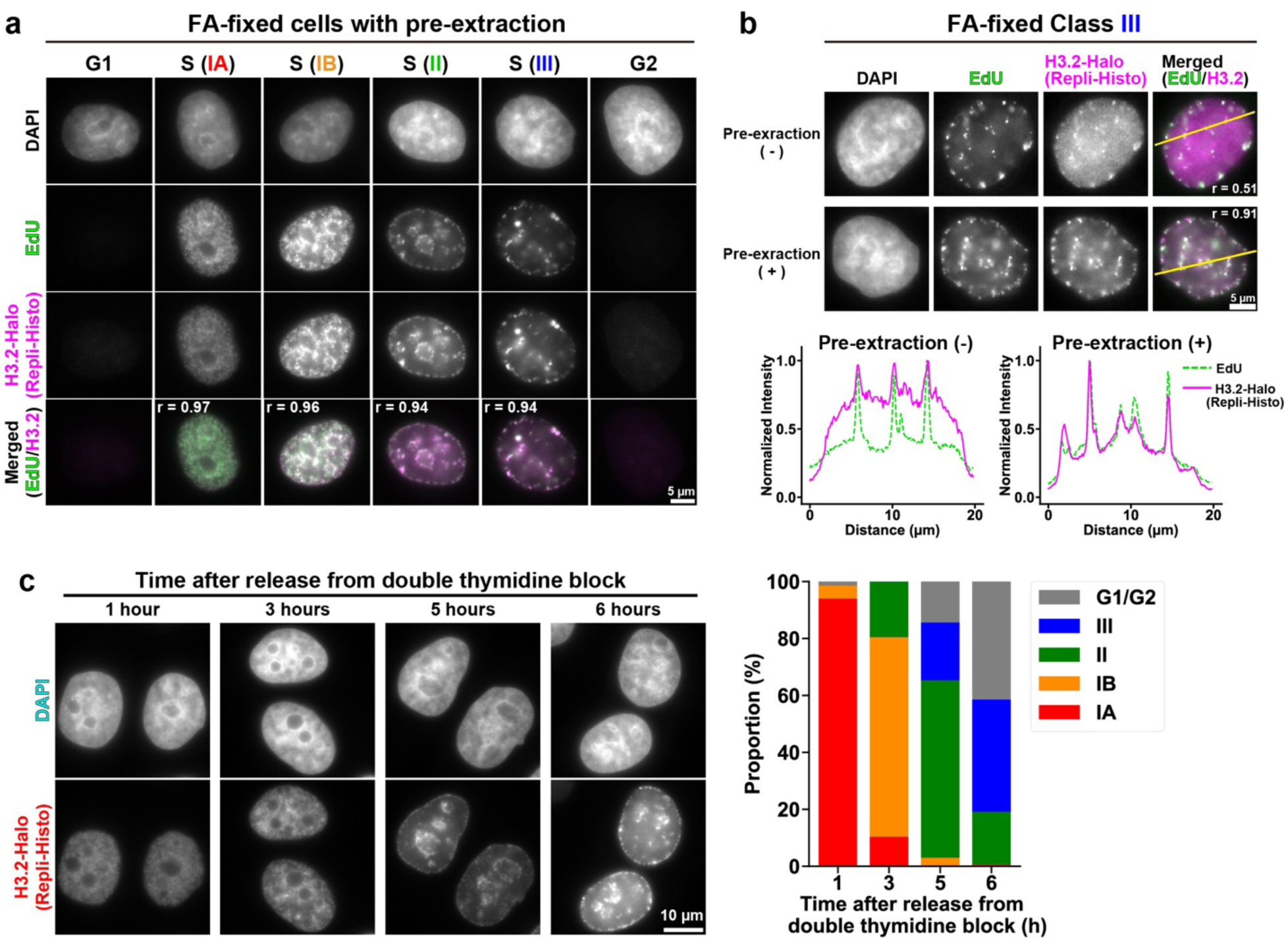
Four classes of chromatin identified with the Repli-Histo labeling. **a**, Asynchronous HeLa cells with DAPI staining (1^st^ row), EdU-pulse (2^nd^ row), and Repli-Histo labeling (3^rd^ row). Chromatin classes (IA, IB, II, and III) were categorized from the patterns of Repli-Histo labeling based on the previous studies ^61^ (see Table 1 for details). The bottom row shows the merged image of EdU (green) and Repli-Histo labeling (magenta) with a Pearson’s correlation coefficient. **b**, HeLa cells with the Class III Repli-Histo labeling (magenta) and EdU labeling (green). Pre-extraction (-), fixed with formaldehyde without extraction; Pre-extraction (+), treated with Triton X-100 before fixation. The intensity line profiles of EdU (green) and Repli-Histo labeling (magenta) across the yellow lines in the top panel are shown. Note that the diffusive Repli-Histo signal in Class III cells is diminished with the pre-extraction treatment. **c**, Left: Repli-Histo labeling at 1, 3, 5, and 6 h after release from the double-thymidine block (top, DAPI; bottom, Repli-Histo labeling of H3.2-Halo). Right: the proportion of each labeling pattern (IA/IB/II/III) among the synchronized HeLa cells in the left panel.

**Fig. S6:**
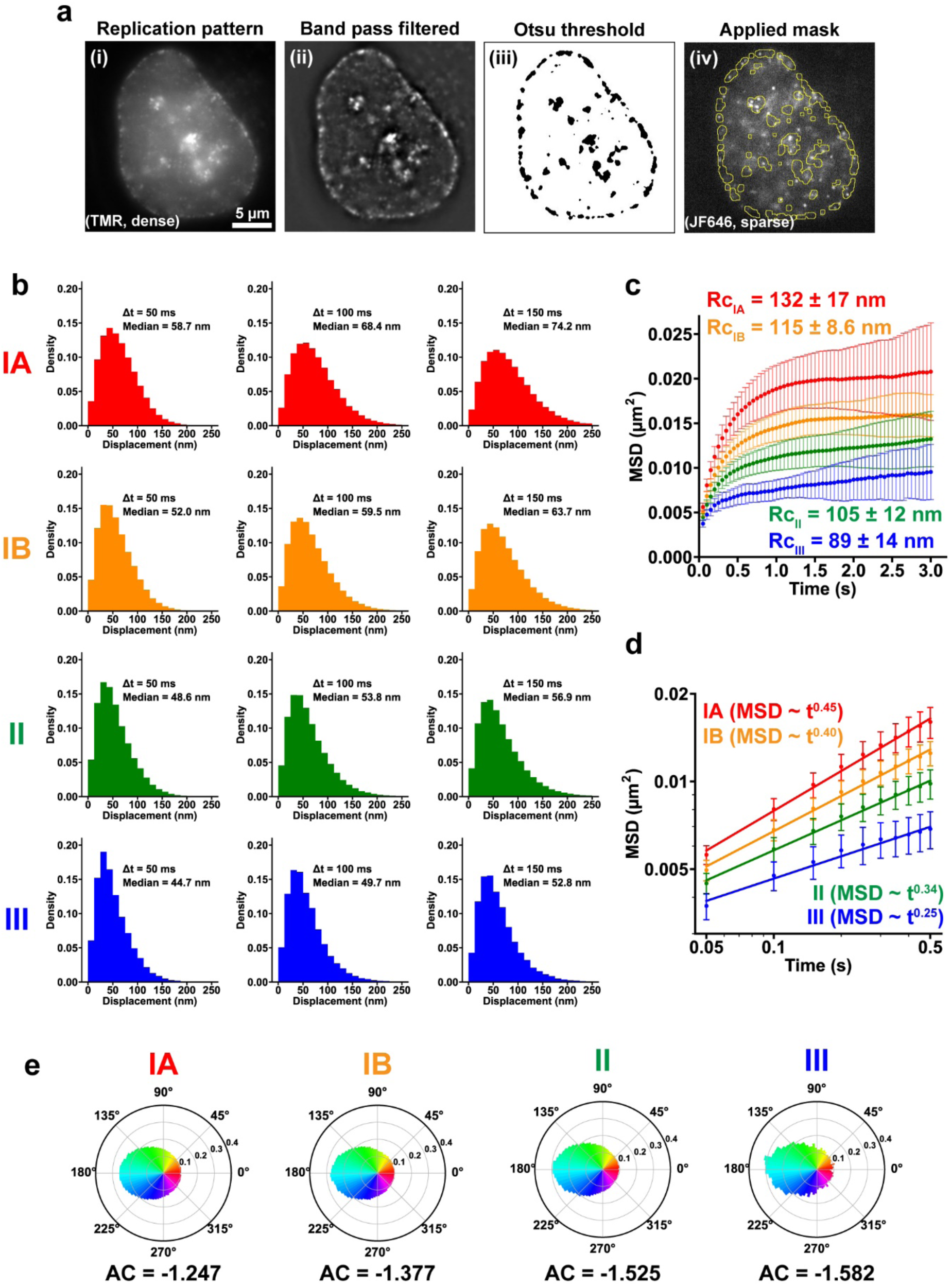
Local nucleosome movement in the four classes of chromatin regions. **a**, Extraction and segmentation of the Class III chromatin regions. The time-averaged dense TMR-labeled images (i) were high-pass filtered (ii) and binarized (iii) to define the Class III chromatin regions (iv). The JF646-single-nucleosome inside the regions (iv) were used in subsequent analyses. **b**, Distributions of the nucleosome displacement (movement) during 50, 100, and 150 ms. Medians of displacement are indicated in each panel. **c**, MSD plots (± SD among cells) of single nucleosomes in chromatin Class IA, IB, II, and III in a longer tracking time range from 0.05 to 3 s. The data set is from Fig. 4b. The estimated radii of constraint (Rc) of the nucleosome motion are shown. **d**, The log-log plot of MSDs from the plot of Fig. 4b. The plots were fitted linearly. The anomalous exponent values calculated from the fitted lines are shown. **e**, Motion angle distributions of nucleosomes in Class IA (red, n = 45 cells), IB (orange, n = 29), II (green, n = 36), and III (blue, n = 10) in living HeLa cells. Their AC values are shown at the bottom.

**Fig. S7:**
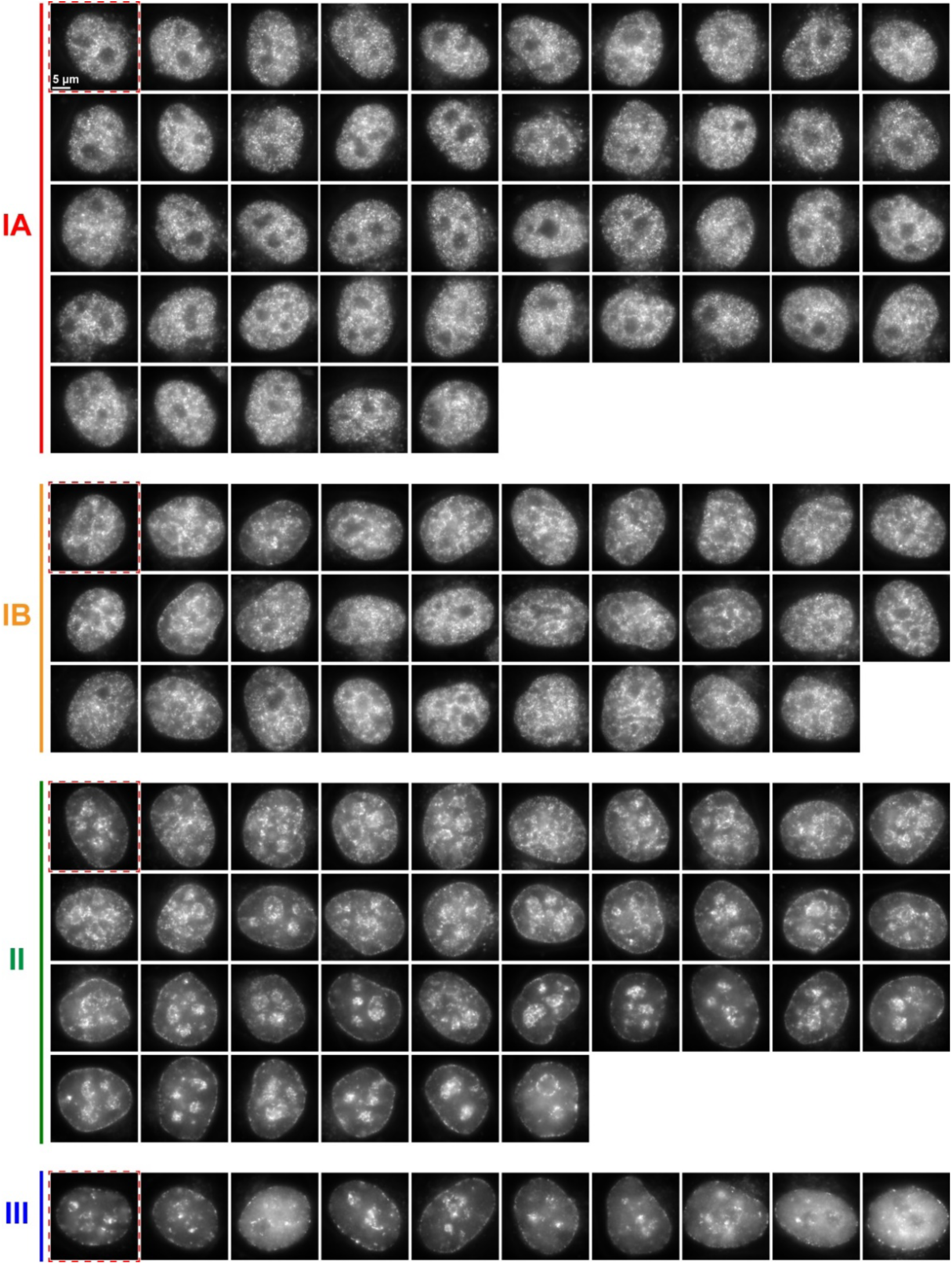
Repli-Histo labeling patterns in live HeLa cells. Repli-Histo labeling patterns of all observed cells for the single-nucleosome imaging in Fig 4b. The labeling patterns with dense TMR labeling are shown. The highlighted cells (red dashed squares) are also shown in Fig. 4a.

**Fig. S8:**
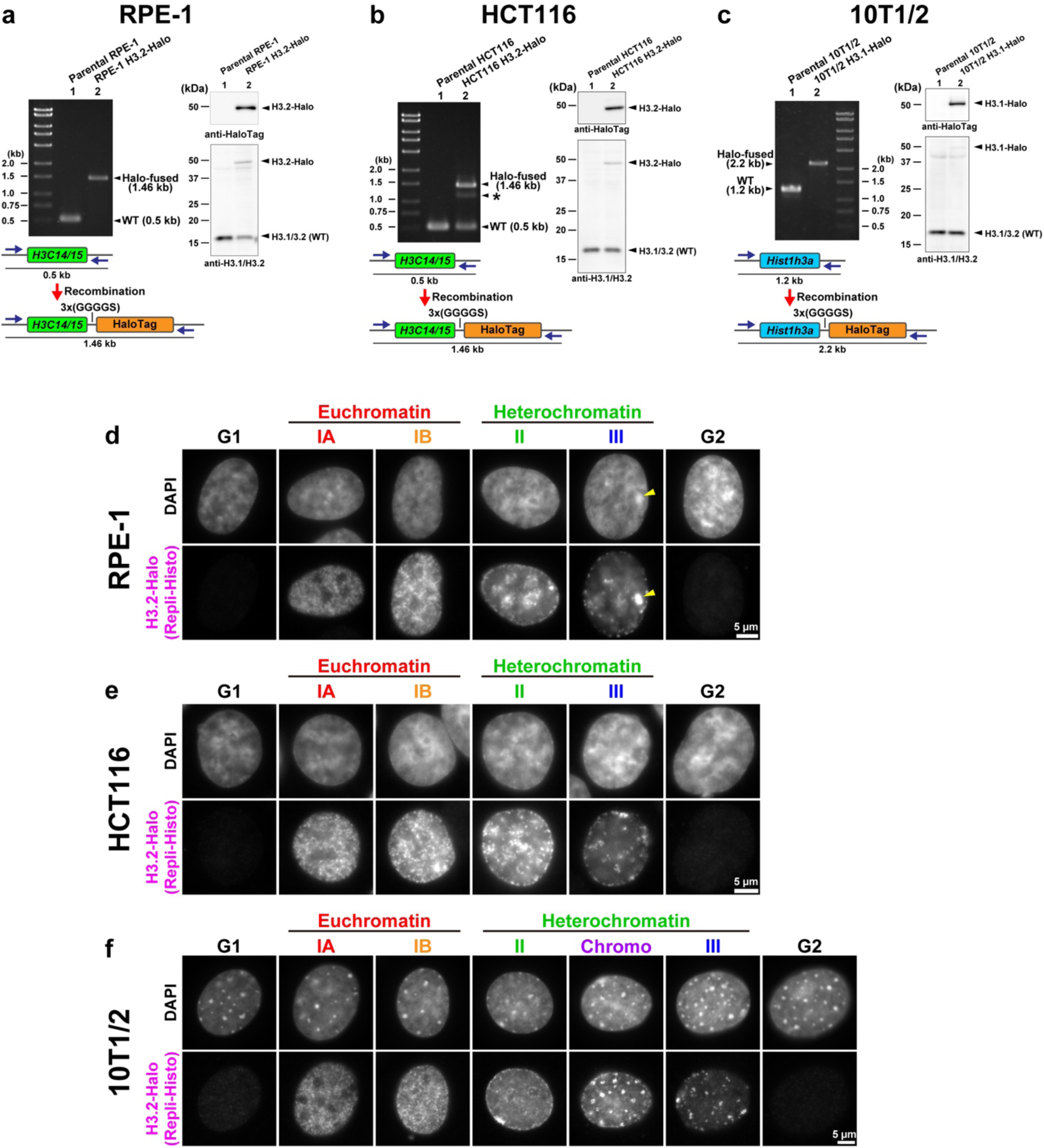
Repli-Histo single-nucleosome imaging in RPE-1, HCT116, and 10T1/2 cells. **a**–**c**, Left: PCR to validate tagging of H3.2/H3.1 with HaloTag in RPE-1, HCT116, and 10T1/2 cells. The asterisk in the HCT116 sample indicates a non-specific band derived from a DNA hybrid of the 1.46- and 0.5-kb fragments created during PCR. Right: western blots of H3.2/H3.1-Halo in lysates of RPE-1, HCT116, and 10T1/2 cells using an anti-HaloTag antibody and a control anti-histone H3.1/H3.2 antibody. Lane 1 is for their parental cells. **d**–**f**, RPE-1 (**d**), HCT116 (**e**), and mouse 10T1/2 (**f**) cells expressing H3.2/H3.1-Halo with Repli-Histo labeling. Chromatin classes (IA, IB, II, and III) were categorized from the patterns of Repli-Histo labeling based on the previous studies ^61^. The top row shows DAPI-stained DNA. Putative inactive X chromosome in RPE-1 cells, which is highly condensed, is indicated by arrow head (d).

**Fig. S9:**
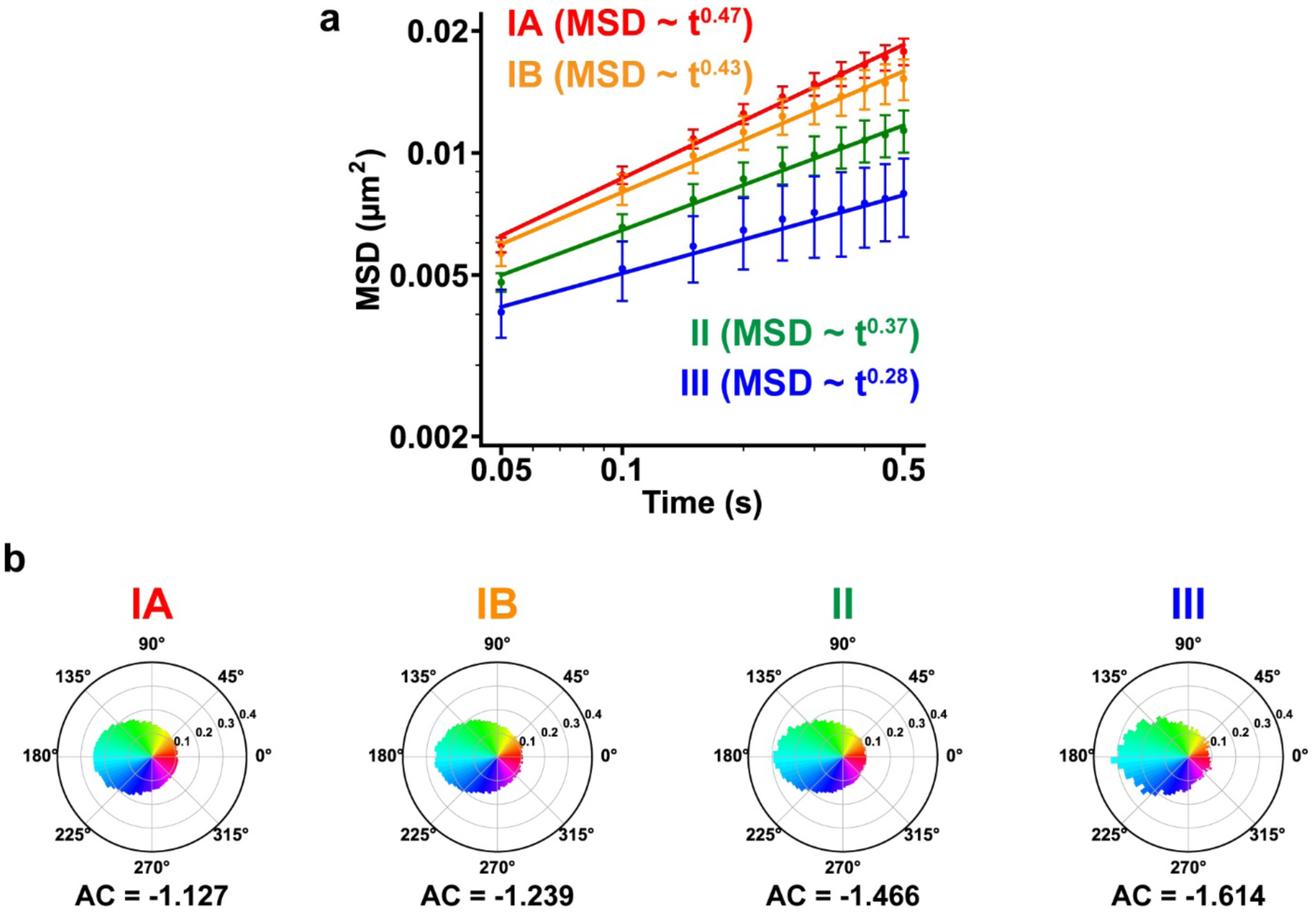
Single-nucleosome imaging with G1-arrested HeLa cells. **a**, The log-log plot of MSDs from the plot of Fig. 5c. The plots were fitted linearly. The anomalous exponent values calculated from the fitted lines are shown. **b**, Motion angle distributions of nucleosomes in Class IA (red, n = 37), IB (orange, n = 18), II (green, n = 15), and III (blue, n = 10) in living HeLa cells. Their AC values are shown at the bottom.

**Fig. S10:**
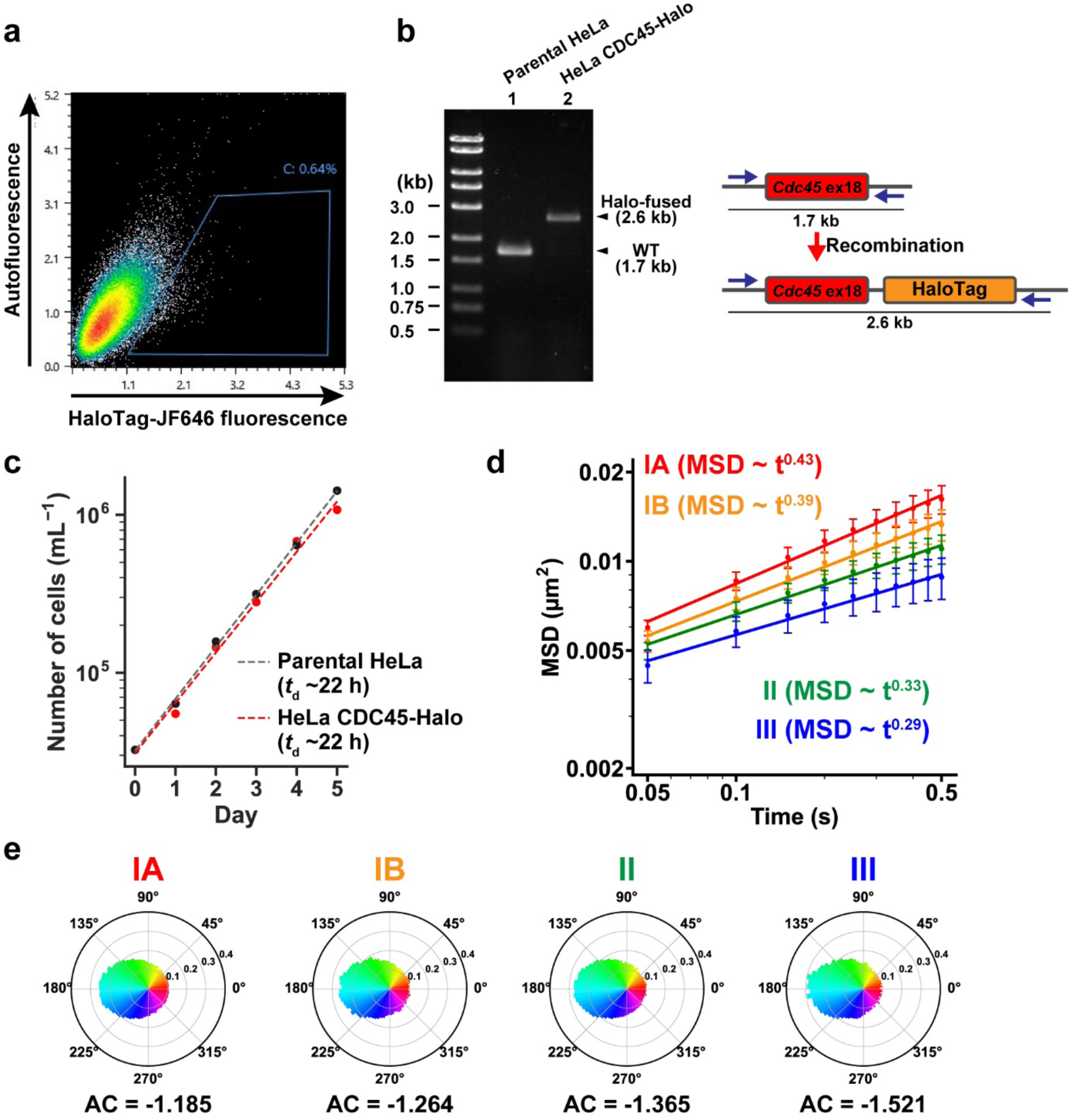
Single-CDC45 imaging reveals the nucleosome motion profile at the S phase. **a**, FACS profile of the transfected HeLa cells. 0.64% of the cells show CDC45-Halo-JF646 fluorescence (blue gate). **b**, PCR to validate tagging of CDC45 with HaloTag. Expected PCR products and the primers used are shown. **c**, Growth curves of parental HeLa (black) and HeLa expressing CDC45-Halo (red) cells cultured for indicated times. The estimated doubling times were 22 h for both cell lines. **d**, Log-log plot of MSDs from the plot of Fig. 6d. The plots were fitted linearly. The anomalous exponent values calculated from the fitted lines are shown. **e**, Motion angle distributions of nucleosomes in Class IA (red, n = 41 cells), IB (orange, n = 15), II (nuclear bottom; green, n = 23), and III (nuclear bottom; blue, n = 24) in living HeLa cells. Their AC values are shown at the bottom.

**Fig. S11:**
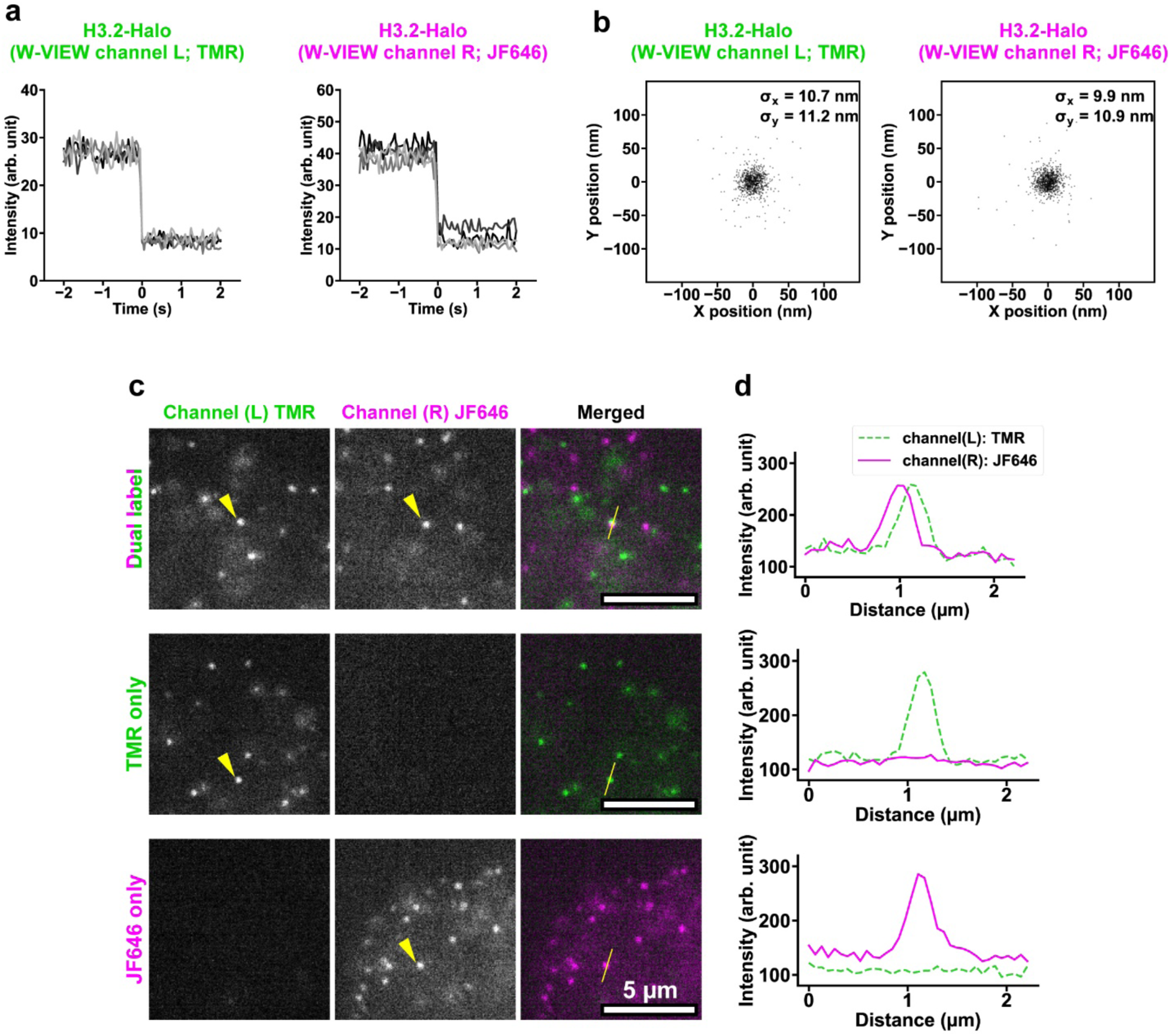
Dual-color single-nucleosome imaging with a beam splitter system. **a**, Single-step photobleaching of five representative TMR- (left) and JF646- (right) labeled nucleosome dots imaged with a beam splitter system. The horizontal axis shows the time before and after photobleaching. **b**, The position determination accuracy of TMR (left) and JF646- (right) labeled nucleosomes imaged with a beam splitter system. Distribution of nucleosome displacements from the centroids in the XY-plane in the 50-ms interval, n = 12 nucleosomes in FA-fixed HeLa cells. SDx and SDy of fitted Gaussian functions were 10.7 nm and 11.2 nm for TMR dots and 9.9 nm and 10.9 nm for JF646 dots. **c**–**d**, Representative single nucleosomes labeled with both TMR and JF646 (top), TMR only (middle) or JF646 only (bottom) in HeLa cells. The intensity line profiles for TMR and JF646 across the yellow lines in (**c**) are plotted in (**d**). Note that our beam splitter system has no crosstalk between the TMR and JF646 signals.

**Fig. S12:**
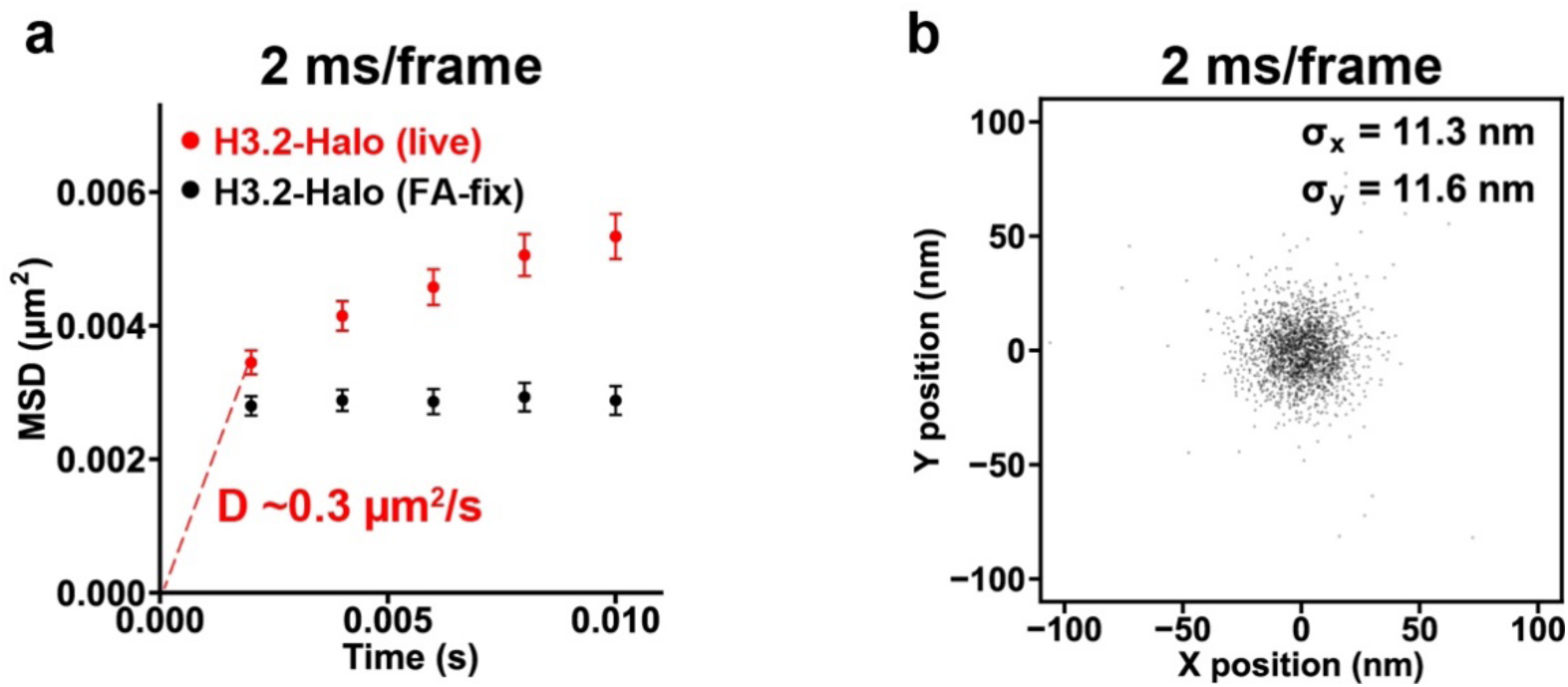
Single-nucleosome imaging at a fast frame rate and estimation of diffusion coefficient. **a**, MSD plots (± SD among cells) of single H3.2-Halo-nucleosomes in living HeLa cells from 2 to 10 ms (n = 16) at 2 ms/frame. Black shows the MSD from FA-fixed control cells (n = 15). The estimated diffusion coefficient from the first time point of the MSD was D ∼0.3 µm^2^/s. **b**, The position determination accuracy of JFX650^152^-labeled H3.2-Halo-nucleosomes at the 2 ms/frame imaging. Distribution of nucleosome displacements from the centroids in the XY-plane in the 2-ms interval, n = 12 nucleosomes in FA-fixed HeLa cells. SDx and SDy of fitted Gaussian functions were 11.3 nm and 11.6 nm, respectively.

**Fig. S13:**
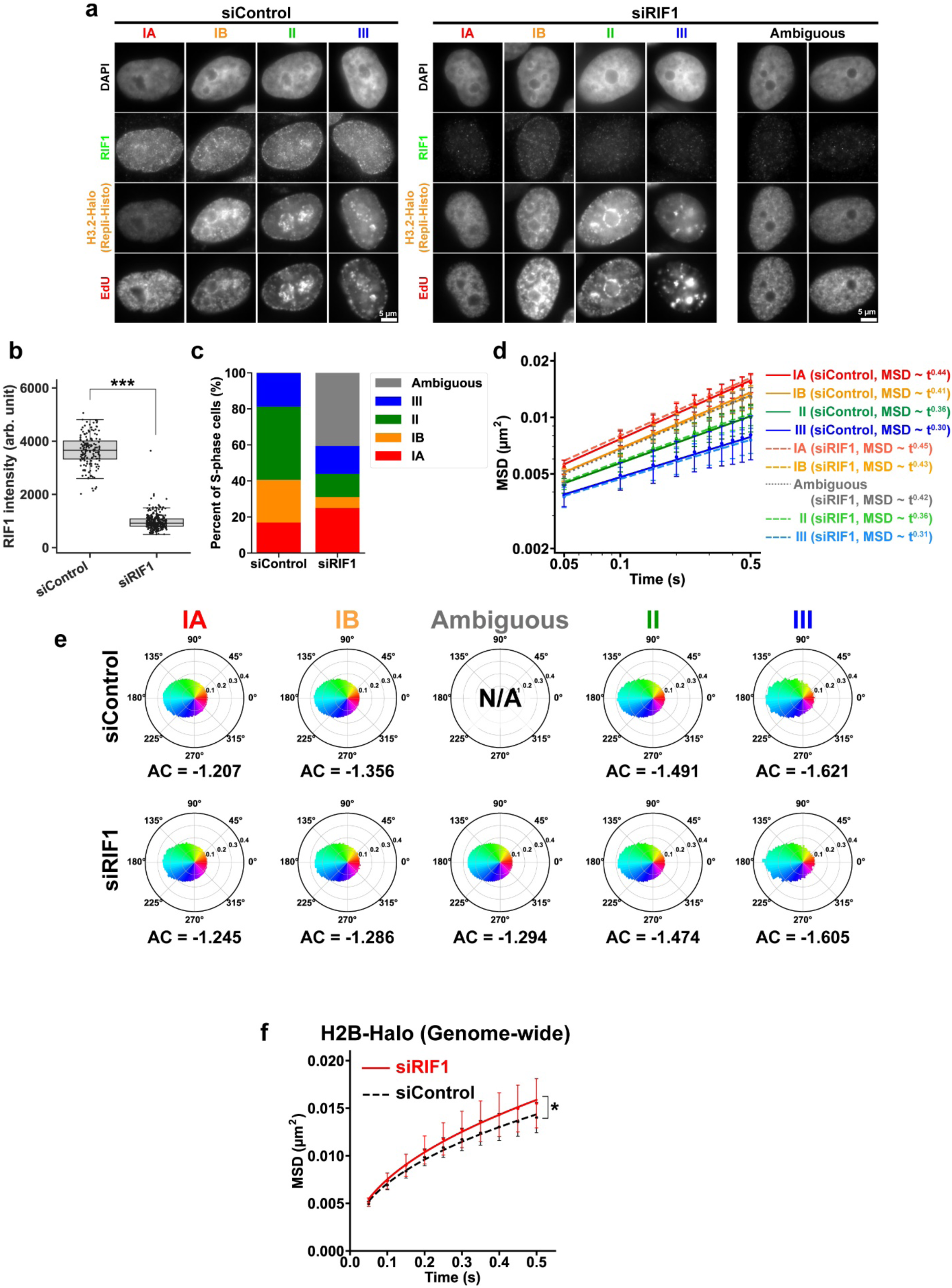
Chromatin motion profiles in RIF1-depleted cells. **a**, Validation of RIF1-KD by immunostaining (2^nd^ row) and the replication patterns visualized by Repli-Histo and EdU-pulse labeling (3^rd^ and 4^th^ rows). See Table 1 for the description of the labeling patterns. **b**, Quantifications of RIF1 intensity in (**a**); n = 159 cells for siControl and n = 297 cells for RIF1-KD. ***: *P* = 6.0 × 10^-69^ by two-sided Mann-Whitney U-test. **c**, The percentage of each Repli-Histo labeling pattern (IA/IB/II/III or ambiguous pattern). Note that ambiguous replication patterns with dispersed (or uniform) replicated regions appear in RIF1-KD cells. **d**, The log-log plot of MSDs from the plot of Fig. 8c. The plots were fitted linearly. The anomalous exponents calculated from the fitted lines are shown. **e**, Motion angle distributions of nucleosomes in Class IA (control, n = 43; RIF1-KD, n = 27), IB (control, n = 35; RIF1-KD, n = 22), II (control, n = 40; RIF1-KD, n = 24), III (control, n = 19; RIF1-KD, n = 23), and ambiguous patterns (RIF1-KD, n = 72) in living HeLa cells. Their AC values are shown at the bottom. **f**, MSD plots (± SD among cells) of genome-wide-labeled single nucleosomes (H2B-Halo) in HeLa cells with indicated conditions: siControl, black; RIF1-depleted (siRIF1), red; n =21 cells for siControl and 17 cells for siRIF1. *: *P* = 0.046 by Kolmogorov-Smirnov test (two-sided).

## Movies S1–S31 legends

**Movie S1**

Left: the euchromatin-specific Repli-Histo labeling in a living HeLa cell. Labeling with dense TMR shows the localization of euchromatin. Right: movie (50 ms/frame) of the corresponding single nucleosomes labeled with JF646 recorded by the sCMOS ORCA-Fusion BT camera (Hamamatsu Photonics). Note that clear and well-separated dots are visualized with single-step photobleaching profiles (Fig. 3c), suggesting that each dot represents a single H3.2-Halo-JF646 molecule in a single nucleosome. Scale bar: 5 µm.

**Movie S2**

Left: the heterochromatin-specific Repli-Histo labeling in a living HeLa cell. Labeling with dense TMR shows the localization of heterochromatin. Right: movie (50 ms/frame) of the corresponding single nucleosomes labeled with JF646. Scale bar: 5 µm.

**Movie S3**

Movie (50 ms/frame) of single nucleosomes labeled with JF646 in a formaldehyde (FA)-fixed HeLa interphase cell. Scale bar: 5 µm.

**Movie S4**

Movie (50 ms/frame) of H3.2-Halo single nucleosomes, labeled genome-wide with JF646 in a living HeLa interphase cell. Scale bar: 5 µm.

**Movie S5**

Left: a living HeLa cell with the Class IA Repli-Histo labeling with dense TMR. Right: movie (50 ms/frame) of the corresponding single nucleosomes labeled with JF646. Scale bar: 5 µm.

**Movie S6**

Left: a living HeLa cell with the Class IB Repli-Histo labeling. Right: movie (50 ms/frame) of the corresponding single nucleosomes labeled with JF646. Scale bar: 5 µm.

**Movie S7**

Left: a living HeLa cell with the Class II Repli-Histo labeling. Right: movie (50 ms/frame) of the corresponding single nucleosomes labeled with JF646. Scale bar: 5 µm.

**Movie S8**

Left: a living HeLa cell with the Class III Repli-Histo labeling. Right: movie (50 ms/frame) of the corresponding single nucleosomes labeled with JF646. Rapidly diffusing dots are free H3.2-Halo (Fig. S5b), and we only focused on the behavior of H3.2-Halo stably incorporated into nucleosomes (Fig. S6a). Scale bar: 5 µm.

**Movie S9**

Left: a living RPE-1 cell with the Class IA Repli-Histo labeling. Right: movie (50 ms/frame) of the corresponding single nucleosomes labeled with JF646. Scale bar: 5 µm.

**Movie S10**

Left: a living RPE-1 cell with the Class IB Repli-Histo labeling. Right: movie (50 ms/frame) of the corresponding single nucleosomes labeled with JF646. Scale bar: 5 µm.

**Movie S11**

Left: a living RPE-1 cell with the Class II Repli-Histo labeling. Right: movie (50 ms/frame) of the corresponding single nucleosomes labeled with JF646. Scale bar: 5 µm.

**Movie S12**

Left: a living RPE-1 cell with the Class III Repli-Histo labeling. Right: movie (50 ms/frame) of the corresponding single nucleosomes labeled with JF646. Scale bar: 5 µm.

**Movie S13**

Left: a G1-arrested living HeLa cell with the Class IA Repli-Histo labeling. Right: movie (50 ms/frame) of the corresponding single nucleosomes labeled with JF646. Scale bar: 5 µm.

**Movie S14**

Left: a G1-arrested living HeLa cell with the Class IB Repli-Histo labeling. Right: movie (50 ms/frame) of the corresponding single nucleosomes labeled with JF646. Scale bar: 5 µm.

**Movie S15**

Left: a G1-arrested living HeLa cell with the Class II Repli-Histo labeling. Right: movie (50 ms/frame) of the corresponding single nucleosomes labeled with JF646. Scale bar: 5 µm.

**Movie S16**

Left: a G1-arrested living HeLa cell with the Class III Repli-Histo labeling. Right: movie (50 ms/frame) of the corresponding single nucleosomes labeled with JF646. Rapidly diffusing dots are free H3.2-Halo, but we only focused on the behavior of H3.2-Halo stably incorporated into nucleosomes. Scale bar: 5 µm.

**Movie S17**

Left: a living G1-phase HeLa cell with CDC45-Halo. Labeling with dense TMR shows the localization of CDC45-Halo. Right: movie (50 ms/frame) of the corresponding single CDC45-Halo molecules labeled with JF646. Most of the CDC45-Halo molecules freely diffuse in the nucleoplasm. Scale bar: 5 µm.

**Movie S18**

Left: a living S-phase HeLa cell with the Class IA CDC45-Halo foci. Right: movie (50 ms/frame) of the corresponding single CDC45-Halo molecules labeled with JF646. CDC45-Halo molecules start stably binding to nucleosomes, likely engaging DNA replication. Scale bar: 5 µm.

**Movie S19**

Left: a living S-phase HeLa cell with the Class IB CDC45-Halo foci. Right: movie (50 ms/frame) of the corresponding single CDC45-Halo molecules labeled with JF646. Scale bar: 5 µm.

**Movie S20**

Left: a living S-phase HeLa cell with the Class II CDC45-Halo foci. Right: movie (50 ms/frame) of the corresponding single CDC45-Halo molecules labeled with JF646. Scale bar: 5 µm.

**Movie S21**

Left: a living S-phase HeLa cell with the Class III CDC45-Halo foci. Right: movie (50 ms/frame) of the corresponding single CDC45-Halo molecules labeled with JF646. Scale bar: 5 µm.

**Movie S22**

Movie (50 ms/frame) of the TMR- and JF646-labeled single nucleosomes with the euchromatin-specific Repli-Histo labeling in a living HeLa cell. The two channels were recorded simultaneously using W-VIEW GEMINI (Hamamatsu Photonics). Left: TMR-labeled nucleosomes (green). Center: JF646-labeled nucleosomes (magenta). Right: the merged image. Scale bar: 5 µm.

**Movie S23**

A magnified view of Movie S22, showing the closely labeled single-nucleosome pairs (yellow arrows). Left: TMR, center: JF646, right: merged. Scale bar, 1 µm.

**Movie S24**

Movie (50 ms/frame) of the TMR- and JF646-labeled single nucleosomes with the heterochromatin-specific Repli-Histo labeling in a living HeLa cell. Left: TMR-labeled nucleosomes (green). Center: JF646-labeled nucleosomes (magenta). (Right) the merged image. Scale bar: 5 µm.

**Movie S25**

Movie (50 ms/frame) of the TMR- and JF646-labeled single nucleosomes with the euchromatin-specific Repli-Histo labeling in an FA-fixed HeLa cell. Left: TMR-labeled nucleosomes (green). Center: JF646-labeled nucleosomes (magenta). Right: the merged image. Scale bar: 5 µm.

**Movie S26**

Movie (50 ms/frame) of the TMR- and JF646-labeled single nucleosomes with the heterochromatin-specific Repli-Histo labeling in an FA-fixed HeLa cell. Left: TMR-labeled nucleosomes (green). Center: JF646-labeled nucleosomes (magenta). Right: the merged image. Scale bar: 5 µm.

**Movie S27**

The 10-nm model diffusion proteins (tracer, blue) in the immobile chromatin environment (red, fluctuation radius (Rf) of 0 nm) during 1 ms of the simulation.

**Movie S28**

The 10-nm model diffusion proteins (tracer, blue) in the fluctuating chromatin environment (red, fluctuation radius (Rf) of 12 nm) during 1 ms of the simulation.

**Movie S29**

The 15-nm model diffusion proteins (tracer, blue) in the immobile chromatin environment (red, fluctuation radius (Rf) of 0 nm) during 1 ms of the simulation. The tracers cannot reach the dense chromatin domain with immobile nucleosomes (right).

**Movie S30**

The 15-nm model diffusion proteins (tracer, blue) in the fluctuating chromatin environment (red, fluctuation radius (Rf) of 12 nm) during 1 ms of the simulation. Local nucleosome fluctuation increases chromatin accessibility, and large tracers can penetrate the dense chromatin domain (right).

**Movie S31**

Left: a RIF1-depleted living HeLa cell with ambiguous pattern Repli-Histo labeling. Right: movie (50 ms/frame) of the corresponding single nucleosomes labeled with JF646. Scale bar: 5 µm.

## Notes

### Competing Interest Statement

The authors have declared no competing interest.

## References

1. Olins, D.E. & Olins, A.L. Chromatin history: our view from the bridge. Nat Rev Mol Cell Biol 4, 809–814 (2003).

2. Luger, K., Mader, A.W., Richmond, R.K., Sargent, D.F. & Richmond, T.J. Crystal structure of the nucleosome core particle at 2.8 A resolution. Nature 389, 251–260 (1997).

3. Koyama, M. & Kurumizaka, H. Structural diversity of the nucleosome. J Biochem 163, 85–95 (2018).

4. McGinty, R.K. & Tan, S. Nucleosome structure and function. Chem Rev 115, 2255–2273 (2015).

5. Misteli, T. The Self-Organizing Genome: Principles of Genome Architecture and Function. Cell 183, 28–45 (2020).

6. Maeshima, K., Iida, S. & Tamura, S. Physical Nature of Chromatin in the Nucleus. Cold Spring Harb Perspect Biol 13 (2021).

7. Belmont, A.S. & Bruce, K. Visualization of G1 chromosomes: a folded, twisted, supercoiled chromonema model of interphase chromatid structure. J Cell Biol 127, 287–302 (1994).

8. Nozaki, T. et al. Dynamic Organization of Chromatin Domains Revealed by Super-Resolution Live-Cell Imaging. Mol Cell 67, 282–293 e287 (2017).

9. Bintu, B. et al. Super-resolution chromatin tracing reveals domains and cooperative interactions in single cells. Science 362 (2018).

10. Miron, E. et al. Chromatin arranges in chains of mesoscale domains with nanoscale functional topography independent of cohesin. Sci Adv 6 (2020).

11. Cremer, T. et al. The Interchromatin Compartment Participates in the Structural and Functional Organization of the Cell Nucleus. Bioessays 42, e1900132 (2020).

12. Nozaki, T. et al. Condensed but liquid-like domain organization of active chromatin regions in living human cells. Sci Adv 9, eadf1488 (2023).

13. Zakirov, A.N. et al. Fiber-Like Organization as a Basic Principle for Euchromatin Higher-Order Structure. Front Cell Dev Biol 9, 784440 (2021).

14. Li, Y. et al. Analysis of three-dimensional chromatin packing domains by chromatin scanning transmission electron microscopy (ChromSTEM). Sci Rep 12, 12198 (2022).

15. Maeshima, K., Iida, S., Shimazoe, M.A., Tamura, S. & Ide, S. Is euchromatin really open in the cell? Trends Cell Biol 34, 7–17 (2024).

16. Lieberman-Aiden, E. et al. Comprehensive mapping of long-range interactions reveals folding principles of the human genome. Science 326, 289–293 (2009).

17. Nora, E.P. et al. Spatial partitioning of the regulatory landscape of the X-inactivation centre. Nature 485, 381–385 (2012).

18. Dixon, J.R. et al. Topological domains in mammalian genomes identified by analysis of chromatin interactions. Nature 485, 376–380 (2012).

19. Sexton, T. et al. Three-dimensional folding and functional organization principles of the Drosophila genome. Cell 148, 458–472 (2012).

20. Rao, S.S. et al. A 3D map of the human genome at kilobase resolution reveals principles of chromatin looping. Cell 159, 1665–1680 (2014).

21. Grewal, S.I. & Jia, S. Heterochromatin revisited. Nat Rev Genet 8, 35–46 (2007).

22. Allshire, R.C. & Madhani, H.D. Ten principles of heterochromatin formation and function. Nat Rev Mol Cell Biol 19, 229–244 (2018).

23. Janssen, A., Colmenares, S.U. & Karpen, G.H. Heterochromatin: Guardian of the Genome. Annual review of cell and developmental biology 34, 265–288 (2018).

24. Padeken, J., Methot, S.P. & Gasser, S.M. Establishment of H3K9-methylated heterochromatin and its functions in tissue differentiation and maintenance. Nat Rev Mol Cell Biol 23, 623–640 (2022).

25. Bell, O., Burton, A., Dean, C., Gasser, S.M. & Torres-Padilla, M.E. Heterochromatin definition and function. Nat Rev Mol Cell Biol 24, 691–694 (2023).

26. Alberts, B., et al. Molecular Biology of the Cell, Seventh Edition (2022).

27. Pollard, T.D., Earnshaw, W.C., Lippincott-Schwartz, J. & Johnson, G. Cell Biology, 4th Edition. (2022).

28. Gelleri, M. et al. True-to-scale DNA-density maps correlate with major accessibility differences between active and inactive chromatin. Cell Rep 42, 112567 (2023).

29. Chubb, J.R., Boyle, S., Perry, P. & Bickmore, W.A. Chromatin motion is constrained by association with nuclear compartments in human cells. Curr Biol 12, 439–445 (2002).

30. Shinkai, S., Nozaki, T., Maeshima, K. & Togashi, Y. Dynamic Nucleosome Movement Provides Structural Information of Topological Chromatin Domains in Living Human Cells. PLoS Comput Biol 12, e1005136 (2016).

31. Shaban, H.A., Barth, R. & Bystricky, K. Formation of correlated chromatin domains at nanoscale dynamic resolution during transcription. Nucleic Acids Res 46, e77 (2018).

32. Uchino, S. et al. Live imaging of transcription sites using an elongating RNA polymerase II-specific probe. J Cell Biol 221 (2022).

33. Lerner, J. et al. Two-Parameter Mobility Assessments Discriminate Diverse Regulatory Factor Behaviors in Chromatin. Mol Cell 79, 677–688 e676 (2020).

34. Schermelleh, L., Solovei, I., Zink, D. & Cremer, T. Two-color fluorescence labeling of early and mid-to-late replicating chromatin in living cells. Chromosome Res 9, 77–80 (2001).

35. Kimura, H. & Cook, P.R. Kinetics of core histones in living human cells: little exchange of H3 and H4 and some rapid exchange of H2B. J Cell Biol 153, 1341–1353 (2001).

36. Meshorer, E. et al. Hyperdynamic plasticity of chromatin proteins in pluripotent embryonic stem cells. Dev Cell 10, 105–116 (2006).

37. Hajjoul, H. et al. High-throughput chromatin motion tracking in living yeast reveals the flexibility of the fiber throughout the genome. Genome Res 23, 1829–1838 (2013).

38. Heun, P., Laroche, T., Shimada, K., Furrer, P. & Gasser, S.M. Chromosome dynamics in the yeast interphase nucleus. Science 294, 2181–2186 (2001).

39. Levi, V., Ruan, Q., Plutz, M., Belmont, A.S. & Gratton, E. Chromatin dynamics in interphase cells revealed by tracking in a two-photon excitation microscope. Biophys J 89, 4275–4285 (2005).

40. Albiez, H. et al. Chromatin domains and the interchromatin compartment form structurally defined and functionally interacting nuclear networks. Chromosome Res 14, 707–733 (2006).

41. Germier, T. et al. Real-Time Imaging of a Single Gene Reveals Transcription-Initiated Local Confinement. Biophys J 113, 1383–1394 (2017).

42. Chen, B. et al. Dynamic imaging of genomic loci in living human cells by an optimized CRISPR/Cas system. Cell 155, 1479–1491 (2013).

43. Gu, B. et al. Transcription-coupled changes in nuclear mobility of mammalian cis-regulatory elements. Science 359, 1050–1055 (2018).

44. Ma, H. et al. Cell cycle- and genomic distance-dependent dynamics of a discrete chromosomal region. J Cell Biol 218, 1467–1477 (2019).

45. Zidovska, A., Weitz, D.A. & Mitchison, T.J. Micron-scale coherence in interphase chromatin dynamics. Proc Natl Acad Sci U S A 110, 15555–15560 (2013).

46. Saxton, M.N., Morisaki, T., Krapf, D., Kimura, H. & Stasevich, T.J. Live-cell imaging uncovers the relationship between histone acetylation, transcription initiation, and nucleosome mobility. Sci Adv 9, eadh4819 (2023).

47. Ide, S., Tamura, S. & Maeshima, K. Chromatin behavior in living cells: lessons from single-nucleosome imaging and tracking. BioEssays 44, e2200043 (2022).

48. Lakadamyali, M. Single nucleosome tracking to study chromatin plasticity. Curr Opin Cell Biol 74, 23–28 (2022).

49. Ashwin, S.S., Nozaki, T., Maeshima, K. & Sasai, M. Organization of fast and slow chromatin revealed by single-nucleosome dynamics. Proc Natl Acad Sci U S A 116, 19939–19944 (2019).

50. Semeigazin, A. et al. Behaviors of nucleosomes with mutant histone H4s in euchromatic domains of living human cells. Histochem Cell Biol (2024).

51. Wagh, K. et al. Dynamic switching of transcriptional regulators between two distinct low-mobility chromatin states. Sci Adv 9, eade1122 (2023).

52. Presman, D.M., Benitez, B., Lafuente, A.L. & Vazquez Lareu, A. Chromatin structure and dynamics: one nucleosome at a time. Histochem Cell Biol 162, 79–90 (2024).

53. Hibino, K. et al. Single-nucleosome imaging unveils that condensins and nucleosome-nucleosome interactions differentially constrain chromatin to organize mitotic chromosomes. Nat Commun 15, 7152 (2024).

54. Iida, S. et al. Single-nucleosome imaging reveals steady-state motion of interphase chromatin in living human cells. Science Advances. 8, eabn5626 (2022).

55. Manders, E.M., Kimura, H. & Cook, P.R. Direct imaging of DNA in living cells reveals the dynamics of chromosome formation. J Cell Biol 144, 813–821 (1999).

56. Nagashima, R. et al. Single nucleosome imaging reveals loose genome chromatin networks via active RNA polymerase II. J Cell Biol 218, 1511–1530 (2019).

57. Ryba, T. et al. Evolutionarily conserved replication timing profiles predict long-range chromatin interactions and distinguish closely related cell types. Genome Res 20, 761–770 (2010).

58. Vouzas, A.E. & Gilbert, D.M. Mammalian DNA Replication Timing. Cold Spring Harb Perspect Biol 13 (2021).

59. Nakayasu, H. & Berezney, R. Mapping replicational sites in the eucaryotic cell nucleus. J Cell Biol 108, 1–11 (1989).

60. O’Keefe, R.T., Henderson, S.C. & Spector, D.L. Dynamic organization of DNA replication in mammalian cell nuclei: spatially and temporally defined replication of chromosome-specific alpha-satellite DNA sequences. J Cell Biol 116, 1095–1110 (1992).

61. Dimitrova, D.S. & Berezney, R. The spatio-temporal organization of DNA replication sites is identical in primary, immortalized and transformed mammalian cells. J Cell Sci 115, 4037–4051 (2002).

62. Chagin, V.O. et al. 4D Visualization of replication foci in mammalian cells corresponding to individual replicons. Nat Commun 7, 11231 (2016).

63. Armstrong, C. & Spencer, S.L. Replication-dependent histone biosynthesis is coupled to cell-cycle commitment. Proc Natl Acad Sci U S A 118 (2021).

64. Mendiratta, S., Gatto, A. & Almouzni, G. Histone supply: Multitiered regulation ensures chromatin dynamics throughout the cell cycle. J Cell Biol 218, 39–54 (2019).

65. Ray-Gallet, D. et al. Dynamics of histone H3 deposition in vivo reveal a nucleosome gap-filling mechanism for H3.3 to maintain chromatin integrity. Mol Cell 44, 928–941 (2011).

66. Ahmad, K. & Henikoff, S. The histone variant H3.3 marks active chromatin by replication-independent nucleosome assembly. Mol Cell 9, 1191–1200 (2002).

67. Ran, F.A. et al. Genome engineering using the CRISPR-Cas9 system. Nat Protoc 8, 2281–2308 (2013).

68. Grimm, J.B. et al. A general method to improve fluorophores for live-cell and single-molecule microscopy. Nat Methods 12, 244–250, 243 p following 250 (2015).

69. Bodor, D.L. et al. The quantitative architecture of centromeric chromatin. eLife 3, e02137 (2014).

70. Buschbeck, M. & Hake, S.B. Variants of core histones and their roles in cell fate decisions, development and cancer. Nat Rev Mol Cell Biol 18, 299–314 (2017).

71. Clement, C. et al. High-resolution visualization of H3 variants during replication reveals their controlled recycling. Nat Commun 9, 3181 (2018).

72. Merrill, R.A. et al. A robust and economical pulse-chase protocol to measure the turnover of HaloTag fusion proteins. J Biol Chem 294, 16164–16171 (2019).

73. Gatto, A., Forest, A., Quivy, J.P. & Almouzni, G. HIRA-dependent boundaries between H3 variants shape early replication in mammals. Mol Cell 82, 1909–1923 e1905 (2022).

74. Consortium, E.P. An integrated encyclopedia of DNA elements in the human genome. Nature 489, 57-74 (2012).

75. Xiong, K. & Ma, J. Revealing Hi-C subcompartments by imputing inter-chromosomal chromatin interactions. Nat Commun 10, 5069 (2019).

76. Zhang, J. et al. An integrative ENCODE resource for cancer genomics. Nat Commun 11, 3696 (2020).

77. Zhao, P.A., Sasaki, T. & Gilbert, D.M. High-resolution Repli-Seq defines the temporal choreography of initiation, elongation and termination of replication in mammalian cells. Genome Biol 21, 76 (2020).

78. Van Rechem, C. et al. Collective regulation of chromatin modifications predicts replication timing during cell cycle. Cell Rep 37, 109799 (2021).

79. Lund, E.G., Duband-Goulet, I., Oldenburg, A., Buendia, B. & Collas, P. Distinct features of lamin A-interacting chromatin domains mapped by ChIP-sequencing from sonicated or micrococcal nuclease-digested chromatin. Nucleus 6, 30–39 (2015).

80. Tokunaga, M., Imamoto, N. & Sakata-Sogawa, K. Highly inclined thin illumination enables clear single-molecule imaging in cells. Nat Methods 5, 159–161 (2008).

81. Betzig, E. et al. Imaging intracellular fluorescent proteins at nanometer resolution. Science 313, 1642–1645 (2006).

82. Rust, M.J., Bates, M. & Zhuang, X. Sub-diffraction-limit imaging by stochastic optical reconstruction microscopy (STORM). Nat Methods 3, 793–795 (2006).

83. Selvin, P.R. et al. Fluorescence Imaging with One-Nanometer Accuracy (FIONA). CSH Protoc 2007, pdb top27 (2007).

84. Jaqaman, K. et al. Robust single-particle tracking in live-cell time-lapse sequences. Nat Methods 5, 695–702 (2008).

85. Dion, V. & Gasser, S.M. Chromatin movement in the maintenance of genome stability. Cell 152, 1355–1364 (2013).

86. van Steensel, B. & Belmont, A.S. Lamina-Associated Domains: Links with Chromosome Architecture, Heterochromatin, and Gene Repression. Cell 169, 780–791 (2017).

87. Peng, T. et al. Mapping nucleolus-associated chromatin interactions using nucleolus Hi-C reveals pattern of heterochromatin interactions. Nat Commun 14, 350 (2023).

88. Reznikoff, C.A., Brankow, D.W. & Heidelberger, C. Establishment and characterization of a cloned line of C3H mouse embryo cells sensitive to postconfluence inhibition of division. Cancer research 33, 3231–3238 (1973).

89. Otsuka, A. et al. Chromatin organization and behavior in HRAS-transformed mouse fibroblasts. Chromosoma (2024).

90. Maison, C., Quivy, J.P., Probst, A.V. & Almouzni, G. Heterochromatin at mouse pericentromeres: a model for de novo heterochromatin formation and duplication during replication. Cold Spring Harb Symp Quant Biol 75, 155–165 (2010).

91. Kubota, S. et al. Activation of the prereplication complex is blocked by mimosine through reactive oxygen species-activated ataxia telangiectasia mutated (ATM) protein without DNA damage. J Biol Chem 289, 5730–5746 (2014).

92. Jackson, D.A. & Pombo, A. Replicon clusters are stable units of chromosome structure: evidence that nuclear organization contributes to the efficient activation and propagation of S phase in human cells. J Cell Biol 140, 1285–1295 (1998).

93. Costa, A. & Diffley, J.F.X. The Initiation of Eukaryotic DNA Replication. Annu Rev Biochem 91, 107–131 (2022).

94. Polasek-Sedlackova, H., Miller, T.C.R., Krejci, J., Rask, M.B. & Lukas, J. Solving the MCM paradox by visualizing the scaffold of CMG helicase at active replisomes. Nat Commun 13, 6090 (2022).

95. Pollok, S., Bauerschmidt, C., Sanger, J., Nasheuer, H.P. & Grosse, F. Human Cdc45 is a proliferation-associated antigen. FEBS J 274, 3669–3684 (2007).

96. Bauerschmidt, C., Pollok, S., Kremmer, E., Nasheuer, H.P. & Grosse, F. Interactions of human Cdc45 with the Mcm2-7 complex, the GINS complex, and DNA polymerases delta and epsilon during S phase. Genes Cells 12, 745–758 (2007).

97. Gabriele, M. et al. Dynamics of CTCF- and cohesin-mediated chromatin looping revealed by live-cell imaging. Science 376, 496–501 (2022).

98. Collart, C., Allen, G.E., Bradshaw, C.R., Smith, J.C. & Zegerman, P. Titration of four replication factors is essential for the Xenopus laevis midblastula transition. Science 341, 893–896 (2013).

99. Tanaka, S., Nakato, R., Katou, Y., Shirahige, K. & Araki, H. Origin association of Sld3, Sld7, and Cdc45 proteins is a key step for determination of origin-firing timing. Curr Biol 21, 2055-2063 (2011).

100. Mantiero, D., Mackenzie, A., Donaldson, A. & Zegerman, P. Limiting replication initiation factors execute the temporal programme of origin firing in budding yeast. EMBO J 30, 4805–4814 (2011).

101. Wong, P.G. et al. Cdc45 limits replicon usage from a low density of preRCs in mammalian cells. PLoS One 6, e17533 (2011).

102. Metropolis, N., Rosenbluth, A.W., M.N., R., Teller, A.H. & Teller, E. Equation of State Calculations by Fast Computing Machines J Chem Phys 21, 1087 (1086 pages) (1953).

103. Morelli, M.J. & ten Wolde, P.R. Reaction Brownian dynamics and the effect of spatial fluctuations on the gain of a push-pull network. J Chem Phys 129, 054112 (2008).

104. Hihara, S. et al. Local nucleosome dynamics facilitate chromatin accessibility in living mammalian cells. Cell Rep 2, 1645–1656 (2012).

105. Imai, R. et al. Density imaging of heterochromatin in live cells using orientation-independent-DIC microscopy. Mol Biol Cell 28, 3349–3359 (2017).

106. Maeshima, K. et al. The physical size of transcription factors is key to transcriptional regulation in the chromatin domains. Journal of Physics: condensed matters 27, 064116 (064110 pp) (2015).

107. MacNeill, S. The eukaryotic replisome: A guide to protein structure and function. (Springer, 2012).

108. Volpi, I., Gillespie, P.J., Chadha, G.S. & Blow, J.J. The role of DDK and Treslin-MTBP in coordinating replication licensing and pre-initiation complex formation. Open Biol 11, 210121 (2021).

109. Cvetkovic, M.A. et al. The structural mechanism of dimeric DONSON in replicative helicase activation. Mol Cell 83, 4017–4031 e4019 (2023).

110. Cornacchia, D. et al. Mouse Rif1 is a key regulator of the replication-timing programme in mammalian cells. EMBO J 31, 3678–3690 (2012).

111. Yamazaki, S. et al. Rif1 regulates the replication timing domains on the human genome. EMBO J 31, 3667–3677 (2012).

112. Klein, K.N. et al. Replication timing maintains the global epigenetic state in human cells. Science 372, 371–378 (2021).

113. Filion, G.J. et al. Systematic protein location mapping reveals five principal chromatin types in Drosophila cells. Cell 143, 212–224 (2010).

114. Natsume, T. et al. Acute inactivation of the replicative helicase in human cells triggers MCM8-9-dependent DNA synthesis. Genes Dev 31, 816–829 (2017).

115. Wang, W. et al. Genome-wide mapping of human DNA replication by optical replication mapping supports a stochastic model of eukaryotic replication. Mol Cell 81, 2975–2988 e2976 (2021).

116. Massey, D.J. & Koren, A. High-throughput analysis of single human cells reveals the complex nature of DNA replication timing control. Nat Commun 13, 2402 (2022).

117. Gnan, S. et al. Kronos scRT: a uniform framework for single-cell replication timing analysis. Nat Commun 13, 2329 (2022).

118. Rhind, N. DNA replication timing: random thoughts about origin firing. Nat Cell Biol 8, 1313–1316 (2006).

119. Bechhoefer, J. & Rhind, N. Replication timing and its emergence from stochastic processes. Trends in genetics: TIG 28, 374–381 (2012).

120. Gindin, Y., Valenzuela, M.S., Aladjem, M.I., Meltzer, P.S. & Bilke, S. A chromatin structure-based model accurately predicts DNA replication timing in human cells. Mol Syst Biol 10, 722 (2014).

121. Casas-Delucchi, C.S. et al. Histone hypoacetylation is required to maintain late replication timing of constitutive heterochromatin. Nucleic Acids Res 40, 159–169 (2012).

122. Foti, R. et al. Nuclear Architecture Organized by Rif1 Underpins the Replication-Timing Program. Mol Cell 61, 260–273 (2016).

123. Gnan, S. et al. Nuclear organisation and replication timing are coupled through RIF1-PP1 interaction. Nat Commun 12, 2910 (2021).

124. Sima, J. et al. Identifying cis Elements for Spatiotemporal Control of Mammalian DNA Replication. Cell 176, 816–830 e818 (2019).

125. Brustel, J. et al. Histone H4K20 tri-methylation at late-firing origins ensures timely heterochromatin replication. EMBO J 36, 2726–2741 (2017).

126. Heinz, K.S. et al. Peripheral re-localization of constitutive heterochromatin advances its replication timing and impairs maintenance of silencing marks. Nucleic Acids Res 46, 6112–6128 (2018).

127. Schwaiger, M., Kohler, H., Oakeley, E.J., Stadler, M.B. & Schubeler, D. Heterochromatin protein 1 (HP1) modulates replication timing of the Drosophila genome. Genome Res 20, 771–780 (2010).

128. Dave, A., Cooley, C., Garg, M. & Bianchi, A. Protein phosphatase 1 recruitment by Rif1 regulates DNA replication origin firing by counteracting DDK activity. Cell Rep 7, 53–61 (2014).

129. Hiraga, S. et al. Rif1 controls DNA replication by directing Protein Phosphatase 1 to reverse Cdc7-mediated phosphorylation of the MCM complex. Genes Dev 28, 372–383 (2014).

130. Mattarocci, S. et al. Rif1 controls DNA replication timing in yeast through the PP1 phosphatase Glc7. Cell Rep 7, 62–69 (2014).

131. Manley, S. et al. High-density mapping of single-molecule trajectories with photoactivated localization microscopy. Nat Methods 5, 155–157 (2008).

132. Hoffman, D.P. et al. Correlative three-dimensional super-resolution and block-face electron microscopy of whole vitreously frozen cells. Science 367 (2020).

133. Berger, C. et al. Cryo-electron tomography on focused ion beam lamellae transforms structural cell biology. Nat Methods 20, 499–511 (2023).

134. Iida, S. et al. Orientation-independent-DIC imaging reveals that a transient rise in depletion attraction contributes to mitotic chromosome condensation. Proc Natl Acad Sci U S A 121, e2403153121 (2024).

135. Ide, S., Imai, R., Ochi, H. & Maeshima, K. Transcriptional suppression of ribosomal DNA with phase separation. Sci Adv 6 (2020).

136. Labun, K. et al. CHOPCHOP v3: expanding the CRISPR web toolbox beyond genome editing. Nucleic Acids Res 47, W171–W174 (2019).

137. Maeshima, K. et al. Cell-cycle-dependent dynamics of nuclear pores: pore-free islands and lamins. J Cell Sci 119, 4442–4451 (2006).

138. Saito, Y., Kobayashi, J., Kanemaki, M.T. & Komatsu, K. RIF1 controls replication initiation and homologous recombination repair in a radiation dose-dependent manner. J Cell Sci 133 (2020).

139. Lewis, C.D. & Laemmli, U.K. Higher order metaphase chromosome structure: evidence for metalloprotein interactions. Cell 29, 171–181 (1982).

140. Shimamoto, Y., Tamura, S., Masumoto, H. & Maeshima, K. Nucleosome-nucleosome interactions via histone tails and linker DNA regulate nuclear rigidity. Mol Biol Cell 28, 1580–1589 (2017).

141. Ura, K. & Kaneda, Y. Reconstitution of chromatin in vitro. Methods Mol Biol 181, 309–325 (2001).

142. Ide, S. et al. Telomere-specific chromatin capture using a pyrrole-imidazole polyamide probe for the identification of proteins and non-coding RNAs. Epigenetics Chromatin 14, 46 (2021).

143. Laemmli, U.K. Cleavage of structural proteins during the assembly of the head of bacteriophage T4. Nature 227, 680–685 (1970).

144. Ewels, P.A. et al. The nf-core framework for community-curated bioinformatics pipelines. Nat Biotechnol 38, 276–278 (2020).

145. Ewels, P.A., et al. The nf-core framework for community-curated bioinformatics pipelines. Zenodo. (2022).

146. Zhang, Y. et al. Model-based analysis of ChIP-Seq (MACS). Genome Biol 9, R137 (2008).

147. Davis, C.A. et al. The Encyclopedia of DNA elements (ENCODE): data portal update. Nucleic Acids Res 46, D794–D801 (2018).

148. Schindelin, J., et al. Fiji: an open-source platform for biological-image analysis. Nat Methods 9, 676-682 (2012).

149. Schneider, C.A., Rasband, W.S. & Eliceiri, K.W. NIH Image to ImageJ: 25 years of image analysis. Nat Methods 9, 671–675 (2012).

150. Fox, M.H. A model for the computer analysis of synchronous DNA distributions obtained by flow cytometry. Cytometry 1, 71–77 (1980).

151. van der Walt, S. et al. scikit-image: image processing in Python. PeerJ 2, e453 (2014).

152. Grimm, J.B. et al. A General Method to Improve Fluorophores Using Deuterated Auxochromes. JACS Au 1, 690–696 (2021).

